# The Rise and Fall of SARM1 Base-Exchange Inhibitors

**DOI:** 10.1101/2025.11.11.687870

**Authors:** Thomas Lundbäck, Vijay Chandrasekar, Chendi Gu, Hyoungseok Ju, Robyn McAdam, Maria Palomero, Kasim Sader, Bradley Peter, Lisa Wissler, Philip Nevin, Edmund Foster, Tanguy Jamier, Pravallika Manjappa, Carina Johansson, Jenny Sandmark, Mei Ding, Anette Persson-Kry, Sanhita Mitra, Tugce Munise Satir, Bilada Bilican, Mirko Messa, Graham Fraser, John Linley, Helen Plant, Rachel Moore, Tina Seifert, Michael Lerche, Carina Raynochek, Ewa Nilsson, Nour Majbour, Richard Lucey, Taiana Maia de Oliveira, Qi Wang, Iain Chessell, Perla Breccia, Rebecca Jarvis

## Abstract

The sterile alpha and TIR motif containing 1 (SARM1) enzyme is a key driver of axonal degeneration in response to injury, making it an attractive target for treating chemotherapy-induced peripheral neuropathy (CIPN) and other nervous system diseases. In this study, we identified and optimised a new class of base-exchange inhibitors (BEXi) targeting SARM1 and explored their molecular interactions and conformational effects using cryo-EM, HDX-MS and SAXS. Although BEXi produced robust inhibition across all biochemical and cellular assay formats, application at sub-inhibitory concentrations consistently led to paradoxical SARM1 activation, and in neuronal assays, accelerated neurite degeneration. Further analysis showed that BEXi only delayed, rather than prevented, neurite degeneration when applied to primary neuronal cells, even at exceedingly high inhibitor concentrations. These results prompted us to discontinue BEXi development in favour of alternative strategies, underscoring the complexity of SARM1 as a therapeutic target and the need for comprehensive, mechanistically informed screening cascades.

## Introduction

While axonal degeneration occurs naturally with aging, it is also an early hallmark of several neurodegenerative diseases, including chemotherapy-induced peripheral neuropathy (CIPN), amyotrophic lateral sclerosis (ALS), and multiple sclerosis (MS)^1–3^. Dating back to the 19th century, Augustus Waller^4^ first described the process of distal nerve fibre degeneration following transection. Subsequent research has progressively unveiled the underlying molecular mechanisms driving this degenerative process known as Wallerian degeneration^5–9^. Nerve fibre damage leads to severe complications in affected patients, causing sensory and motor disabilities such as loss of sensation, pain, functional limitations, and, in severe cases, an increased risk of falling. The emergence of neuropathy is a common consequence of chemotherapy treatment^10–13^, where peripheral nerves are damaged by the chemotherapeutic agents. While CIPN can be transient, subsets of patients experience chronic axonopathy^10,14^. In-depth studies of causative and protective mechanisms, as well as strategies to prevent nerve fibre damage through treatments, are imperative for enhancing the quality of life for affected patients.

An understanding of the NAD^+^ salvage pathway’s central role in Wallerian degeneration emerged from studies of a spontaneous mutation in the slow Wallerian degeneration mouse (Wld^S^)^15^, where disintegration of damaged axons is notably delayed. A key feature of Wld^S^ is the expression of a non-natural fusion protein that includes nicotinamide mononucleotide adenylyltransferase 1 (NMNAT1)^16–18^, an enzyme involved in converting nicotinamide mononucleotide (NMN) to nicotinamide adenine dinucleotide (NAD^+^). While wild-type NMNAT1 is primarily located in the nucleus, the fusion protein presents also in the axons of Wld^S^ mice^19,20^. Two additional NMNAT isoforms exist, with normal axonal maintenance of NAD^+^ homeostasis relying on axonal transport of NMNAT2 from the cytoplasm^21^. Upon neuronal damage, NMNAT2 transport to the distal axon halts, leading to a rapid decline in axonal concentration, a consequent loss of NAD^+^, and subsequent energy crisis^22^. This finding explains the protective role of the Wld^S^ fusion protein, whose enzymatic activity sustains NAD^+^ levels^23,24^, a conclusion supported by the protective effects of exogenous NAD^+^ precursors^25^. Further research into prodegenerative genes identified sterile alpha and TIR motif-containing 1 (SARM1), which amplifies the impact of NMNAT2 loss by rapidly depleting remaining NAD^+^levels^26,27^. Recognised as a key orchestrator of axon degeneration across multiple neurodegenerative diseases, SARM1 inhibitors are now of significant therapeutic interest^8,28,29^.

SARM1 is a multi-domain enzyme that presents as an auto-inhibited assembly in healthy neurons, with expression also found in the gastrointestinal tract, pancreas, and some immune cells^6,30^. The octamer organises structurally to an inner ring and outer ring, with the inner ring formed by tandem sterile alpha motif (SAM) domains, and the outer ring made up of interweaving armadillo repeat domains (ARM) and catalytic toll-interleukin-1 receptor (TIR) homology domains from the eight protomers^31,32^. Through this arrangement the TIR domains remain wedged in an inactive state until allosteric activation is triggered at the ARM domain following damage to the axons. This activation is mechanistically driven by changes to the ratio of NMN and NAD^+^, which bind to the same site in the ARM domain, but functioning inversely as the allosteric activator and inhibitor, respectively^33–39^. Activation of SARM1 transitions this enzyme from an auto-inhibited state to an active NAD^+^ hydrolase, rapidly degrading NAD^+^ and sequentially generating nicotinamide (NAM) and adenosine diphosphate ribose (ADPR) or cyclic ADPR (cADPR)^40^. Besides hydrolysis to ADPR and cyclisation to cADPR, SARM1 can also engage in base-exchange reactions, where the stabilised oxonium ion transition state in the active site can react with matching compounds carrying a basic nitrogen – as exemplified by the formation of nicotinic acid adenine dinucleotide (NaADP^+^)^41,42^. In addition to NAD^+^, SARM1 can process NADP^+42,43^, and ongoing research aims to determine the relative importance of its substrates and products in driving axon degeneration^29,44,45^.

The SARM1 role in axonal degeneration is corroborated by multiple studies in knockout mice, where genetic deletion of SARM1 affords protection across models of chemotherapy-induced and mechanical nerve damage^26,46^ (see *e.g.* Loring & Thompson, 2020 for an overview^47^). Protection can be reproduced pharmacologically using antisense oligonucleotides^48–50^, adenovirus-mediated expression of a dominant negative mutant form of SARM1^51^, and small molecule inhibitors^41,52–63^, in the latter cases with protection demonstrated both *in vitro* and *in vivo*. Hitherto identified small molecule inhibitor classes include compounds that covalently engage cysteine 311 to allosterically regulate SARM1 activity as well as uncompetitive inhibitors formed in the active site of SARM1 through a base-exchange reaction. Parent compounds can be relatively small, as exemplified by 5-Iodoisoquinoline (DSRM-3716)^53^, which translates to excellent ligand efficiency. Two base-exchange inhibitors of SARM1 are currently being evaluated in clinical phase 1 studies by Lilly and NuraBio, respectively (LY3873862 – NCT05492201 and NB-4746). While interest in SARM1 inhibition remains strong, recent publications have also highlighted a potential challenge with these types of inhibitors^60–66^, as engagement of SARM1 at low target occupancy was shown to activate SARM1 and accelerate axon degeneration both *in vitro* and *in vivo*. Further investigations are needed to better understand the physiological relevance of these findings.

In this study, we describe our efforts to identify and optimise new SARM1 inhibitors through high-throughput screening (HTS) and medicinal chemistry to define structure-activity relationships (SAR) for a new class of SARM1 inhibitors. Using a comprehensive suite of biochemical, biophysical, and cellular *in vitro* assays, we optimised the potencies of these inhibitors and thoroughly characterised their mechanisms of action, including molecular-level analysis using cryo-electron microscopy (cryo-EM). Our findings demonstrated robust inhibition across all assay settings, with consistent results observed between human and rodent species. However, in line with recent studies, we also observed that sub-inhibitory concentrations of BEXi result in SARM1 activation and accelerated neurite degeneration. Consequently, we have decided to halt further development of these compound series, underscoring the need for alternative therapeutic strategies to target SARM1 effectively.

## Results

### Inhibitor discovery and characterisation

At the inception of our HTS campaign the regulatory roles of NMN and NAD^+^ as allosteric activators and inhibitors of SARM1 activity^35^ had not been fully elucidated. To boost enzymatic activity of human recombinant truncated SARM1^Δ1–27^ protein, our biochemical assay development included testing of multiple assay buffer conditions. This led us to include n-Dodecyl-β-D-maltoside (DDM) in the assay buffer, which was compatible with the application of acoustic mist-ionisation mass spectrometry (AMI-MS)^67,68^ to measure NAD^+^ turnover to ADPR in 384-well plates. This approach gave an assay with suitable signal to background ratio and linearity in response with respect to enzyme concentration and time (see Supplementary Fig. 1 for details). Armed with this assay we interrogated the AstraZeneca HTS deck of 1.8 million compounds and identified new inhibitors of SARM1, examples of which are shown in Fig. 1A.

**Figure 1.**
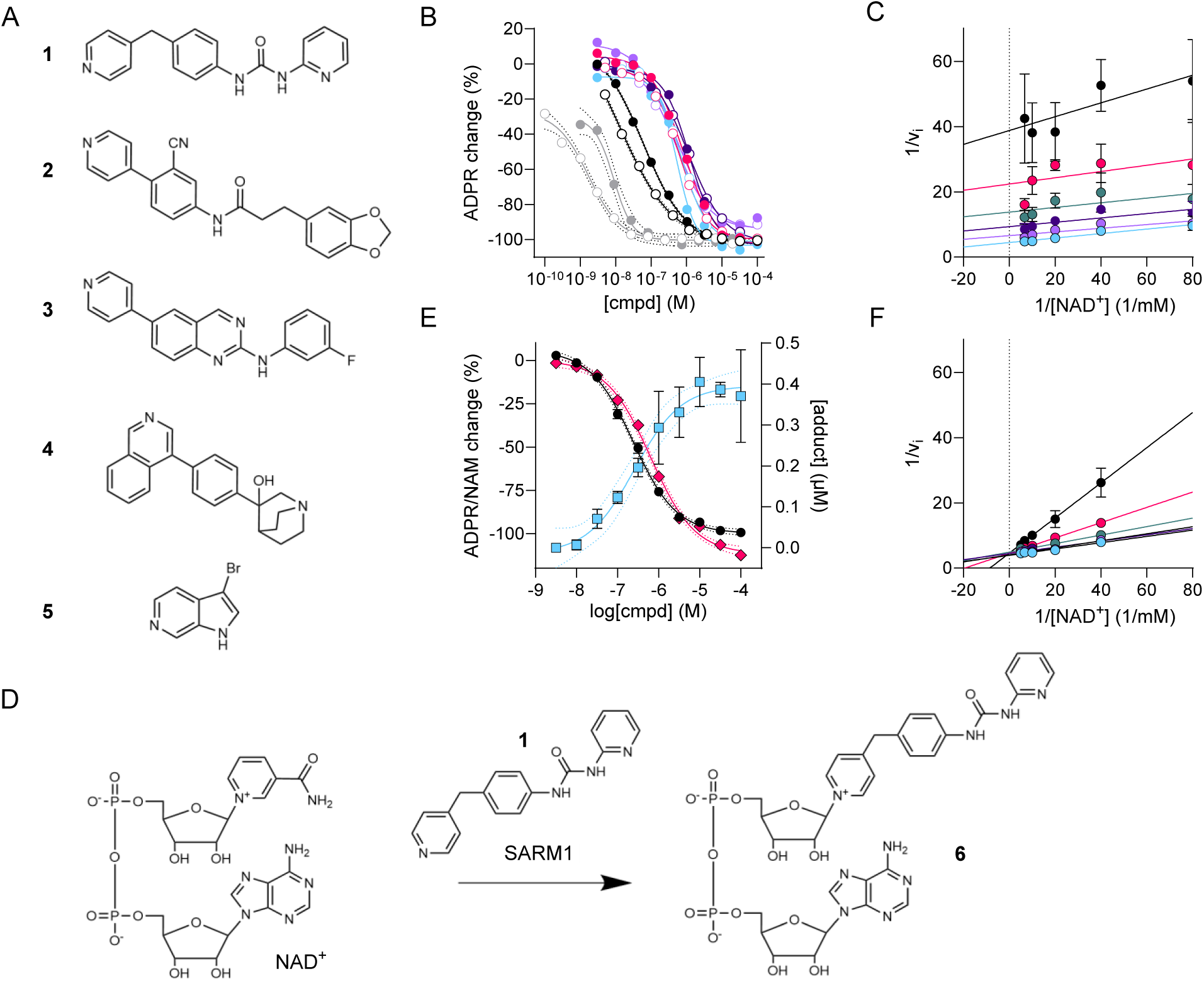
| Hit identification and characterisation. A) Chemical structures of hit compounds **1**-**5** identified in this study. B) Inhibition of SARM1 hydrolysis of NAD^+^ to ADPR with data presented as averages from multiple independent test occasions (N) in two complementary biochemical assays based on SARM1^Δ1–27^ (closed symbols) and SARM1^FL^ (open symbols), respectively. Symbols denote compound **1** (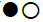; N=232; 22), **2** (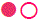; N=2; 21), **3** (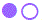; N=6; 1), **4** (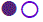; N=2; 3), **5** (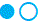; N=2; 1), and **6** (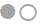; N=2; 6). The solid lines represent the best fits to a four-parameter logistic equation in GraphPad Prism. An equivalent plot including error bars is available in Supplementary Fig. 2D. C) Inhibitor **1** demonstrates uncompetitive inhibition of SARM1^Δ1–27^ with respect to NAD^+^. Each symbol and line in the double reciprocal Lineweaver-Burk plot represent a different inhibitor concentration with threefold dilutions from 0.41 µM (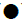) down to 5.1 nM (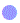) and a DMSO control (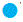). D) Outline of the expected base-exchange reaction in the active site of SARM1, whereby the parent compound **1** replaces nicotinamide to form a new dinucleotide species **6**. E) The concentration dependent inhibition of SARM1^Δ1–27^ enzymatic activity by inhibitor **1** is similarly reflected measuring release of NAM (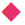) and formation of ADPR (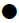), respectively. This inhibition coincided with the appearance of **6** (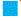), which is generated by SARM1^Δ1–27^ in the presence of **1** and NAD^+^. F) The adduct **6** shows competitive inhibition of SARM1^Δ1–27^ activity with respect to the NAD^+^ substrate. See the 1C legend for info on inhibitor concentrations and symbols.

We arrived at this set of compounds following multiple counter screens to remove artefacts as detailed in Supplementary Figs. 2A&B alongside screening statistics. Follow-up characterisation included concentration response testing in the same screening assay and a complementary biochemical assay to ensure compounds were active towards full-length human SARM1, *i.e.* including the first 27 N-terminal amino acids. This was achieved based on lysates from HEK293 cells over-expressing SARM1^FL^, demonstrating equivalent data within experimental uncertainty (Supplementary Fig. 2C). Concentration response curves are shown in Fig. 1B, with best-fit inhibitory potencies summarised in Table 1. These hits also showed activity in a cell-based assay performed in HEK293 over-expressing a constitutively active form of human SARM1 lacking the N-terminal mitochondrial localisation and ARM domains (SARM1^SST^)^27^. Cell activity against the truncated enzyme eliminates the ARM domain as a potential site of action for these compounds. To evaluate activity in a disease relevant neuronal model expressing endogenous SARM1, we used a rat primary dorsal root ganglion (DRG) cell model, where multiple hits showed protection against vincristine induced axon degeneration at sub-µM level. Additional details on these assays are provided in a later paragraph describing structure-activity relationships for **1**.

**Table 1.** | Example Screening Hits.

| Compound | SARM1<sup>Δ1-27</sup><br>biochemical IC<sub>50</sub><br>μM | SARM1<sup>SST</sup> cell*<br>IC<sub>50</sub> μM | Neurite protection<br>rat DRG IC<sub>50</sub> μM | LogD<br>pH 7.4 | Aq. Solubility<br>pH 7.4 μM |
| --- | --- | --- | --- | --- | --- |
| **1** | 0.080 | 0.15 | 0.30 | 3.6 | 6.3 |
| **2** | 0.77 | 2.2 | >1.0x10<sup>2</sup> | 3.5 | <1.6 |
| **3** | 0.50 | 0.37 | >1.0x10<sup>2</sup> | 4.3 | <1.1 |
| **4** | 1.1 | 14 | ND | 1.6 | 0.99x10<sup>3</sup> |
| **5** | 0.52 | 0.52 | 0.92 | 2.6 | >1.0x10<sup>3</sup> |

\* The SARM<sup>SST</sup> numbers refer to the cell assay using NAD-Glo™ as readout.
LogD and aqueous solubility were measured as previously described for our DMPK Wave 1 panel of assays<sup>69</sup>.

It became apparent during hit annotation, which involved resynthesis as well as nearest neighbour expansion, that none of the hit series confirmed binding to the recombinant human SARM1 protein in biophysical assays (data not shown). The lack of binding was initially concerning, so we engaged in additional enzyme kinetics and structural studies to elucidate their mechanism of action (MoA). Firstly, when NMN was corroborated in the literature as an endogenous allosteric activator of SARM1^34–36^, the identified compounds were tested to understand if they retained the same inhibitory potencies when replacing DDM with NMN – which was the case (Supplementary Fig. 2E). Finally, potential promiscuous non-specific mechanisms, such as PAINS and colloidal compound aggregation^70^, were probed using jump dilution experiments, which confirmed reversible inhibition (Supplementary Fig. 2F).

We next investigated the substrate concentration dependence of inhibition by applying the biochemical assay at 20-fold higher NAD^+^ concentration, *i.e.* 0.5 mM versus 25 µM (Supplementary Fig. 2G). Observed potencies are similar under both conditions, suggesting a non-competitive or uncompetitive inhibition mechanism. To distinguish between these mechanisms, we evaluated a broader span of substrate concentrations as illustrated for inhibitor **1** (Fig. 1C). Data confirmed uncompetitive inhibition, explaining why binding to the protein were not detectable in the absence of substrate. At this time there had not been any disclosures of mechanism-based inhibitors of SARM1. Base-exchange (BEX) was a known feature of the related enzyme CD38^71–73^. Taken together with the observation of an aza nitrogen as a common structural feature amongst the hit compounds (Fig. 1A), we suspected this could be at play. The expected reaction scheme in the active site of SARM1 is depicted in Fig. 1D.

### Chemical and Enzymatic synthesis of adduct 6

To verify base exchange with compound **1**, adduct **6** (Fig. 1D) was chemically synthesised through a multistep sequence and isolated using preparatory HPLC (see Supplemental Methods). With **6** in hand as a reference, we confirmed SARM1-catalysed base-exchange in the presence of NAD^+^ by LC-MS (Fig. 1E). The availability of purified **6** also allowed enzymatic studies (Fig. 1F), confirming significantly improved potency (Table 1) and competitive inhibition with respect to the NAD^+^ substrate. We observed similar SARM1 inhibition when basing measurements on either nicotinamide or ADPR release for **1** (Fig. 1E), as expected for a potent BEX inhibitor that is retained in the active site of SARM1. A different behaviour is anticipated for a base-exchange substrate that renders a product that readily leaves the active site, as exemplified for **Probe 1** that we identified in our hit annotation efforts (Supplementary Figs. 2H&I). Its application in a cell target engagement assay in human iPSC derived neuronal cells is described in a later paragraph.

To facilitate the preparation of additional adducts, and avoid lengthy chemical synthesis, an enzyme catalysed procedure was developed. In contrast to the procedure reported by Shi *et al.*^57^ adducts were generated from the parent small molecule inhibitors and NAD^+^ using SARM1^Δ1–27^ immobilised on beads, which allowed for recycling of the enzyme and significantly reduced the amount of protein required (Supplementary Methods). This method enabled the production of multiple adducts with conversions of 35-98% (LC-MS) from the parent molecule which were used in subsequent biophysical and structural studies.

### Biophysical and structural studies

To study the impact of BEX inhibitors on SARM1 conformation, and to compare it with changes induced by physiologically relevant allosteric regulators, we performed hydrogen-deuterium exchange mass spectrometry (HDX-MS) experiments. Samples of recombinant human SARM1^Δ1–27^ were treated with NAD^+^, NMN, adduct **6** or control buffer, mixed with a deuterium-based buffer and incubated as a function of time at 20°C. Quenched and digested samples were analysed by electrospray ionization mass spectrometry (ESI-MS) and observed changes in the deuterium uptake were mapped onto a published cryo-EM structure of SARM1 (PDB ID 7CM6) to illustrate protected and exposed areas (Fig. 2A). Incubation with NAD^+^ showed increased protection in the allosteric NAD^+^-binding site in the ARM domain as expected (Fig. 2A, Supplementary Fig. 3A, and Supplementary Table 2), while the interface between the TIR domain and neighbouring ARM domain showed modest protection, suggesting a more closely packed conformation of the SARM1 octamer. This is broadly in line with previous HDX data on SARM1^74^, with the TIR domain being locked in an inactive conformation in the auto-inhibited state of the protein at high NAD^+^ concentrations^34–36^.

**Figure 2.**
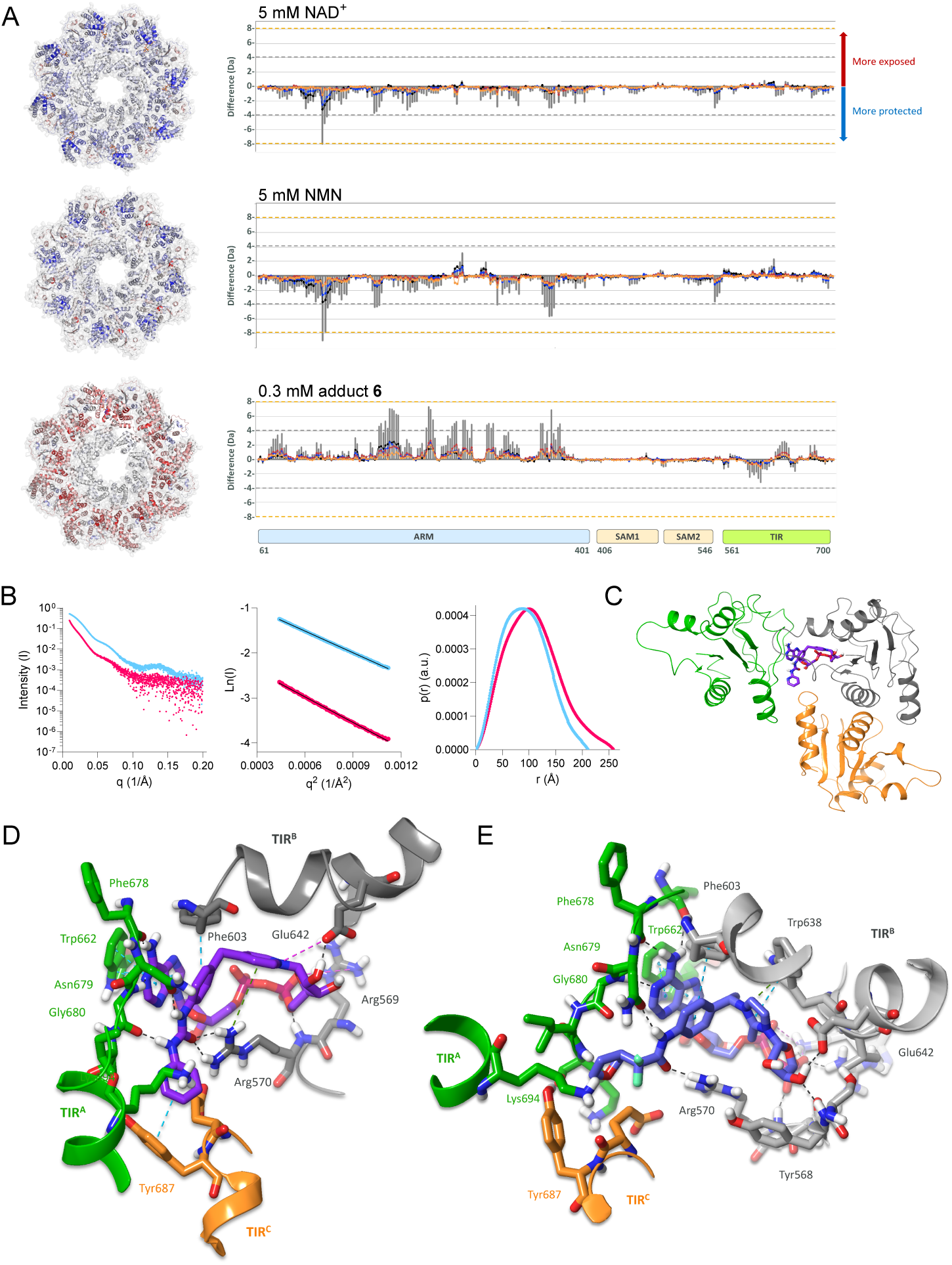
| Structural and biophysical characterisation of BEXi binding. A) HDX-MS studies in the presence of allosteric inhibitor NAD^+^ (5 mM, top panel), allosteric activator NMN (5 mM, middle panel) and BEXi adduct **6** (0.3 mM, bottom panel). Results are shown as butterfly plots on the right: the y-axis indicates the difference in deuterium uptake (Da) in absence *vs* presence of ligand; the x-axis denotes SARM1 peptides arranged in order of increasing residue number from the N-terminus; vertical grey bars represent the total difference of each peptide summed over all time points; coloured lines show uptake difference from 0.5 to 30 min labelling time, and on the left: mapped to a cryo-EM structure of the SARM1 octamer (7CM6): peptides with no mass difference are grey; peptides with increased protection in the presence of ligand are blue, and peptides that display increased exposure in the presence of ligand are red. The colour intensity indicates the degree of deuterium labelling. B) Left: Experimental solution scattering profiles for SARM1 in its apo and NMN bound states (blue dots) versus the conformation of the adduct **6** liganded state (red dots). Apo and NMN bound states are isostructural. Middle and right: Guinier fit and p(r) function of NMN (blue dots) and adduct **6** (red dots) bound states. Fits to the data are shown as solid lines with R_g_ = 71.3±1.1 Å and 82.8±1.2 Å, respectively. C) Cryo-EM structure of adduct **6** (purple) in complex with SARM1^TIR^. The assembly of the TIR domains form the orthosteric pocket at the interface of three distinct TIR monomers (shown in green, grey, and orange). D) Detail of interactions of adduct **6** with the three distinct TIR domains. Hydrogen bonds are shown as black dashed lines, π-π interactions as cyan dashed lines and electrostatic interactions as magenta dashed lines. E) Corresponding interactions of adduct **17** (blue).

Replacing the allosteric inhibitor NAD^+^ with the activator NMN resulted in similar protection of the same allosteric binding site in the ARM domain (Fig. 2A, Supplementary Fig. 3B, and Supplementary Table 2), in agreement with current literature that these metabolites compete for the same site. However, in contrast to NAD^+^, incubation with NMN additionally resulted in exposure of helices 13 (aa 255-260) and 15 (aa 282-286) in the ARM domain. As these helices are located at the interface of the ARM and TIR domains, this observation suggests that NMN binding at the ARM domain triggers a release and activation of the TIR domain. Interestingly, we did not observe a corresponding exposure of residues at the TIR domain interface. We speculate that these residues are instead involved in the dimerization of TIR domains from two protomers within the activated SARM1 octamer, as observed in crystal structures of TIR domains (PDB ID 6O0Q^32^, thereby masking the helix from deuterium exchange.

While the HDX data primarily showed local changes to the SARM1 conformation in the presence of either NAD^+^ and NMN, incubation with adduct **6** showed more dramatic effects with the entire ARM domain becoming exposed (Figure 2A, Supplementary Fig. 3C and Supplementary Table 3). The active site in the TIR domain, however, showed increased protection, in agreement with binding of adduct **6** to the active site. No protection was observed in the allosteric NAD^+^/NMN binding site in the ARM domain, indicating no secondary binding of adduct **6** to the allosteric site under these conditions. To further characterise the conformation of adduct **6** stabilised SARM1, we performed small angle X-ray scattering (SAXS) coupled to inline size exclusion chromatography (SEC-SAXS). While NMN and apo states showed no significant difference in scattering pattern, in the presence of adduct **6** SARM1 displayed a different scattering profile and increased the invariant parameters R_g_ and D_max_ (Fig. 2B), indicating that the SARM1 octamer undergoes significant expansion when binding adduct **6.** The HDX-MS and SAXS studies collectively indicate that the inhibited conformation induced by adduct **6** assumes a significantly more open form that is distinctly different from the NAD^+^-induced auto-inhibited conformation, in agreement with cryo-EM structures of other SARM1-BEXi adduct complexes^57,61^.

We next engaged in cryo-EM studies of SARM1 in the presence of the BEX adduct **6**. Recombinant human TIR domains were shown to self-associate into two-stranded filaments, a feature of the SARM1 activated state, confirming prior literature findings^55,57^ (Supplementary Figs. 4A-C and Supplementary Table 4). The orthosteric binding site occupies the interface of three TIR domains: TIR^A^, TIR^B^, and TIR^C^ (Figs. 2C&D). The ADPR segment of the adduct mirrors the interactions of NAD^+^, including adenine ring interactions with Phe678^B^ and Gly680^A^, π-stacking with Trp662^A^ and a salt bridge with the catalytic Glu642 in TIR^B^ (see Supplementary Fig. 4H for an interaction map). The phenyl ring forms a π-edge interaction with Phe603^A^. The urea NH groups are stabilized by hydrogen bonds to Asn679^A^ and Gly680^A^, and the carbonyl oxygen binds to the flexible side chain of Arg570^A^. The terminal pyridine fits between chains TIR^A^ and TIR^C^, interacting with Lys694^A^ and Tyr687^C^, and maintaining the binding pocket’s integrity. Taken together these results support a scenario where conversion of **1** to **6** results in the dramatic conformational change that is associated with SARM1 activation, resulting in oligomerisation of the **6**-associated TIR domains, in line with a recent disclosure for another BEXi^75^.

### SAR-driving *in vitro* cell assays

To understand cell potency of identified BEXi we employed a HEK293 cell assay with transient overexpression of a constitutively active form of human SARM1 containing SAM-SAM-TIR domains without the N-terminal ARM domain (SARM1^SST^). This overexpression drives depletion of intracellular NAD^+^ concentrations, the rescue of which was monitored using NAD-Glo™ (exemplified for compound **1** in Fig. 3A). SARM1-driven depletion of NAD^+^ leads to a concomitant loss of cell viability, which was rescued by SARM1 BEXi as measured using CellTiter-Glo® (Fig. 3A). We profiled in-house inhibitors alongside published inhibitors in these assays and observed correlation with the biochemical assays (Fig. 3B), with half a log unit potency drop-off for the NAD-Glo™ assay. This correlation held true also for the cell viability readout (Supplementary Fig. 5A), albeit without the potency drop-off, *i.e.* protection is observed at lower concentrations for the cell viability readout (Supplementary Fig. 5B – see also Fig. 3A where the concentration response is left-shifted in the viability assay).

**Figure 3.**
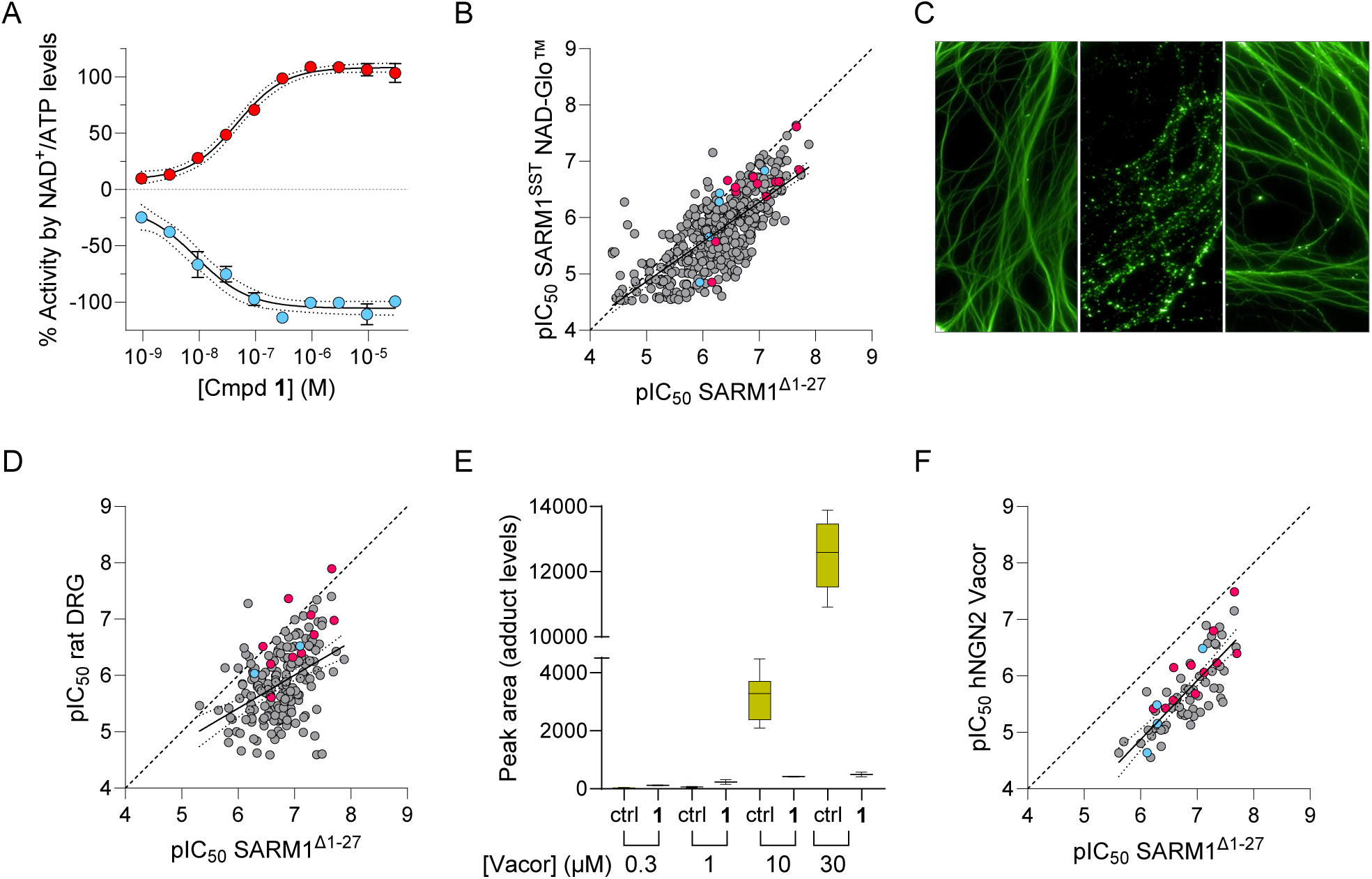
| SARM1 *in vitro* SAR driving cell assays. A) Inhibition profiles for **1** in HEK293 cells transiently transfected with constitutively active SARM1^SST^. The inhibition of SARM1 measured using rescue of intracellular NAD^+^ levels by NAD-Glo™ (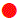; top panel) correlates with an improvement of cell viability (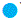; bottom panel). B) The measured potencies in the HEK293 SARM1^SST^ cell assay correlates with measured potencies in the biochemical assay across BEXi chemotypes (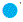, Table 1 hits; 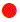, Table 2 compounds including competitor compounds; 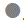, Other BEXi), albeit with half a log unit drop-off. The solid line represents the best-fit straight line with a slope of 0.70, a correlation coefficient r^2^=0.54 and with the 95% confidence bands shown by dotted lines. The dashed line represents equality. C) Rat embryonic DRG neurons cultured for 7 days and then treated (from left to right panel) with vehicle, 40 nM vincristine, or 40 nM vincristine and 10 µM Compound **14** (see Table 2 further down the main text), respectively. At completion of the assay cells were stained for βIII tubulin and imaged. D) Correlation of observed potencies in the rat DRG cell assay (y-axis) and the biochemical assay based on human recombinant SARM1^Δ1–27^ (x-axis). See the 3B legend for info on lines and symbols. The slope of the solid line is 0.59, with a correlation coefficient r^2^=0.18. E) Data for the human iPSC NGN2 cell target engagement assay as a function of Vacor concentration, with measured peak areas for base-exchange adduct formation in the absence and presence of 10 µM **1**. F) Correlation of measured potencies in the hNGN2 target engagement assay with measured potencies in the biochemical assay. Lines and symbols are as denoted in 3B with a slope of and 1.04 and r^2^=0.63.

**Table 2.**
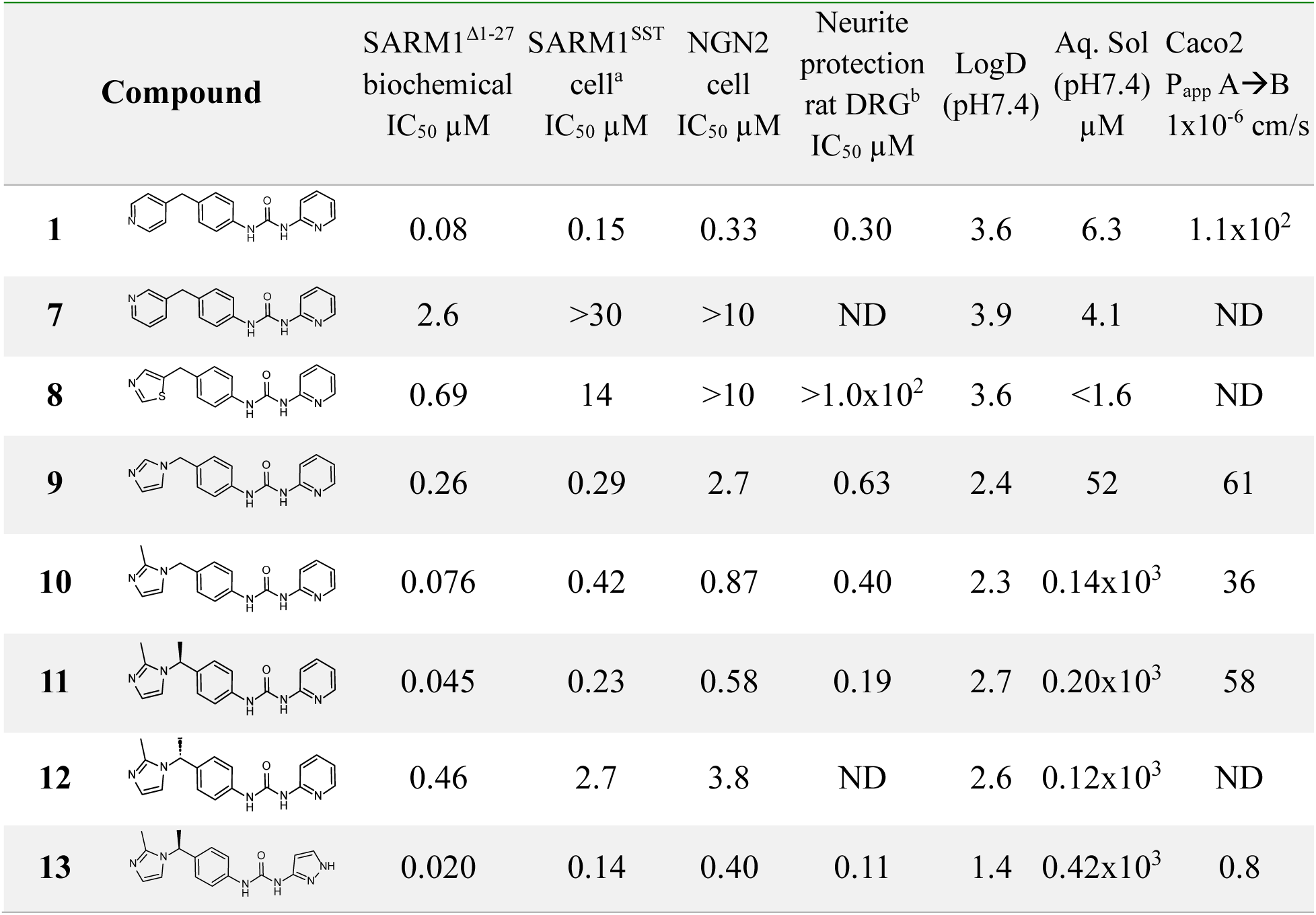
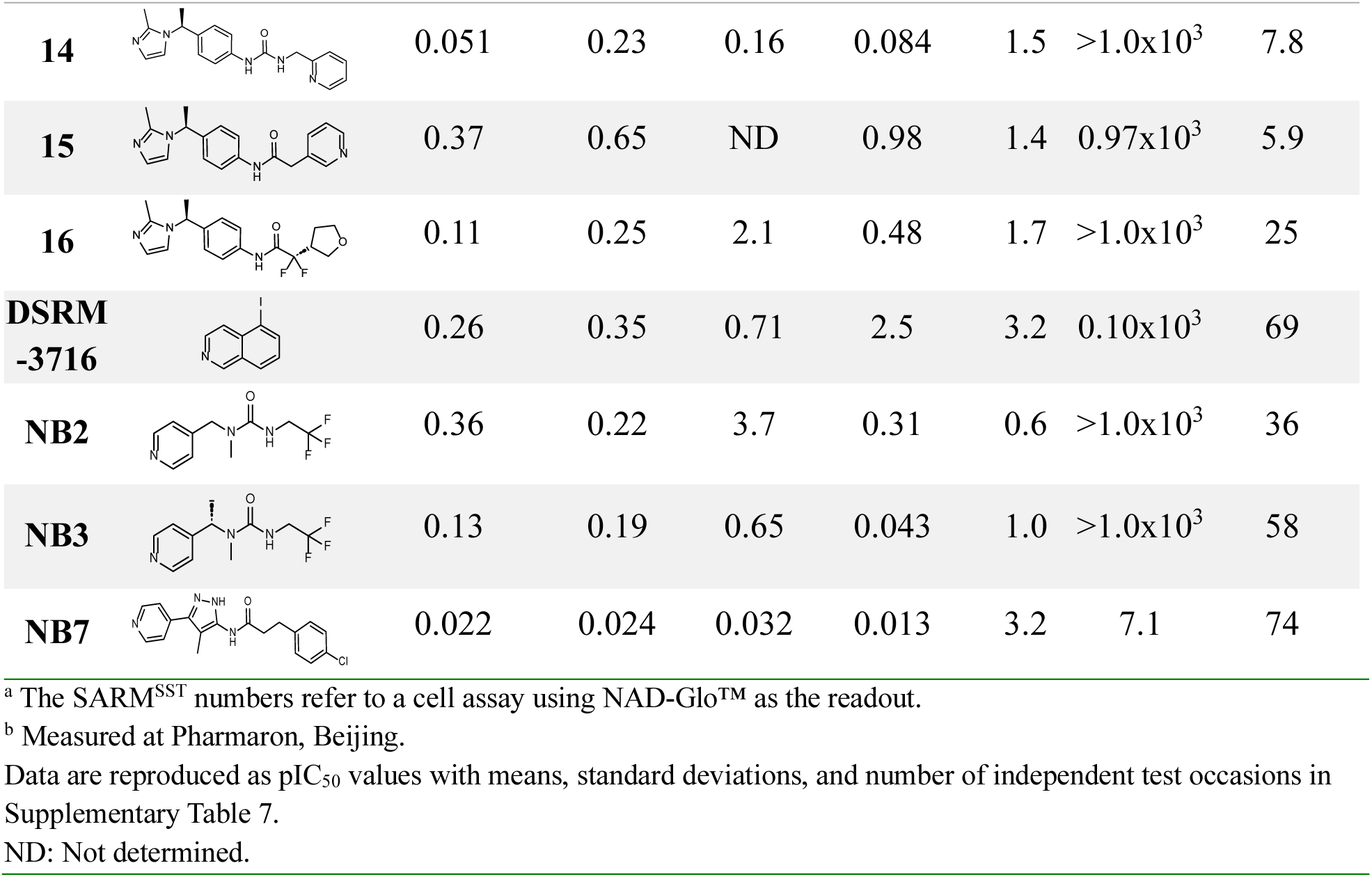
| *In vitro* data for analogs of Compound 1 and Literature compounds.

Whilst the assays in HEK293 cells showed inhibitory activity against constitutively active SARM1 in cells, we wished to demonstrate the activity of compounds in a more disease-relevant context. To this end we developed an assay in primary rat dorsal root ganglion (DRG) neurons using the chemotherapeutic agent vincristine to trigger SARM1 activation (executed at Pharmaron, Beijing). DRGs were isolated from embryos of SD rats (15.5 days post coitus) and cultured for 7 days before treatment with vincristine and SARM1 inhibitors. After 48 h, immunofluorescent staining against beta III-tubulin (TUBB3)^76^ and imaging showed the ability of SARM1 inhibitors to block the degeneration of axons caused by vincristine (Fig. 3C). Applying this assay, we observed reasonable correlation to the biochemical assay based on human recombinant protein across multiple BEXi chemotypes (Fig. 3D), and to the HEK293 cell assay overexpressing SARM1^SST^ (Supplementary Figs. 5C&D).

To ensure translation to a more relevant human cell setting, in which full-length SARM1 is endogenously expressed, we developed an in-house inducible pluripotent stem cell model with doxycycline-inducible expression of neurogenin 2 (hNGN2; described in detail in Supplementary Fig. 6)^77^. This cell model was adapted into 384-well microplate format and combined with an in-house discovered base-exchanging substrate (Supplementary Figs. 2H&I) to afford a cell target engagement assay, where SARM1 activity can be followed by LC-MS, monitoring formation of a base-exchange adduct (Fig. 3E), an approach analogous to the reported SARM1 probes PC6 and 1a^41,78^. To verify specificity of this approach we first applied the SARM^SST^ assay in HEK293 cells and demonstrated SARM1 and time dependent appearance of adduct, while untransfected cells did not produce any detectable base-exchange product (Supplementary Fig. 5E). Temporal control of SARM1 activation in the hNGN2 cells was afforded by adding 10 µM Vacor, a neurotoxin known to be intracellularly metabolised by nicotinamide phosphoribosyltransferase (NAMPT) to a mononucleotide species (VMN) that is a potent allosteric activator of SARM1^79–81^. Using this assay, we demonstrated a Vacor concentration dependence for BEX adduct formation and complete inhibition by **1** applied at 10 µM concentration (Fig. 3E), demonstrating activation and inhibition of SARM1 base-exchange activity, respectively. As illustrated in Fig. 3F and Supplementary Fig. 5F, potency measurements for different BEXi chemotypes in this target engagement assay correlates with those observed in the biochemical assay and the rat DRG assay.

While applying this set of assays for SAR optimisation, as described below, we also assessed reported SARM1 inhibitors as they became public to verify concordance with literature findings. Relevant BEXi include DSRM-3716, NB-2, NB-3, and NB-7 (Table 2), all of which gave equivalent potency in our assays to that reported^53,55^.

### Structure activity relationship of Compound 1 analogs

The biochemical potency of **1** together with its activity against vincristine-induced axon degeneration in rat DRG neurons and its modular structure made it an attractive starting point for further optimisation. In addition, **1** shows no inhibition of CD38, the closest neighbour of SARM1 and another enzyme which produces cADPR (Supplementary Table 5). Compound **1** shows weak inhibition of NAMPT, although lack of inhibition does not exclude it from being a substrate of NAMPT like the neurotoxin Vacor^79–81^. Compound **1** has low solubility, moderately high lipophilicity, originated from a kinase program and has potential for significant off-target pharmacology. In common with many other compounds incorporating an exposed pyridyl group, **1** is a potent inhibitor of cytochrome P450s (CYP) (see Supplementary Table 5). Initial synthetic exploration focused on understanding the importance of the aza nitrogen common to all hit compounds, improving physicochemical properties and the secondary pharmacology profile, to identify a tool compound suitable for studying the pharmacology of SARM1 BEXi.

Comparison of **1** with **7** indicates the importance the vector of the nitrogen lone pair plays in the potency (Table 2). It is noted that changes to potency in general translated well between biochemical and cellular assay formats (see Table 2, Fig. 3, and Supplementary Figs. 2&5), with some notable exceptions. Thiazole **8** and imidazole **9** retain some biochemical activity, indicating that there is some flexibility in the nature and basicity of the aza heterocycle tolerated. However, **8** did not demonstrate good potency in cellular assays. Imidazole **9** retained potency in the SARM1^SST^ cell assay and had reduced lipophilicity and improved solubility compared with **1** but still retained potent inhibition of CYP enzymes (Supplementary Table 6). However, introduction of a 2-methyl substituent to the imidazole to give **10** improved potency and eliminated CYP inhibition. Additional methyl substitution adjacent to the imidazole (**11, 12**) further improved potency and indicated the chiral preference for the (*S*)-isomer, which was also seen with subsequent analogs.

The facile synthesis of the urea allowed rapid exploration of the 2-pyridyl moiety. SAR around this group was tight with greatest potency retained in compounds incorporating a 2-aza heterocycle such as **13** and **14,** which can retain interactions with Tyr687 and Lys694 in a comparable way to the pyridine of **1** (Fig. 2D). Compound **14** had greater solubility and demonstrated higher passive permeability than **13**. A series of classic urea replacements were explored (not shown) but adequate potency could not be retained. Compound **15** represented a promising amide, albeit with around 8-fold drop off from urea **11**. Optimisation of the carboxamide moiety in the amide series resulted in **16**, which had increased potency and improved permeability.

A base-exchange adduct of compound **16** (adduct **17**) was isolated using the biocatalytic procedure described above, confirming that the ability to form isolatable ADPR adducts is a common feature of SARM1 BEXi from both pyridyl and imidazolyl compounds despite differing basicity. Measured pK_a_ (most basic center) for compounds **1**, **13**, and **15** was 5.72, 7.64, and 7.66 respectively (Supplementary Table 6). The cryo-EM structure of adduct **17** bound to SARM1^TIR^ (Fig. 2E & Supplementary Fig. 4G&I) showed the imidazolyl moiety, which is presumably able to delocalise the positive charge, making a π-cation interaction with the sidechain of Trp638^B^. The amide moiety occupies the cleft between TIR^A^ and TIR^B^ and is stabilised by NH interaction with Asn679^A^ and carbonyl interaction with Arg570^B^. The difluoro tetrahydrofuran motif places one fluorine atom proximal to the NH_2_ group of Asn679^A^. The ether oxygen has a stabilizing interaction with the backbone NH of Lys682^A^, rationalizing the ten-fold greater potency of the (*R*) configuration at the tetrahydrofuran, compared to the diastereoisomer (not shown). This differs from the interactions made by the related motif, described as a privileged structure in a recent disclosure of SARM1 BEXi^63^, which interacts with Asp594, Val595, Leu598 and Asn679.

The exploration of compound **1** described above afforded us a set of compounds demonstrating good activity in rat and human neuronal assays with high solubility, good permeability and an optimised profile in terms of CYP inhibition and secondary pharmacology (Supplementary Table 8), which allowed us to characterise the pharmacology of SARM1 BEXi in more detail. Whilst our assay in rat DRG neurons demonstrated the ability of SARM1 BEXi to block axon degeneration caused by 40 nM vincristine following a 48 h treatment, we wished to explore the duration of protection at longer time points with a less aggressive concentration of vincristine. Using the same rat DRG system and time-course live imaging every 4 h we monitored the response to 5 nM vincristine in the presence of compound **14** at concentrations from 0.004 to 10 µM (Fig. 4A). At increasing concentrations above the IC_50_ of 84 nM (at 48 h with 40 nM vincristine) we saw increased suppression of degeneration but even at 10 µM, degeneration is not fully suppressed at 120 h. Reasoning that we may need high multiples of IC_50_ (calculated based on the 48 h time point) to fully suppress degeneration over longer periods, we evaluated the more potent compound **NB-7** (Fig. 4B) at 100 µM and observed a similar incomplete protection at 120 h, despite no compound washout.

**Figure 4.**
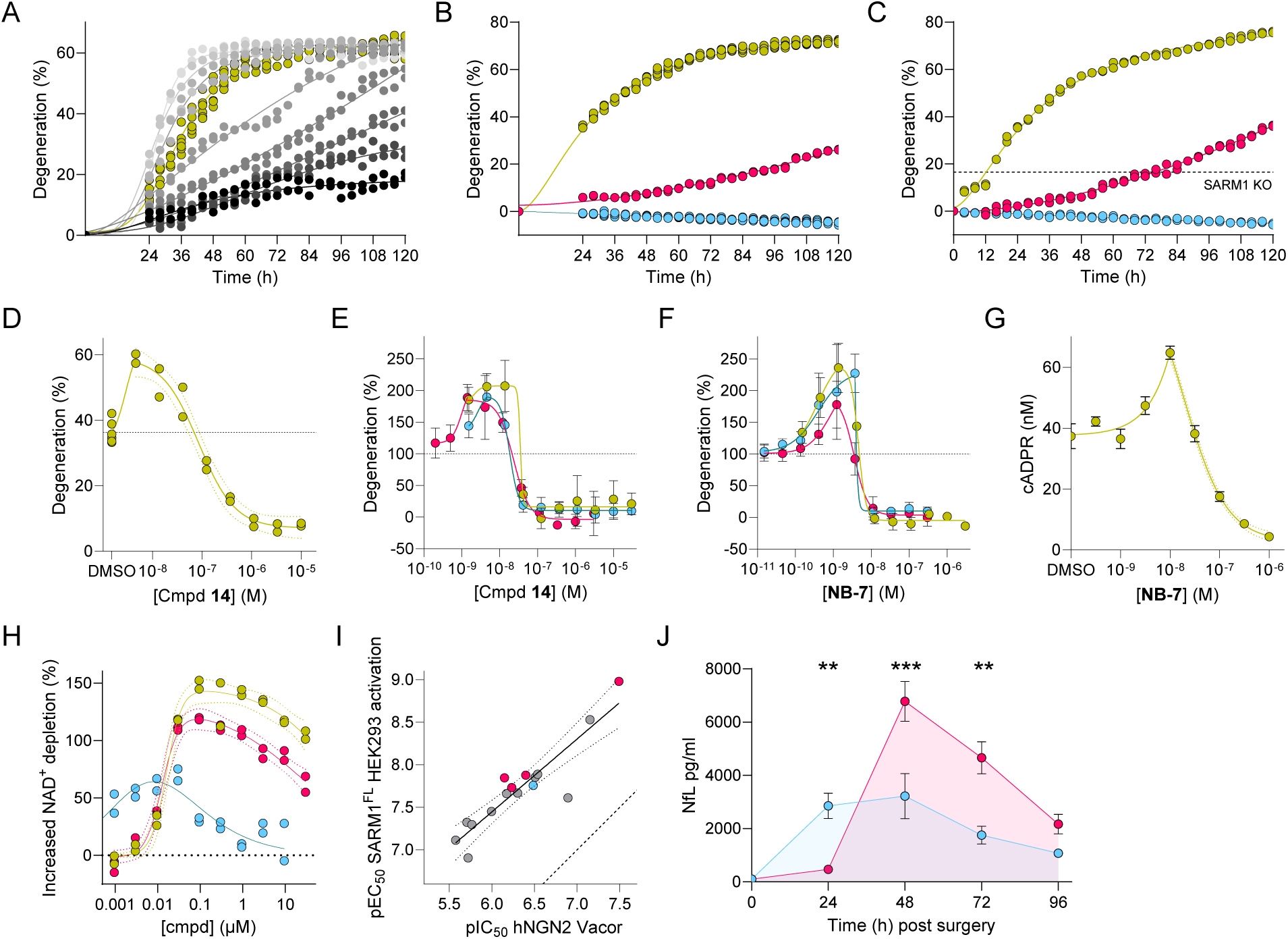
| SARM1 *in vitro* neuronal pharmacology. A) Time course of neurite degeneration in rat DRG neurons treated with 5 nM vincristine and different concentrations of compound **14** over 120 h. The symbols denote DMSO control (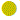) and concentrations from 0.004 (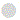) to 10 µM (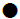). Data are provided with a separate symbol for each technical replicate (n=2; control n=6). B) Time course of axon degeneration in rat DRG neurons treated with 5 nM vincristine only (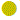) or with 100 µM **NB-7** present over 120 h (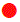), plus a control in the absence of vincristine (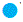). Each technical replicate is shown (n=2; control n=6). C) Time course of axon degeneration in mouse DRG neurons treated with 10 nM vincristine and 250 µM compound **14** over 120 h, with symbols as in 4B. The dashed line represents the level of degeneration observed in SARM1 KO mouse DRGs. D) Degeneration of rat DRG neurons treated with 5 nM vincristine and different concentrations of compound **14** at 40 h (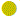). Each technical replicate is illustrated (n=2). The dashed line represents degeneration caused by vincristine alone. E) and F) Treatment effect of SARM1 inhibitors **14** and **NB-7** respectively on neurite degeneration of hNGN2 neurons induced by Vacor (10 µM). The three symbols denote the mean and standard deviations of three technical replicates at three independent test occasions (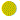,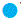,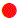). The dashed line represents the extent of neurite degeneration caused by Vacor alone. G) Neuronal concentration of cADPR as a function of **NB-7** concentration. Each data point represents the mean and standard deviation of responses from three technical replicates. H) Concentration response curves of compounds 1 (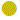), 13 (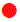), and NB-7 (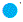) in a HEK293 cell assay with dox-inducible overexpression of SARM1^FL^ and using NAD-GLO™ as readout Each technical replicate is provided (n=2). I) The measured potencies for BEXi induced activation in the HEK293 SARM1^FL^ cell assay (x-axis) correlates with measured inhibitory potencies in the hNGN2 cell target engagement assay (y-axis), albeit with appr. 1.5 log units drop-on across BEXi chemotypes (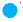, Table 1 hits; 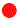, Table 2 compounds including competitor compounds; 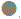, Other BEXi). The solid line represents the best-fit straight line with a slope of 0.86, a correlation coefficient r^2^=0.66 and with the 95% confidence bands shown by dotted lines. The dashed line represents equality. J) Plasma NfL levels in a complete sciatic nerve ligation model in female C57BL/6J mice treated with vehicle (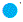) vs **NB-2** at 100 mg/kg (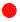). Each data point represents the mean and standard error of mean of responses from eight individual animals in each group.

To investigate the possibility that vincristine can cause degeneration by both SARM1-dependent and independent mechanisms, we switched to an analogous primary mouse DRG model, in which we could determine the extent of protection afforded by SARM1 knockout (Supplementary Fig. 7A). Using the more soluble compound **14** to achieve higher concentrations, Fig. 4C shows that in mouse DRG treated with vincristine, even a concentration of 250 µM (>3 orders of magnitude above the IC_50_) cannot suppress degeneration at 120 h to the level of genetic knockout of SARM1. The titration time course in Fig 4A also showed that at concentrations below the IC_50_ (48 h) the degeneration was accelerated. This was most evident at the 40 h timepoint, as shown in Fig 4D. Having observed an apparent activation in rat DRG treated with vincristine we turned to the hNGN2 cells to explore this phenomenon in human neurons. Time course imaging of neurite length in hNGN2 neurons treated with 10 µM Vacor showed the same phenomenon of activation at sub-critical concentrations, whilst affording protection at higher concentrations, for both compound **14** and **NB-7** (Figs. 4E&F). As shown in Fig. 4G, the increased degeneration caused by **NB-7** was also associated with an increase in neuronal cADPR, indicating the observed phenomenon is a SARM1 dependent effect.

Additional insight as to the direct involvement of SARM1 in the compound-induced accelerated degeneration came from experiments in HEK293 cells stably transfected with an expression vector allowing for doxycycline-induced expression of human SARM^FL^. Titration of doxycycline resulted in a concentration-dependent depletion of intracellular NAD^+^ concentrations (Supplementary Fig. 7B, resulting from increased basal SARM1 activity or concentration-driven higher order forms), while controls in the absence of doxycycline had no change to NAD^+^ levels. As shown in Fig. 4H this depletion of NAD^+^ could be salvaged by **NB-7**, and partly rescued by **1** and **13**. However, at low BEXi concentrations we instead observed a concentration-dependent acceleration of NAD^+^ depletion – in analogy to the observations in DRGs and NGN2 cells (Figs. 4D-G). As illustrated in Fig. 4H the activation effect could be fitted to obtain an approximate EC_50_ value for the activatory effect. To explore the mechanistic reason for this compound-induced activation we compared the potencies from this “aberrant activation” with the inhibitory potencies from the hNGN2 target engagement assay across a broad set of BEXi and observed a strong correlation (Fig. 4I). Given that this comparison is done over multiple SARM1 BEXi chemotypes we concluded that this effect is mediated by activation of SARM1. This was supported by the observation of BEXi-induced activatory effects also in the hNGN2 cell model (Fig. 3E), an effect that was only apparent at low Vacor concentrations where SARM1 is not fully activated by VMN (see Supplementary Fig. 7C).

Having detected aberrant activation of SARM1 by molecules which undergo SARM1 catalysed base exchange in both rodent and human cellular systems we wanted to investigate this class of compounds *in vivo*. The complete sciatic nerve ligation (SNL) model serves as a surrogate model of nerve injury^82^. A significant rise in plasma neurofilament light chain (NfL) is observed between 24-48 h after surgery, indicating an acute axon degeneration, after which the NfL level declines. Previous studies, in the analogous sciatic nerve transection model, have shown that a single administration of NB-2 (100 mg/kg) within 8 h of injury suppresses NfL release at 24 h^55^. We also observed a suppression of NfL at 24 h but at longer timepoints, when exposure levels have dropped and the compound has been cleared, we instead observed an elevated NfL level in the plasma indicative of more significant axonal damage than in the untreated control group (Fig. 4J). These observations are consistent with the aberrant activation of SARM1 observed at sub-efficacious concentrations *in vitro* and also with the reports that BEX inhibitors cause irreversible activation of SARM1^61^. We observed this phenomenon in two independent experiments, performed at independent sites and it is consistent with observations from other groups using NB-2 and NB-3^60,61,63,65,66^.

## Discussion

Since the identification of SARM1 as a central mediator of programmed axonal degeneration triggered under pathological conditions, the concept of SARM1 inhibition has evolved as a promising neuroprotective strategy with the potential to prevent or slow axonal loss across a variety of neurological conditions. At the outset of our study SARM1 was understood to be a multi-domain protein with an auto-inhibited resting state and a catalytic TIR domain NADase, activated upon axonal injury via conformational changes and oligomerization. The detailed molecular structure and mechanisms of allosteric regulation by endogenous ligands were not yet resolved.

Our study successfully identified a new class of SARM1 inhibitors through HTS screening of the AstraZeneca compound library, followed by extensive characterization in both biochemical and cellular assays. Using full length SARM1^Δ1–27^ the screen identified potent inhibitors which also showed cell activity against truncated SARM1^SST^, suggesting inhibition *via* the catalytic domain. Compounds showed equivalent inhibition of SARM1 activated by the detergent DDM as well as the endogenous allosteric activator, NMN. Hit compounds and analogs of compound **1** showed activity in disease-relevant cellular models, namely prevention of vincristine-induced axon degeneration in primary rat dorsal root ganglion neurons and Vacor-induced SARM1 activation and neurite degeneration in human iPSC-derived neurons, confirming their on-target and neuroprotective activity. Optimisation of **1** led to compounds with potency and *in vitro* properties suitable for use as pharmacological probes which we applied, together with other reported SARM1 inhibitors^53, 55^, to explore the mechanism of inhibition and pharmacology of orthosteric SARM1 inhibitors.

We used SARM1 enzyme kinetics to demonstrate that these inhibitors act via an uncompetitive mechanism with respect to the NAD^+^ substrate. Chemical synthesis of adduct **6** allowed us to confirm the adduct as a product of SARM1-mediated base exchange and as the true inhibitory species, which is competitive with NAD^+^. The chemotypes identified in our HTS indicate that a range of compounds containing an aromatic nitrogen atom can perform base exchange. Whilst others have pursued pyridine-based compounds,^55,63,64^ our study focused on imidazole-based compounds, with their potential to access alternative chemical space. We were able to isolate the ADPR adduct generated by biocatalytic conversion of compound **16**, indicating that imidazoles can also form stable adducts. Compounds with a thiazole warhead are known as inhibitors of the related enzyme, CD38^83^ and despite being less nucleophilic than pyridine or imidazole and expected to generate a less stable adduct, compound **8** showed good potency in biochemical assays, but weak potency in cell assays.

The bioconversion of parent compound to an inhibitory ADPR adduct by SARM1 itself raises key questions for a drug discovery program. The concept of generating the inhibitor only in cells in which SARM1 is active is elegant and we hypothesised the conversion of a permeable small molecule to a more polar, less permeable adduct may enhance the cellular retention at the site of action. Despite having cryo-EM structures available, potency optimisation was challenging with changes in structure having only minor impact on activity. We considered this to be consistent with the parent molecule constituting only a small part of the total inhibitory adduct. Our ability to generate and isolate adducts from multiple chemotypes indicate they exist more than transiently and therefore the broader consequence of these exogenous NAD^+^-like molecules on NAD^+^-consuming enzymes must be considered. A recent report indicates that PAD11, an example of an exogenous adduct formed via SARM1 catalysed BEX, can traverse gap junctions and therefore the effects of these adducts may spread to neighbouring neurons^84^. In addition, the difficulty in understanding the rate of conversion of ligand to adduct and overall adduct concentration would make it challenging to determine PK/PD relationships.

Our HDX-MS and SAXS data (Figs. 2A&B) are consistent with the condensed octamer associated with inhibitory NAD^+^ binding and previously determined by cryo-EM by Sporny *et al.*^85^. We also show, in the context of these experiments, that the binding of activator NMN causes only minor conformational changes, most notably at the ARM-TIR interface. By contrast, the data for adduct **6** show a profoundly different conformation induced by binding of orthosteric BEX ligands. The data presented in Fig. 2B is consistent with the model of orthosteric inhibition proposed by Shi *et al.*^57^ and indicates a conformation that is not represented within the natural conformational sampling of the protein. These data are also consistent with a recent pre-print describing a two-step activation of SARM1 by BEXi which starts with an initial activation mediated by NMN leading to the formation of BEX adducts^61^, followed by the adducts acting as molecular glues to promote the self-propagation of filaments containing an active NADase. The HDX-MS data for adduct **6** is also consistent with the cryo-EM structure, indicating that BEX inhibitors occupy a site at the interface of oligomerised TIR domains.

Our cell-based assays displayed concordance with each other and with biochemical data. Notably, we developed a mass spectrometry-based reporter of base exchange activity, using a probe molecule that gave rise to a non-inhibitory adduct. This enabled us to measure inhibition of SARM1 enzyme activity induced by Vacor in human neurons, which correlated with neurite protection. Similarly, inhibition of vincristine induced axonal degeneration in rodent models strongly supports the concept of SARM1 inhibition as disease-modifying in CIPN. However, whilst we could show high levels of protection in assays designed for the screening and optimisation of compounds, we also observed more subtle effects in bespoke assays that were deployed to better understand the time course of SARM1 activation and inhibition. Notably, the neuroprotection conferred by BEX inhibitors in rodent DRGs was incomplete and did not match the effect of SARM1 knock out at extended timepoints, despite excessively high concentrations of the inhibitor (Figs. 4A-C). This suggests exceedingly high levels of target occupancy need to be sustained to achieve complete inhibition.

A more significant observation was that at sub-IC_50_ concentrations inhibitors could, paradoxically, exacerbate axon degeneration (Figs. 4D-F). This observation was supported by a concomitant increase in cADPR, confirming it is a SARM1-dependent effect (Fig. 4G). The phenomenon was observed in both rodent and human neurons and direct involvement of SARM1 could be corroborated using a HEK293 cell model overexpressing SARM1 (Fig. 4H). The same phenomenon appears with all orthosteric BEX inhibitors we have tested (Fig. 4I). Our test set included pyridyl, imidazole, thiazole and aza-biaryl warheads and represented small fragment-like compounds such as **DSRM-3716** and **5** as well as the linear compound **NB-7** and compounds like **14** and **16** with less planar conformations which make different interactions in the orthosteric binding site. The phenomenon is not apparent in assay systems with a high level of SARM1 activation. Hence, we did not observe activation using the constitutively active SST construct or when using high levels of vincristine to drive a rapid degeneration, where there is no headroom to detect further activation. Additionally, the activation caused by BEX inhibitors at sub-efficacious concentration was not observed for the compounds alone but only in the presence of low-level activation of SARM1 (Supplementary Fig. 7D). This shows the importance of careful systematic variation to assay conditions in bespoke mechanistic enzymology and cellular studies to avoid potential pitfalls that may not be apparent in screening assays. Since the conclusion of our studies several groups have reported observations of aberrant activation caused by BEX inhibitors^60–66^.

We rationalised the observed activation at low concentration by considering the formation of an extended array of catalytic sites as shown in Supplementary Fig. 4G. Two TIR domains are required to generate a catalytically competent complex which can generate an exogenous adduct *via* base exchange. Our cryo-EM structures (Fig. 2C-E) suggest that adduct binding can stabilise a third TIR domain which promotes the assembly of additional catalytic sites and increases overall enzyme activity. In this model the occupancy and blockade of one catalytic site creates new ones, like a pharmacological Hydra. The implication is that a high level of target occupancy will be required to fully suppress SARM1 activity using orthosteric inhibitors, as we observed in DRG neurons. Moreover, Wenbin *et al.* have shown elegantly^61^ that BEX inhibitors lead to ADPR-adducts which induce a SARM1 phase transition leading to an irreversibly activated state.

We have shown compelling evidence for the aberrant activation caused by orthosteric SARM1 inhibitors *in vitro*. To investigate whether this phenomenon was present *in vivo* we used the reported tool compound **NB-2** in a sciatic nerve ligation model of neuropathic pain and measured the axonal marker NfL in plasma as indication of nerve damage. Using the reported dose of 100 mg/kg^55^, which is expected to give coverage above the efficacious level for 12 h, we could prevent NfL release into plasma at 24 h (Fig. 4J). However, by 48 h, levels of plasma NfL had spiked to twice the maximum level seen in the untreated group. A delay in degeneration is anticipated whilst SARM1 is inhibited however, the increased maximum and overall area under the curve suggests an enhanced degeneration when full inhibition is lost. We note that additional activation mechanisms can be at play for BEXi with structural resemblance to Vacor and 3-AP, which are known to be converted by intracellular NAMPT to allosteric activators acting at the ARM domain of SARM1^65^. Hence some BEXi can also become the initial trigger of SARM1 activation that is further exacerbated when activated SARM1 forms the dinucleotide adduct in the orthosteric site.

We believe aberrant SARM1 activation at sub-critical inhibitor concentrations represents a crucial translational hurdle. *In vivo* activation of SARM1 mediated by BEX inhibitors at sub-efficacious doses has been observed in rodent models of experimental autoimmune encephalomyelitis (EAE), Vacor mediated toxicity and sciatic nerve ligation. Paradoxical exacerbation of degeneration when drug levels are insufficient suggests that intermittent or suboptimal exposure could worsen patient outcomes. This demands rigorous attention to pharmacokinetics, target engagement and dose selection in the clinical setting. In addition, as the aberrant activation is not detected without some level of SARM1 activation and indeed, the true inhibitory/activatory species is not generated without SARM1 activation, it is possible that safety signals are not detectable in a healthy volunteer population. In summary, whilst inhibiting SARM1 is a promising neuroprotective strategy, the orthosteric BEX class of inhibitors pose a significant safety risk, as shown by our studies and those discussed above. The field now faces the challenge of identifying new mechanisms and modalities for translating this promising biology into safe and effective therapies.

## Methods

### Reagents

#### Expression and purification of SARM1^Δ^^1–27^

The SARM1 cDNA corresponding to residues 28-724 with N-terminal 6His and Avi tags was synthesised (GenScript) and cloned into a pFastBac1 vector. *Spodoptera frugiperda* 9 (Sf9) cells were infected and grown in ExpiSf CD Medium (Thermo Fisher) according to standard protocols. Cells were harvested 48 h post-infection by centrifugation (3400 g, 15 min, 4°C) and resuspended in Buffer A (40 mM HEPES pH 8, 0.4 M NaCl, 8 mM imidazole, 0.008 % Tween-20, 4 % glycerol and 5 mM TCEP) supplemented with 0.04 µL/mL DNAse I and Complete protease inhibitor (1 tablet/50 mL; Roche). The sample was lysed by sonication and clarified by centrifugation (38400 g, 45 min, 4°C). The supernatant was loaded onto a 5 mL HisTrap Crude FF column (Cytiva) pre-equilibrated with Buffer A and eluted with 300 mM imidazole in Buffer A. The eluate was pooled and concentrated to ∼ 11 mL using a 10 kDa MWCO centrifugal device (Pall). The sample was subsequently purified using a Superdex 200 26/60 column (Cytiva) pre-equilibrated with Buffer B (40 mM HEPES pH 8, 0.4 M NaCl, 5 % glycerol and 1 mM TCEP). The peak fractions containing pure SARM1 were pooled, flash-frozen and stored at –80°C. For a description of SARM1^Δ1–27^ immobilisation for biocatalysis, please refer to the Supplementary Information.

The TIR domain cDNA, comprising residues 560-700 and containing a N-terminal 6His-TEV tag, was synthesised (GenScript) and cloned into a pET24 vector. *Escherichia coli* BL21 Gold competent cells (Agilent) were transformed with the TIR plasmid and grown in ZYP-5052 autoinduction media. Expression was induced by lowering the growth temperature to 20°C for 16 h. Cells were harvested by centrifugation (3400 g, 15 min, 4°C) and resuspended in Buffer C (50 mM HEPES pH 8, 0.5 M NaCl, 8 mM imidazole, 0.008 % Tween-20, 4 % glycerol and 5 mM TCEP) supplemented with 0.2 µL/mL DNAse I, 0.1 mg/mL Lysozyme and Complete protease inhibitor (1 tablet/50 mL; Roche). The sample was lysed by sonication and clarified by centrifugation (38400 g, 45 min, 4°C). The supernatant was loaded onto a 5 mL HisTrap Crude FF column (Cytiva) pre-equilibrated with Buffer C and eluted with 500 mM imidazole in Buffer C. The eluate was pooled and buffer exchanged into Buffer D (10 mM HEPES pH 7.5, 0.15 M NaCl, 5 mM imidazole and 1 mM TCEP) using a HiPrep 26/10 desalting column (Cytiva). The 6His tag was subsequently cleaved by overnight incubation with 1:10 (w/w) TEV protease and removed using subtractive IMAC. The resulting flowthrough fractions were pooled, concentrated, and further purified using a Superdex 75 16/60 column (Cytiva) pre-equilibrated with Buffer E (10 mM HEPES pH 7.5 and 0.15 M NaCl). The peak fractions containing pure TIR domain were pooled, flash-frozen, and stored at –80°C.

#### HEK293 cells transiently transfected with SARM1^SST^

cDNA encoding the human SAM-SAM-TIR domains, a constitutively active N-terminal ARM domain truncated SARM1 (409-724), was cloned into the pcDNA3.1 vector (Thermo Fisher Scientific). HEK293 cells were transiently transfected with the SAM-SAM-TIR plasmid (100 µg DNA/1×10^8^ cells) via electroporation on MaxCyte STX (MaxCyte, Rockville). Transiently transfected cells were cryopreserved in DMEM (Dulbecco’s Modified Eagle Medium, 31966 Gibco) supplemented with 10 % FBS (10270 Thermo Fisher Scientific) and 5 % DMSO with 10 million cells/vial. Cryopreserved HEK-SAM-SAM-TIR cells are stored at –150°C until use.

#### HEK293 cells with stable over-expression of doxycycline-inducible human SARM^FL^

cDNA encoding the human SARM1^FL^ protein was cloned into the pcDNA4/TO vector (V1020-20 Thermo Fisher Scientific). HEK293-Trex cells were transfected with human SARM1^FL^ construct. The generation of stable, single cell clone of HEK293-TRex cells expressing inducible SARM1^FL^ protein was carried out in DMEM (11965 Gibco) supplemented with 10% FBS (10270 Thermo Fisher Scientific), 2 mM L-glutamine (25030081 Gibco), 5 µg/ml blasticidin (R210-01 Thermo Fisher Scientific) and 300 µg/ml Zeocin (R250-01 Thermo Fisher Scientific) according to the manufacturer’s protocol. HEK293-TRex SARM1^FL^ cells were expanded and cryopreserved at –150^°^C until use.

#### Cell lysates generated from HEK293-TRex SARM1^FL^ cells

Cryopreserved HEK293-TRex SARM1^FL^ cells were thawed and cultured in DMEM supplemented with 10% FBS and 2mM L-glutamine in T225 flasks for 5 days and the expression of full length SARM1 was induced by culturing the cells in culture medium supplemented with 100 ng/mL of Doxycycline (D9891 Sigma) during the last 24 hours before harvesting. Cells were washed with cold PBS and harvested with Accutase (A1110501 Gibco). Harvested cells were pelleted by centrifugation at 300g for 5 min, washed with cold PBS, and resuspended in cold lysis buffer composed of 10mM Tris-HCl, pH 7.4, 5mM TCEP and one times complete EDTA-free protease inhibitor cocktail (4693159001 Roche). Cells are lysed by sonicating the samples on ice using a Branson Sonifier250 (Thermo Fisher Scientific). Sonicated cell lysates were centrifuged at 16,000g for 30 min at 4°C to pellet any insoluble fraction. The supernatants were collected and aliquoted, stored at –80°C until use. Protein concentration of the cell lysate samples was determined using Pierce BCA protein assay kit (23225 Thermo Fisher Scientific).

#### Cryopreserved iNGN2 cells

The generation and characterization of the human NGN2 cells are described in detail in Supplementary Fig. 6.

### Small molecule inhibitors

Compounds were prepared according to literature procedures or to the experimental procedures described in the Supplemental Material.

### Biochemical assays

Ten microlitres of purified SARM1^Δ1–27^ at the indicated concentration (generally 20-100 nM) was incubated with 25 µM of NAD^+^ and the indicated concentration of compounds in reaction buffer (10 mM Tris-HCl, 25 µM EDTA, 0.01% n-Dodecyl-β-D-maltopyranoside [DDM], pH 7.5) at 25°C in 384 well plates. For inhibitor testing, compound dissolved in DMSO was added to plates prior to addition of SARM1^Δ1–27^ followed by addition of NAD^+^ to initiate reactions. For compounds in concentration response format, 100 nL of compound and a final DMSO concentration of 1% was used. Reactions were quenched after 30-120 minutes by addition of 40 µL of 0.1% formic acid followed by centrifugation for 2 minutes at 2,500 x g. Concentration response experiments included control samples representing no inhibition (DMSO) and full inhibition (N-ethyl-3-imino-N-phenyl-3H-1,2,4-dithiazol-5-amine; inhibitor control compound). Quenched samples were analysed by AMI-MS or LC-MS/MS.

In a second variant of the assay DDM was replaced with 10 µM NMN and the enzyme source was changed from purified recombinant human protein to a HEK293 lysate overexpressing SARM1^FL^ as detailed below. The lysate was applied at a concentration of 0.1 mg/mL measured as total protein concentration, while all other conditions remained the same.

#### Acoustic mist ionization mass spectroscopy (AMI-MS) measurements

Samples in 384 well plates (Labcyte PP-0200) were ionized using an acoustic mist ionization (AMI) source composed of an Echo550 acoustic liquid handler (Labcyte), a high-voltage power supply (RIGOL), and a heated transfer interface (Waters), and introduced into a Xevo G2-XS qToF mass spectrometer (Waters). The transfer interface and mass spectrometer were operated using MassLynx software (Waters). The misting event repetition rate was set at 1400 Hz with a power of 11.5 dB, polarity switching every 10 nL, charging cone voltage at ±3 kV, and transfer interface heated at 200°C. The transit velocity of droplets within the interface was controlled by allowing cone gas flow at 50 L/h. The mass spectrometer was operated in positive ion sensitivity mode with a source temperature of 100 °C, cone voltage of 20 V, and target enhancement at 550. Data were acquired over a range of 500 – 700 m/z. Samples were written into a single data acquisition file and automatically post-processed into an individual mass spectrum for each sample. NAD^+^ and ADPR were quantified by measuring the intensities of [M+H]^+^ at 664.1287 ± 0.04 m/z and 560.0991 ± 0.04 m/z, respectively.

#### LC-MS/MS measurements

Samples (1 µL) were injected into an UPLC system (ACQUITY; Waters) containing an ACQUITY UPLC HSS T3 column (2.1 x 30 mm, 1.8 µm; Waters) kept at 40°C at a flow rate of 1 mL/min in 0.2% acetonitrile with 0.1% formic acid. Mobile phase A was water containing 0.1% formic acid and mobile phase B was acetonitrile containing 0.1% formic acid. Analytes were separated using a linear gradient of 0.2-95% acetonitrile with 0.1% formic acid over 0.1-0.7 min and directed into a Xevo TQ-S triple quadrupole mass spectrometer (Waters) with an electrospray ionization (ESI) source. The source temperature and desolvation temperature were set to 150°C and 450°C, respectively. Mass analysis was done by multiple reaction monitoring (MRM), and MRM parameters were optimized using MassLynx QuanOptimize software (Waters) by flow injection ESI MS of pure analyte standards dissolved in 0.1% formic acid. Peak integration and quantification were done using MassLynx TargetLynx software (Waters).

Data normalization, curve fitting, and calculations of Z’-factor and IC_50_ values were done using Genedata Screener (Genedata) and GraphPad Prism (GraphPad).

### Biophysical assays

#### HDX-MS

The buffer used for the HDX-MS experiments was 40 mM TRIS pH 7.8, 250 mM NaCl, in H_2_O or D_2_O. Prior to the experiments 10 µM SARM1^Δ1–27^ in H_2_O buffer was preincubated with 5 mM of NAD^+^, 5 mM NMN or 0.3 mM of adduct **6** for 30 minutes at room temperature. A reference apo state sample with no compound addition was treated in a similar way. Exchange reactions were carried out using a CTC PAL sample handling robot (LEAP Technologies). Reactions were conducted by incubating 3 μL of protein samples with 57 µL of D_2_O buffer for times of 0.5, 1, 10, and 30 min at 20°C. For the NAD^+^ and NMN samples, the respective nucleotide was added to a concentration of 5 mM in the D_2_O buffer to maintain full occupancy. The exchange reaction was stopped by the addition of 50 μL of quench solution (2 M Urea, 0.1%TFA, pH 3.0) at 0°C. Samples were subsequently injected onto an online pepsin digestion system and subjected to digestion using a BEH pepsin column (Waters) 2.1×30mm in 0.3% formic acid in water at 150 μL min^−1^ and digested peptides were trapped using a 2.1×5mm, 1.7 μm, C18 trap (ACQUITY UPLC BEH C18 VanGuard Pre-Column, Waters) column for 3 min. The desalted peptides were separated and eluted using a C18 reverse phase column (ACQUITY UPLC BEH C18 Column, 1.7 μm, 2.1×100mm, Waters) with a 6 min 5–40% (vol/vol) acetonitrile (containing 0.1% formic acid) gradient at 40 μL min^−1^. Resulting peptides were ionized by electrospray onto a SYNAPT G2-Si mass spectrometer (Waters) acquiring in MS^E^ mode for detection and mass measurements. Peptides from an unlabelled protein were identified using Protein Lynx Global Server 2.0 searches of a protein database containing the SARM1 sequence. Each deuterium labelling experiment was performed in at least triplicate. Relative deuterium levels for each peptide were calculated by subtracting the average mass of the deuterium labelled sample from that of the undeuterated control sample. All mass spectra were processed with DynamX 3.0 (Waters). The data normalisation was calculated with in-house software written in MATLAB (Mathworks). The HDX–MS data were calculated using the mean deuteration level per amino acid, as reported.^86^ The differential deuterium uptake between SARM1-liganded and SARM1-apo states were plotted onto the crystal structure of SARM1 (7cm6) using Pymol (Schrödinger). For details of the TIR domain, a composite model based on 7CM6, and the crystal structure (6O0Q) was used as some regions of the TIR domain were not present in the cryo-EM model. No back-exchange calculations were done. A difference >0.5 Da was considered significant.

#### Small-Angle X-ray Scattering

SAXS measurements were conducted at Diamond Light Source at the 12.4 keV beamline B21^87^ utilizing inline SEC-SAXS. Fifty μL of SARM1^Δ1–27^ at 8 mg/mL in the absence of ligands, with 1 mM NMN or 0.5 mM adduct 6, was loaded onto a Shodex KW-404 column using an Agilent 1200 HPLC system and a running buffer comprising 40 mM HEPES pH 8, 400 mM NaCl, 5 % Glycerol.

Scattering intensity was collected using a EIGER 4 M hybrid photon-counting detector (Dectris Ltd, Baden, Switzerland) in the momentum transfer (*q*) range of 0.0032–0.38 Å^−1^ [*q* = 4π sin(*θ*)/*λ*, where 2*θ* is the scattering angle] at a flow rate of 0.16 mL/min with each dataset comprising 600 frames with 3 seconds of exposure. Data merging, averaging and subtraction were performed using the data processing tool Scatter (https://www.bioisis.net).

#### Cryo-EM studies

SARM1^TIR^ at 6.83 mg/ml was incubated with 2 mM adduct **6** for 1 hour, and 2.5 µL vitrified on Quantifoil R1.2/1.3 Cu 200 mesh grids made hydrophilic with a 20 mA 120s glow discharge (Quorum GloQube) on a Vitrobot Mark IV (Thermo Fisher Scientific) at 4°C, 100% relative humidity, blot time 5s, blot force –7 (an arbitrary value matching the calibration procedure as described previously)^88^.

Adduct **17** was prepared in a similar way with 4.3 mg/ml SARM1^TIR^ incubated with 1 mM adduct **17** for 1 hour, and vitrification of 2.5 µL, with the addition of the use of a 2 nm carbon support layer (Quantifoil R1.2/1.3 Cu 300 mesh 2 nm carbon), which improved the yield of helical filaments and improved the ice thickness control. A shorter glow discharge time of 10s at 20 mA on the GloQube was required to avoid disrupting the thin carbon layer. Other parameters of the vitrification on the Vitrobot were similar with 4°C, 100% relative humidity, blot force –5, and blot time 6s.

Data was collected on a Titan Krios G3 equipped with a Falcon 4i detector for adduct **6** (3370 movies – Supplementary Fig. 4A) and a Titan Krios G4 equipped with Falcon 4i for adduct **17** (18,317 movies) at 120k (resulting in pixel sixes of 0.656 and 0.642, calibrated using Ultrafoil gold images^89^ with aberration free image shift and fringe free imaging using a fluence rate of 4.25e/px/s and total fluence of 43e-/ Å^2^ for adduct **6** and fluence rate of 5.25e/px/s and total fluence of 60e/ Å^2^ for adduct **17**.

Image processing was performed in CryoSparc V 4.7.1^90^. Pre-processing was performed in CryoSparc Live, with particle picking using both blob picking and template picking approaches, with template picking allowing better sampling of the potentially large variable helical pitch filamentous particles. 2D class averaging was performed to remove bad particles, and to remove duplicate particles. The large pitch of the filaments allowed a large coverage of views of the filament (Supplementary Fig. 4C), and therefore standard single particle processing was used for 3D reconstruction^57,75^. Ab initio model generation with three classes was used (large starting and final batch sizes –8000). The resulting good class was then refined using non-uniform refinement with a large window size – smaller windows and local refinement using small mass to avoid non-uniformity of the filaments was attempted but not found to increase the map quality significantly. Additionally imposing C2 symmetry in non-uniform refinement was found to increase the overall resolution (data not shown), as expected from TIR-paired filaments, but not pursued due to possible local differences at the interface of the TIR domains. The filaments are also likely helical, and helical symmetry could be used to increase the number of subunits being averaged to increase the resolution, with a similar risk to imposing C2 symmetry. We did not pursue this for our maps of TIR-adduct **6**,**17** as both have local resolutions around the adducts at around 2.4Å, sufficient to explore the chemistry of adduct binding.

### Cellular assays

#### In vitro cell assay in HEK293 cells transiently transfected with SARM1^SST^

Cryopreserved HEK293 cells transiently transfected with SARM1^SST^ were thawed, resuspended in DMEM supplemented with 10 % FBS, seeded at a density of 0.6 x 10^6^ cells/mL with 25 µL/well in 384-well cell plates (781091 Grenier) and incubated for 20-24 hours at 37°C prior to compound dosing. Seventy-five nL of DMSO neutral control or compounds dissolved in DMSO were dosed to appropriate wells on cell plates on an Echo 655 (Labcyte Inc, San Jose, CA) liquid dispenser. One set of plates were used for NAD/NADH-Glo™ assay, where the cells were treated with compounds for 4 hours at 37°C before being lysed with 15 µL of lysis solution composed of 0.13 N NaOH (320331 Sigma) supplemented with 1.3 % DTAB (D8638 Sigma) followed by adding 20 µL of 0.4 N HCl (415413 Sigma). The amount of NAD^+^ in cell lysates was quantified using the NAD/NADH-Glo™ assay kit (G9072 Promega) following the manufactures’ recommendation. NADH existing in lysate samples were destroyed by heating sealed plates at 60°C for 15-20 minutes on a plate heater. The acid was neutralized by adding 20 µL of 0.4 M of Trizma Base solution (T1503 Sigma) to acid treated samples. To reduce the amount of NAD/NADH-Glo™ detection reagent needed, we set up a miniaturised detection format by transferring 4 µL of lysate samples to 384-well white low-volume HiBASE detection plates (784075 Grenier) on a Bravo Liquid Handling platform (Agilent Technologies, Singapore) followed by adding 4 µL of NAD/NADH-Glo™ detection reagent. The plates containing lysate samples and NAD/NADH detection reagent were incubated at room temperature for 30-60 minutes, and luminescence signal was detected on a PHERAstar FS plate reader (BMG Labtech, Ortenberg, Germany). The amount of luminescence detected is proportional to the amount of NAD^+^ in a sample.

For the CellTiter-Glo® Luminescent Cell Viability Assay, a second set of the SARM1^SST^ plates were treated with 10 µL of the CellTiter-Glo® Substrate and Buffer mixture from the CellTiter-Glo® Luminescent Cell Viability Assay kit (G7570) and incubated at room temperature for 30 mins, following the manufactures’ recommendation. Luminescence signal was detected on a PHERAstar FS plate reader (BMG Labtech, Ortenberg, Germany). The amount of luminescence detected is proportional to the amount of ATP in a sample and is an indicator of metabolically active live cells.

#### In vitro cell assay in HEK293-TRex SARM1^FL^ cells

Cultured or cryopreserved HEK293-TRex SARM1^FL^ cells were resuspended in DMEM supplemented with 1% FBS and indicated concentrations of Doxycycline (D9891 Sigma), seeded at a density of 0.4 x 10^6^ cells/mL with 25 µL/well in 384-well cell plates (781091 Grenier). The cell plates were incubated for 6 or 24 hours at 37°C to induce the expression of the full length human SARM1 protein before compound treatments and lysed. NAD^+^ in cell lysates was quantified as described above.

#### iNGN2 target engagement assay

Cryopreserved iNGN2 cells were seeded at a density of 0.7 x 10^6^ cells/mL at 25 µL/well volume in 384-well PhenoPlate, poly-D-lysine-coated culture plates (PerkinElmer 6057500) in Advanced DMEM/F12 (Thermo Fisher, 12634-010) supplemented with N-2 Supplement (Thermo Fisher, 17502-048), Y-27632 dihydrochloride (Tocris Bioscience 1254) and doxycycline (Sigma D9891), and cultured for 24hrs. Cultured cells were co-treated with 10 µM Vacor and 5 µM base-exchanging substrate along with BEXi compounds for a further 24 hrs before cell lysis with AlphaLISA Assay Buffer (Revvity AL000C). The cell lysate was applied to LC-MS/MS (Waters UPLC TQS system) and analysed using TargetLynx software (Waters) to integrate the MS peaks as described above in the LC-MS/MS measurements section. Data normalization, curve fitting, and calculations of Z’-factor and IC_50_ values were done using Genedata Screener (Genedata) and GraphPad Prism (GraphPad).

#### iNGN2 Neurite degeneration assay

Cryopreserved iNGN2 cells were seeded at a density of 0.05 x 10^6^ per cm^2^ in 96-well plates pre-coated with poly-D-lysine (PDL) (Grenier, 655946), poly(ethyleneimine) (PEI) (Sigma, P3143), and Matrigel (Corning, 354230) and grown for 7 days *in vitro* (DIV) in NeuroBasal Plus growth media (Thermo Fisher, A3582901) supplemented with B27 (Thermo Fisher, 17504044), Glutamax (Thermo Fisher, 35050061), neurotrophic factors (PeproTech, AF-450-10, AF-450-02, and 450-03), doxycycline (Sigma, D9891), heparin (Sigma, H3393), DAPT (Tocris, 1254) and cytarabine (Sigma, C1768). Cultured cells were co-treated with 10 µM Vacor and half-log serial dilutions of SARM1 BEXi and immediately placed into an S3 IncuCyte® Live-cell Analysis System. Phase images were acquired every 4 hours for 48 hours. The Incucyte NeuroTrack software module was applied to quantify mean neurite length per well (mm/mm^2^). Neurite length data for a given timepoint were normalised so that the mean of the healthy vehicle control condition defined 0% degeneration and the mean of the 10 µM Vacor condition defined 100% degeneration.

For cADPR measurements the cells were instead washed and plates followed by freezing at –70°C pending analysis. cADPR in the NGN2 cells was quantitated by liquid chromatography coupled with triple quadrupole mass spectrometry (LC-MS/MS). A stock solution of cADPR at 100 µM was prepared in UPLC grade water (Elga), using solid reference material from Merck (C7344). Dilutions of this stock were performed to create calibration standards and QCs in a solution of 10 % formic acid (Sigma/Merck) in UPLC grade water, over a range of 1-200 nM, and 30 µL of each were added to a 96 well 1 mL lo-bind plate (Eppendorf). Plates containing cells were removed from storage in an ultra-freezer set to –70 °C, before addition of 30 µL 10% formic acid in water to each well and mixing at 850 rpm in a thermomixer (Eppendorf) set at +4 °C for 30 minutes. After mixing, 60 µL of a solution of 3 µg/mL Adenosine-15N5 5-Monophosphate (AMP-15N5) (Merck 662658) in 10% formic acid in water was added to each well as an internal standard. Both plates were mixed at 850 rpm for 1 minute before 30 µL was transferred from each well into a fresh 96 well 1 mL lo-bind plate. To this plate, 200 µL of UPLC grade acetonitrile (Merck) was added to each well and the plate mixed at 1000 rpm for 1 minute and centrifuged at 2000 G for 3 minutes. 200 µL was then transferred to another fresh 96 well 1 mL lo-bind plate, and evaporated using nitrogen in a Minivap (porvair) set at 50 °C. Once dry, 100 µL was added to each well of the plate, which was then mixed at 1000 rpm for 2 minutes, and centrifuged at 2000 G for 3 minutes before analysis via LC-MS/MS.

Liquid Chromatography was performed on a Acquity I-Class (Waters), on an Atlantis T3 3uM 3×100m column (Waters) on a gradient using 0.2% formic acid in water and 0.2% formic acid in LC-MS/MS grade acetonitrile as mobile phases. 10 µL was injected into the flow of 0.6 mL/minute, and an analytical gradient starting at 100% aqueous phase for 0.2 minutes, before increasing the organic phase to 3.5% from 0.2 to 2 minutes was run. The column was flushed with 95% organic and re-equilibrated before subsequent injections, maintaining the flow rate throughout. A Sciex 6500+ triple quadrupole Mass Spectrometer (Sciex) was set to acquire using multiple reaction monitoring (MRM) mode, using transitions and settings as detailed below. Source settings were: 600 °C temperature, 5500 V ion spray voltage, GS1 and GS2 set to 40 psi, and CAD set to Low.

| ID | Q1 (m/z) | Q3 (m/z) | DP | CE | Retention time (mins) |
| --- | --- | --- | --- | --- | --- |
| cADPR | 542.3 | 428.1 | 100 | 28 | 1.8 |
| cADPR | 542.3 | 136.1 | 100 | 44 | 1.8 |
| AMP-15N5 | 397.1 | 125.3 | 100 | 30 | 1.6 |

Data processing was performed using Analyst v1.7.2 (Sciex) using peak area ratio to generate a 1/x2 linear regression of the 542.3>428.1 transition for quantification, whilst monitoring the 542.3>136.1 transition.

#### Primary rodent dorsal root ganglia (DRG) neuron culture

Each well of a 96-well plate was coated with poly-L-lysine at room temperature overnight. After aspirating the solution, the wells were washed with Dulbecco’s Phosphate-Buffered Saline (DPBS) and air-dried. They were then incubated with laminin solution (5 μg/mL in DPBS) for at least two hours at 37°C. Just before plating neurons, the laminin solution was removed, and the wells were washed with DPBS and air-dried.

DRGs were isolated from pregnant Sprague-Dawley (15.5 days post-coitus) and C57BL (13.5 days post-coitus) rats and mice, respectively. Both animals were euthanized through CO_2_ asphyxiation followed by cervical dislocation. The DRGs were extracted from the embryos and kept on ice in Leibovitz’s L-15 medium. For the rat DRGs, dissociation was achieved using TrypLE Express at 37°C for 30 minutes. The mouse DRGs were dissociated using the Papain Dissociation System Kits for approximately 10 minutes according to the manufacturer’s instructions. After dissociation, both DRG preparations were filtered through a 100 μm cell strainer. For the mouse DRGs, an albumin-inhibitor solution was added before filtering. The rat DRGs were directly resuspended in a neurobasal complete medium with supplements: B-27, L-glutamine, 5-fluoro-2’-deoxyuridine, uridine, nerve growth factor, and antibiotics. For the mice, dissociated DRGs were briefly incubated in a pre-coated petri dish at 37°C for 2 minutes before centrifugation at 1000 rpm for 5 minutes, then resuspended in a complete medium. For both cultures, the cells were counted and diluted to a concentration of 1 x 10^7^ cells/mL. Subsequently, 0.5 μL of the cell suspension was aliquoted into each well of pre-coated 96-well plates. Following an initial incubation at 37°C (rats for 10-15 minutes, mice for 10 minutes), 100 μL of complete medium was added to each well. Both cultures were maintained at 37°C, with rat cultures incubated for 7 days and mouse cultures for 10 days before compound treatment.

#### Live image analysis of primary DRG neurons

Primary rat and mouse DRG neurons were co-treated with Vincristine, at concentrations of 5 nM for rat neurons and 10 nM for mouse neurons, and serial dilutions of an SARM1 inhibitor. Neurite density was assessed using the IncuCyte® Live-cell Analysis System every 4 hours for 5 days. For each well of the 96-well plate, nine images were acquired. The neurite density (*L*), defined as the length of nerve fibers per unit area, was quantified using the neuro track module of the IncuCyte® system based on the formula:

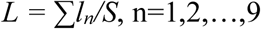

where *l_n_* is the total neurite length in each image, and *S* is the total imaged area.

Recognizing that neurite densities can vary significantly between different wells and images, a longitudinal and time-dependent parameter, *P_T_*, was utilized to evaluate the degree of neurodegeneration. For any given well, the percentage of degeneration (% Degeneration) was calculated using the formula:

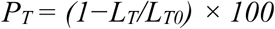

In this equation, *L_T0_* is the initial length of nerve fibers per unit area, and *L_T_* is the length of nerve fibers per unit area at the time point of detection. This calculation provides a quantitative measure of neurodegeneration over time, facilitating a better understanding of changes in neurite integrity under varying experimental conditions.

#### Immunocytochemistry of primary rat DRG neurons

Rat primary DRG neurons were co-treated with 40 nM of Vincristine and serially diluted SARM1 inhibitors. Forty-eight hours later, the medium was removed, and wells were washed twice with 100 μL of DPBS. Cells were fixed with 40 μL of 4% PFA for 10 minutes at room temperature. Wells were then washed three times with 100 μL of ice-cold DPBS. Permeabilization was performed using 40 μL of 0.5% Triton® X-100 for 10 minutes, followed by three 5-minute washes with 100 μL of DPBS. Blocking occurred for 30 minutes in DPBS with 5% FBS, 2% BSA, and 0.1% Tween-20. Anti-beta III Tubulin antibody (ab41489) and Anti-NeuN antibody (ab104225) were incubated overnight at 4°C. Wells were washed three times with 100 μL of DPBS (5 minutes each), then incubated with secondary antibodies for 2 hours at room temperature. Four 5-minute washes with 100 μL of DPBS followed. Each well was sealed with 50 μL of 90% glycerol, and images were acquired using a High Content Analysis System (Operetta CLSTM, PerkinElmer).

### In vivo studies

#### Animals for in vivo experiments

The studies were performed with approval of the Animal Welfare and Ethical Review Body at the University of Cambridge, under the UK Home Office establishment licence X81BD37B1, UK Home Office project licences PB943A152 and PPP4335588, and the Animal (Scientific Procedures) Act 1986. All animals were obtained from Charles River, UK. After arrival, animals were housed in individual cages lined with aspen wood chip bedding, along with provision of artificial enrichment. Food (Safe Diets, R105) and water were provided to all animals ad libitum. Cages were ventilated at the rate of 75 air changes per hour. All animals were allowed to acclimatise for 7 days before any procedure was undertaken.

The holding rooms were maintained at a temperature of 19 to 23°C, humidity of 45 to 65 %, with 15 to 20 air changes per hour. The room was provided with artificial light of 60 lux between 7 am and 7 pm, with all procedures being conducted during this time. All studies were conducted using female C57BL/6J mice. Animals were of 6 to 8 weeks of age and above 18g at the start of the study.

After the acclimatisation period, all animals underwent Uno Pico-ID (Uno life science solutions; 15000050) insertion for identification purposes, post which the animals were allowed to recover for 7 days before handling was resumed. All animals were weighed weekly upon arrival, and daily for the duration of the study. A weight loss threshold of 15% from pre-surgery was set as a threshold. Power analysis was performed using data obtained from a pilot study, based on which we have used 8 animals per group per time point in all studies.

#### Complete sciatic nerve ligation model

All animals underwent surgery for complete sciatic nerve ligation. No sham surgeries were performed due to project licence restrictions. Mice were anaesthetised with inhalational isoflurane (Isoflo®, Zoetis). An approximately 1cm incision was placed at the level of the left mid-thigh and the sciatic nerve exposed and isolated by blunt dissection. A suture (8/0 Vicryl: Ethicon; catalogue number J548G) was then tied around the entire nerve tightly. The incision was then closed using GLUture topical tissue adhesive (World Precision Instruments, catalogue number 503763).

#### Formulation and dosing of NB2

The vehicle for formulated by magnetically stirring 20% w/v Sulfobutylether-β-cyclodextrin (SBE-β-CD; CAS 182410-00-0) with 100% v/v purified water (CAS 7732-18-5). This was added to 5% polyethylene glycol 400 (PEG-400; CAS 25322-68-3) and pH adjusted to 5-7 using 1M hydrochloric acid (CAS 7647-01-0). NB2 was formulated in 5% v/v PEG 400 / 95% v/v (20% w/v SBE-β-CD in purified water) [pH 5 – 7] and dosed orally using disposable polypropylene feeding tubes (Instech; FTP-20-30), 30 minutes post-surgery at 100 mg/kg with a dosing volume of 10 mL/kg.

#### Plasma separation from whole blood

In-life blood samples were performed at pre-determined time points pre– and post-surgery. A 50 µL K3 EDTA Minivette® (Sarstedt) was used to collect tail vein microsample in-life. Terminal samples were collected by performing cardiac bleeds under anaesthesia, using a 25G needle (BD Microlance™, 300600) attached to a 1 mL Terumo™ syringe (Scientific Laboratory Supplies, SYR6200). The collected blood was then transferred to 1.3ml K3 EDTA tubes for plasma separation by centrifugation at a speed of 1500xg for 10 minutes at 4°C.

#### ELISA assay for NfL measurement

NfL in the plasma samples were determined using the NF-light™ serum ELISA assay obtained from Uman Diagnostics (Catalogue number 20-8002 RUO) in accordance with the manufacturer’s instructions, and measured on the Envision plate reader (Perkin Elmer). Dilution factors for the different groups and timepoints were determined based on analysis from a pilot study.

## Data Availability

The cryo-EM structures in this study will be deposited to RSCB-PDB. All data supporting the findings of this study are available within the paper and its Supplementary Information. The source data underlying all figures are provided as a Source Data file.

## Acknowledgements

We would like to thank Lorraine Miller, Claire Fisher and Matthew Legate (Animal Sciences and Technologies, AstraZeneca UK) for their earnest support in conducting the *in vivo* studies. This work was funded by the BioPharmaceuticals R&D organisation at AstraZeneca.

## Author Contributions

R.J., Q.W. and I.C. provided project context and R.J. and T.L. supervised the work. B.P., E.N., and C.R. generated the purified recombinant human proteins. P.N. and C.G. developed the biochemical assays and performed and analysed the mechanism of action studies in this setting. H.P. and R.M. planned and performed the HTS campaign. M.D. and V.C. developed the SAR-driving cellular assays and V.C. performed and analysed the mechanism of action studies in these assays. M.P. and P.B. performed medicinal chemistry design and analysis. H.J., J.L. and G.F. planned, supervised, and analysed the data generated in the DRG cell models – with the data generated at Pharmaron (Beijing). A.P.-K., S.M., T.M.S., B.B., and M.M. designed, generated, and characterised the hNGN2 cell model. R.M. performed and analysed the data in the differentiated hNGN2 cell model. E.F. profiled compounds in safety assays. T.J. profiled compounds in PK assays. P.M performed *in vivo* studies. N.M. and R.L. performed analysis of NfL and cADPR. K.S., L.W. and T.M.O. performed and analysed the cryo-EM studies. C.J. and J.S. planned, performed, and analysed the HDX studies.

M.L. performed and analysed the SAXS studies. T.S. and B.P. performed the base-exchange adduct synthesis based on immobilised recombinant SARM1. T.L. and R.J. wrote the first draft of the manuscript, and all authors contributed to and approved the final version.

## Materials & Correspondence

Thomas Lundbäck, Rebecca Jarvis

## Tables

Integrated in manuscript to facilitate review.

## Figures

Integrated in manuscript to facilitate review.

## References

1. Wang, J.T., Medress, Z.A. & Barres, B.A. Axon degeneration: molecular mechanisms of a self-destruction pathway. J Cell Biol 196, 7–18 (2012).

2. Lingor, P., Koch, J.C., Tonges, L. & Bahr, M. Axonal degeneration as a therapeutic target in the CNS. Cell Tissue Res 349, 289–311 (2012).

3. Salvadores, N., Sanhueza, M., Manque, P. & Court, F.A. Axonal Degeneration during Aging and Its Functional Role in Neurodegenerative Disorders. Front Neurosci 11, 451 (2017).

4. Waller, A. XX. Experiments on the section of the glossopharyngeal and hypoglossal nerves of the frog, and observations of the alterations produced thereby in the structure of their primitive fibres. Philosophical Transactions of the Royal Society of London 140, 423–429 (1850).

5. Freeman, M.R. Signaling mechanisms regulating Wallerian degeneration. Curr Opin Neurobiol 27, 224–31 (2014).

6. Coleman, M.P. & Hoke, A. Programmed axon degeneration: from mouse to mechanism to medicine. Nat Rev Neurosci 21, 183–196 (2020).

7. Arthur-Farraj, P. & Coleman, M.P. Lessons from Injury: How Nerve Injury Studies Reveal Basic Biological Mechanisms and Therapeutic Opportunities for Peripheral Nerve Diseases. Neurotherapeutics 18, 2200–2221 (2021).

8. Coleman, M.P. Axon Biology in ALS: Mechanisms of Axon Degeneration and Prospects for Therapy. Neurotherapeutics 19, 1133–1144 (2022).

9. Alexandris, A. & Koliatsos, V.E. NAD+, Axonal Maintenance, and Neurological Disease. Antioxid Redox Signal (2023).

10. Seretny, M. et al. Incidence, prevalence, and predictors of chemotherapy-induced peripheral neuropathy: A systematic review and meta-analysis. Pain 155, 2461–2470 (2014).

11. Wang, M. et al. Redefining chemotherapy-induced peripheral neuropathy through symptom cluster analysis and patient-reported outcome data over time. BMC Cancer 19, 1151 (2019).

12. Molinares, D., Kurtevski, S. & Zhu, Y. Chemotherapy-Induced Peripheral Neuropathy: Diagnosis, Agents, General Clinical Presentation, and Treatments. Curr Oncol Rep 25, 1227–1235 (2023).

13. Suzuki, K. et al. Neurological Outcomes of Chemotherapy-Induced Peripheral Neuropathy in Patients With Cancer: A Systematic Review and Meta-Analysis. Integr Cancer Ther 22, 15347354231185110 (2023).

14. Maihofner, C., Diel, I., Tesch, H., Quandel, T. & Baron, R. Chemotherapy-induced peripheral neuropathy (CIPN): current therapies and topical treatment option with high-concentration capsaicin. Support Care Cancer 29, 4223–4238 (2021).

15. Lunn, E.R., Perry, V.H., Brown, M.C., Rosen, H. & Gordon, S. Absence of Wallerian Degeneration does not Hinder Regeneration in Peripheral Nerve. Eur J Neurosci 1, 27–33 (1989).

16. Coleman, M.P. et al. An 85-kb tandem triplication in the slow Wallerian degeneration (Wlds) mouse. Proc Natl Acad Sci U S A 95, 9985–90 (1998).

17. Conforti, L. et al. A Ufd2/D4Cole1e chimeric protein and overexpression of Rbp7 in the slow Wallerian degeneration (WldS) mouse. Proc Natl Acad Sci U S A 97, 11377–82 (2000).

18. Mack, T.G. et al. Wallerian degeneration of injured axons and synapses is delayed by a Ube4b/Nmnat chimeric gene. Nat Neurosci 4, 1199–206 (2001).

19. Beirowski, B. et al. Non-nuclear Wld(S) determines its neuroprotective efficacy for axons and synapses in vivo. J Neurosci 29, 653–68 (2009).

20. Babetto, E. et al. Targeting NMNAT1 to axons and synapses transforms its neuroprotective potency in vivo. J Neurosci 30, 13291–304 (2010).

21. Milde, S., Gilley, J. & Coleman, M.P. Subcellular localization determines the stability and axon protective capacity of axon survival factor Nmnat2. PLoS Biol 11, e1001539 (2013).

22. Loreto, A. et al. Mitochondrial impairment activates the Wallerian pathway through depletion of NMNAT2 leading to SARM1-dependent axon degeneration. Neurobiol Dis 134, 104678 (2020).

23. Araki, T., Sasaki, Y. & Milbrandt, J. Increased nuclear NAD biosynthesis and SIRT1 activation prevent axonal degeneration. Science 305, 1010–3 (2004).

24. Wang, J. et al. A local mechanism mediates NAD-dependent protection of axon degeneration. J Cell Biol 170, 349–55 (2005).

25. Sasaki, Y., Araki, T. & Milbrandt, J. Stimulation of nicotinamide adenine dinucleotide biosynthetic pathways delays axonal degeneration after axotomy. J Neurosci 26, 8484–91 (2006).

26. Osterloh, J.M. et al. dSarm/Sarm1 is required for activation of an injury-induced axon death pathway. Science 337, 481–4 (2012).

27. Gerdts, J., Summers, D.W., Sasaki, Y., DiAntonio, A. & Milbrandt, J. Sarm1-mediated axon degeneration requires both SAM and TIR interactions. J Neurosci 33, 13569–80 (2013).

28. Gerdts, J., Summers, D.W., Milbrandt, J. & DiAntonio, A. Axon Self-Destruction: New Links among SARM1, MAPKs, and NAD+ Metabolism. Neuron 89, 449–60 (2016).

29. Loreto, A., Merlini, E. & Coleman, M.P. Programmed axon death: a promising target for treating retinal and optic nerve disorders. Eye (Lond) (2024).

30. Conforti, L., Gilley, J. & Coleman, M.P. Wallerian degeneration: an emerging axon death pathway linking injury and disease. Nat Rev Neurosci 15, 394–409 (2014).

31. Sporny, M. et al. Structural Evidence for an Octameric Ring Arrangement of SARM1. J Mol Biol 431, 3591–3605 (2019).

32. Horsefield, S. et al. NAD(+) cleavage activity by animal and plant TIR domains in cell death pathways. Science 365, 793–799 (2019).

33. Di Stefano, M. et al. A rise in NAD precursor nicotinamide mononucleotide (NMN) after injury promotes axon degeneration. Cell Death Differ 22, 731–42 (2015).

34. Bratkowski, M. et al. Structural and Mechanistic Regulation of the Pro-degenerative NAD Hydrolase SARM1. Cell Rep 32, 107999 (2020).

35. Figley, M.D. et al. SARM1 is a metabolic sensor activated by an increased NMN/NAD(+) ratio to trigger axon degeneration. Neuron 109, 1118–1136 e11 (2021).

36. Sasaki, Y. et al. Nicotinic acid mononucleotide is an allosteric SARM1 inhibitor promoting axonal protection. Exp Neurol 345, 113842 (2021).

37. Waller, T.J. & Collins, C.A. An NAD+/NMN balancing act by SARM1 and NMNAT2 controls axonal degeneration. Neuron 109, 1067–1069 (2021).

38. Loreto, A., Antoniou, C., Merlini, E., Gilley, J. & Coleman, M.P. NMN: The NAD precursor at the intersection between axon degeneration and anti-ageing therapies. Neurosci Res 197, 18–24 (2023).

39. Antoniou, C. et al. Chronically Low NMNAT2 Expression Causes Sub-lethal SARM1 Activation and Altered Response to Nicotinamide Riboside in Axons. Mol Neurobiol (2024).

40. Essuman, K. et al. The SARM1 Toll/Interleukin-1 Receptor Domain Possesses Intrinsic NAD(+) Cleavage Activity that Promotes Pathological Axonal Degeneration. Neuron 93, 1334–1343 e5 (2017).

41. Li, W.H. et al. Permeant fluorescent probes visualize the activation of SARM1 and uncover an anti-neurodegenerative drug candidate. Elife 10(2021).

42. Angeletti, C. et al. Programmed axon death executor SARM1 is a multi-functional NAD(P)ase with prominent base exchange activity, all regulated by physiological levels of NMN, NAD, NADP and other metabolites. bioRxiv (2021).

43. Essuman, K. et al. TIR Domain Proteins Are an Ancient Family of NAD(+)-Consuming Enzymes. Curr Biol 28, 421–430 e4 (2018).

44. Waller, T.J. & Collins, C.A. Multifaceted roles of SARM1 in axon degeneration and signaling. Front Cell Neurosci 16, 958900 (2022).

45. Li, Y. et al. Sarm1 activation produces cADPR to increase intra-axonal Ca++ and promote axon degeneration in PIPN. J Cell Biol 221(2022).

46. Geisler, S. et al. Prevention of vincristine-induced peripheral neuropathy by genetic deletion of SARM1 in mice. Brain 139, 3092–3108 (2016).

47. Loring, H.S. & Thompson, P.R. Emergence of SARM1 as a Potential Therapeutic Target for Wallerian-type Diseases. Cell Chem Biol 27, 1–13 (2020).

48. Gould, S.A. et al. Sarm1 haploinsufficiency or low expression levels after antisense oligonucleotides delay programmed axon degeneration. Cell Rep 37, 110108 (2021).

49. Loreto, A., et al. SARM1 activation induces reversible mitochondrial dysfunction and can be prevented in human neurons by antisense oligonucleotides. in bioRxiv (2024).

50. Loreto, A. et al. SARM1 activation induces reversible mitochondrial dysfunction and can be prevented in human neurons by antisense oligonucleotides. Neurobiol Dis 213, 106986 (2025).

51. Geisler, S. et al. Gene therapy targeting SARM1 blocks pathological axon degeneration in mice. J Exp Med 216, 294–303 (2019).

52. Loring, H.S., Parelkar, S.S., Mondal, S. & Thompson, P.R. Identification of the first noncompetitive SARM1 inhibitors. Bioorg Med Chem 28, 115644 (2020).

53. Hughes, R.O. et al. Small Molecule SARM1 Inhibitors Recapitulate the SARM1−/− Phenotype and Allow Recovery of a Metastable Pool of Axons Fated to Degenerate. Cell Reports 34, 108588 (2021).

54. Bosanac, T. et al. Pharmacological SARM1 inhibition protects axon structure and function in paclitaxel-induced peripheral neuropathy. Brain 144, 3226–3238 (2021).

55. Bratkowski, M. et al. Uncompetitive, adduct-forming SARM1 inhibitors are neuroprotective in preclinical models of nerve injury and disease. Neuron 110, 3711–3726 e16 (2022).

56. Feldman, H.C. et al. Selective inhibitors of SARM1 targeting an allosteric cysteine in the autoregulatory ARM domain. Proc Natl Acad Sci U S A 119, e2208457119 (2022).

57. Shi, Y. et al. Structural basis of SARM1 activation, substrate recognition, and inhibition by small molecules. Mol Cell 82, 1643–1659 e10 (2022).

58. Chen, J. & Li, H. Characterization of Novel SARM1 Inhibitors for the Treatment of Chemotherapy-Induced Peripheral Neuropathy. Biomedicines 12(2024).

59. Tang, Q. & Yin, H. Pyridine-based small molecule inhibitors of SARM1 alleviate cell death caused by NADase activity. Chem Commun (Camb) (2024).

60. Leahey, R., et al. SARM1 orthosteric base exchange inhibitors cause subinhibitory SARM1 activation. bioRxiv (2024).

61. Wenbin, Z., et al. SARM1 Activation Promotes Axonal Degeneration Via a Two-Step Liquid-to-Solid Phase Transition. bioRxiv (2024).

62. Mani, A. et al. SARM1 base-exchange inhibitors induce SARM1 activation and neurodegeneration at low doses. bioRxiv (2025).

63. Giroud, M., et al. Discovery of a Potent SARM1 Base-Exchange Inhibitor with In Vivo Efficacy. J Med Chem (2025).

64. Green, S.A. et al. Optimization of Brain Penetrant SARM1 Orthosteric Inhibitors and Discovery of Their Paradoxical Subinhibitory Activation. ACS Med Chem Lett 16, 1147–1154 (2025).

65. Reardon, H.T., et al. Base exchange inhibitors of SARM1 form mononucleotide adducts and activate SARM1 in vivo. bioRxiv (2025).

66. Leahey, R.R. et al. Therapeutic safety implications of SARM1 active site inhibitors: subinhibitory concentrations cause neurodegeneration. npj Drug Discovery 2(2025).

67. Sinclair, I. et al. Novel Acoustic Loading of a Mass Spectrometer: Toward Next-Generation High-Throughput MS Screening. J Lab Autom 21, 19–26 (2016).

68. Sinclair, I. et al. Acoustic Mist Ionization Platform for Direct and Contactless Ultrahigh-Throughput Mass Spectrometry Analysis of Liquid Samples. Anal Chem 91, 3790–3794 (2019).

69. Wernevik, J. et al. A Fully Integrated Assay Panel for Early Drug Metabolism and Pharmacokinetics Profiling. Assay Drug Dev Technol 18, 157–179 (2020).

70. Irwin, J.J. et al. An Aggregation Advisor for Ligand Discovery. J Med Chem 58, 7076–87 (2015).

71. Aarhus, R., Graeff, R.M., Dickey, D.M., Walseth, T.F. & Lee, H.C. ADP-ribosyl cyclase and CD38 catalyze the synthesis of a calcium-mobilizing metabolite from NADP. J Biol Chem 270, 30327–33 (1995).

72. Sauve, A.A., Munshi, C., Lee, H.C. & Schramm, V.L. The reaction mechanism for CD38. A single intermediate is responsible for cyclization, hydrolysis, and base-exchange chemistries. Biochemistry 37, 13239–49 (1998).

73. Preugschat, F., Tomberlin, G.H. & Porter, D.J. The base exchange reaction of NAD+ glycohydrolase: identification of novel heterocyclic alternative substrates. Arch Biochem Biophys 479, 114–20 (2008).

74. Hou, Y.N. et al. A conformation-specific nanobody targeting the nicotinamide mononucleotide-activated state of SARM1. Nat Commun 13, 7898 (2022).

75. Zhang, W. et al. SARM1 activation promotes axonal degeneration via a two-step phase transition. Nat Chem Biol (2025).

76. Moskowitz, P.F., Smith, R., Pickett, J., Frankfurter, A. & Oblinger, M.M. Expression of the class III beta-tubulin gene during axonal regeneration of rat dorsal root ganglion neurons. J Neurosci Res 34, 129–34 (1993).

77. Zhang, Y. et al. Rapid single-step induction of functional neurons from human pluripotent stem cells. Neuron 78, 785–98 (2013).

78. Huang, K. et al. Base-Exchange Enabling the Visualization of SARM1 Activities in Sciatic Nerve-Injured Mice. ACS Sens 8, 767–773 (2023).

79. Buonvicino, D. et al. Identification of the Nicotinamide Salvage Pathway as a New Toxification Route for Antimetabolites. Cell Chem Biol 25, 471–482 e7 (2018).

80. Loreto, A. et al. Neurotoxin-mediated potent activation of the axon degeneration regulator SARM1. Elife 10(2021).

81. Sverkeli, L.J., Hayat, F., Migaud, M.E. & Ziegler, M. Enzymatic and Chemical Syntheses of Vacor Analogs of Nicotinamide Riboside, NMN and NAD. Biomolecules 11(2021).

82. Seltzer, Z., Dubner, R. & Shir, Y. A novel behavioral model of neuropathic pain disorders produced in rats by partial sciatic nerve injury. Pain 43, 205–218 (1990).

83. Haffner, C.D., et al. Discovery, Synthesis, and Biological Evaluation of Thiazoloquin(az)olin(on)es as Potent CD38 Inhibitors. J Med Chem 58, 3548–71 (2015).

84. Zhu, W.J. et al. Gap junction intercellular communications regulates activation of SARM1 and protects against axonal degeneration. Cell Death Dis 16, 13 (2025).

85. Sporny, M. et al. Structural basis for SARM1 inhibition and activation under energetic stress. Elife 9(2020).

86. Klein, T. et al. Structural and dynamic insights into the energetics of activation loop rearrangement in FGFR1 kinase. Nat Commun 6, 7877 (2015).

87. Cowieson, N.P. et al. Beamline B21: high-throughput small-angle X-ray scattering at Diamond Light Source. J Synchrotron Radiat 27, 1438–1446 (2020).

88. Sader, K., Matadeen, R., Castro Hartmann, P., Halsan, T. & Schlichten, C. Industrial cryo-EM facility setup and management. Acta Crystallogr D Struct Biol 76, 313–325 (2020).

89. Dickerson, J.L., Leahy, E., Peet, M.J., Naydenova, K. & Russo, C.J. Accurate magnification determination for cryoEM using gold. Ultramicroscopy 256, 113883 (2024).

90. Punjani, A., Rubinstein, J.L., Fleet, D.J. & Brubaker, M.A. cryoSPARC: algorithms for rapid unsupervised cryo-EM structure determination. Nat Methods 14, 290–296 (2017).

