## Supplementary Information for "The Rise and Fall of SARM1 Base-Exchange Inhibitors"

**Supplementary Figures**

Supplementary Fig. 1: HTS assay development and validation

Supplementary Fig. 2: Hit counter screening and triage

Supplementary Fig. 3: Hydrogen-Deuterium Exchange Mass Spectrometry data

Supplementary Fig. 4: Structural studies using cryo-EM

Supplementary Fig. 5: *In vitro* cell assays

Supplementary Fig. 6: Development of an in-house iPSC-derived NGN2 cell model

Supplementary Fig. 7: Additional activation phenomena in cell assays

**Supplementary Tables**

Supplementary Table 1 | Data reproduced from Table 1 in the main text including uncertainty estimates

Supplementary Table 2 | Data reproduced from Table 1 in the main text including uncertainty estimates

Supplementary Table 3 | Data reproduced from Table 1 in the main text including uncertainty estimates

Supplementary Table 4 | Cryo-EM data collection, refinement and validation statistics

Supplementary Table 5 | Activity for selected compounds at key $NAD^+$-related enzymes

Supplementary Table 6 | CYP isoform inhibition & basicity of selected compounds

Supplementary Table 7 | Data reproduced from Table 2 in the main text including uncertainty estimates

Supplementary Table 8 | Secondary Pharmacology Screen for selected compounds

**Supplementary Methods**

Synthesis of inhibitors and adducts.

$^1$H NMR of adduct **6**, urea **14** and amide **16.**

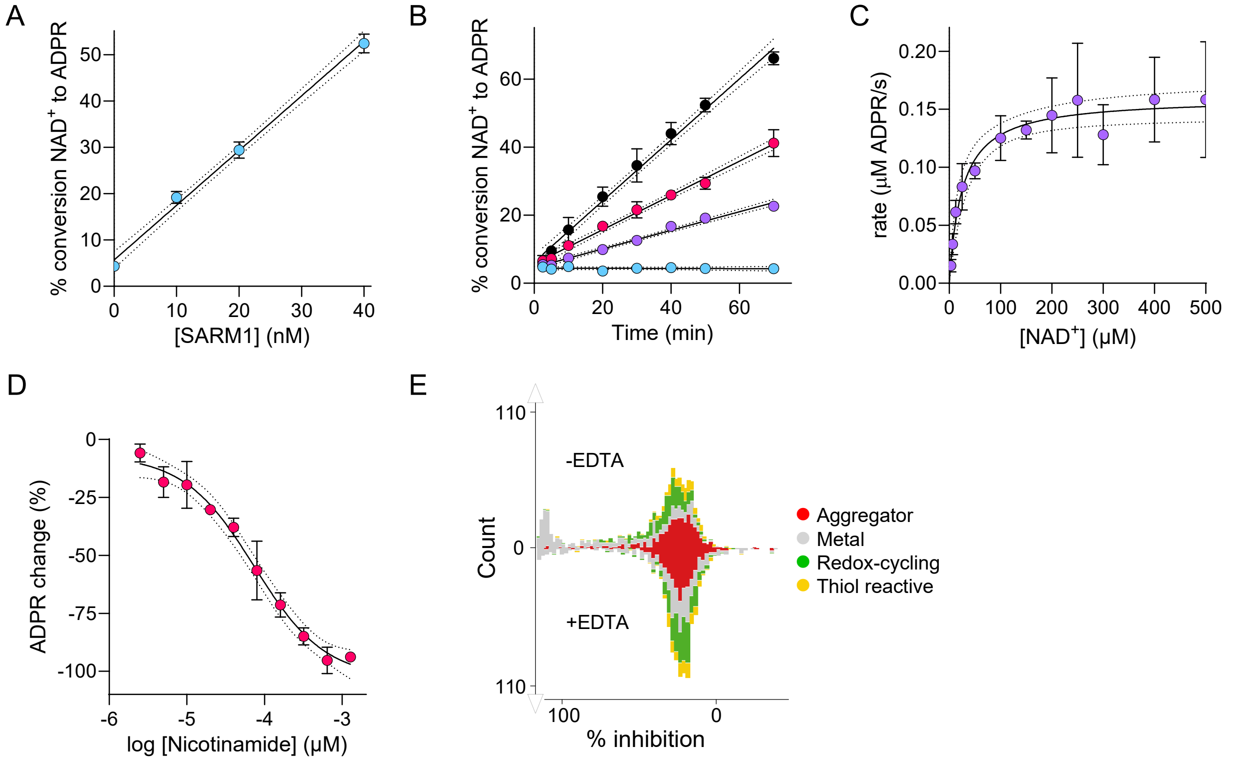


Supplementary Fig. 1 | HTS assay development and validation. A) Measured conversion of 25 µM $NAD^+$ substrate to ADPR after 50 minutes incubation as a function of recombinant human $SARM1^{\Delta 1\text{-}27}$ measured using AMI-MS (●). The solid line represents the best fit straight line to the data with a slope of 1.18 and $r^2$=0.99, while the dashed lines represent the 95% confidence bands. B) Measured conversion of 25 µM $NAD^+$ substrate to ADPR as a function of time and titrating $SARM1^{\Delta 1\text{-}27}$ concentrations at 0 (●), 10 (●), 20 (●), and 40 nM (●). The solid and dashed lines are as described for 1A with slopes of 0.00, 0.27, 0.50 and 0.90 with $r^2$ values of 0.01, 0.97, 0.98, and 0.98, respectively. C) Rate of enzymatic conversion as a function of substrate concentration. The solid line represents the best fit of data to the Michaeli-Menten model with $v_{max}$=0.16 µM ADPR/s, $K_m$=25 µM and $r^2$=0.79. D) Concentration response curve demonstrating product inhibition of $SARM1^{\Delta 1\text{-}27}$ activity by nicotinamide, with symbols denoting the average and standard deviation of three technical replicates (n=3). The solid line represents the best fits to a four-parameter logistic equation in GraphPad Prism, with an $IC_{50}$ of 74 µM, a Hill slope of -0.96 and $r^2$=0.97. E) Results from tests of assay sensitivity to a set of compounds with known unwanted mechanism of interference (uMOI), also referred to in the literature as pan-assay interference compounds (PAINS), in the absence and presence of 25 µM EDTA. Data are plotted as histograms with number of appearances (y-axis) versus the extent of apparent inhibition (x-axis).

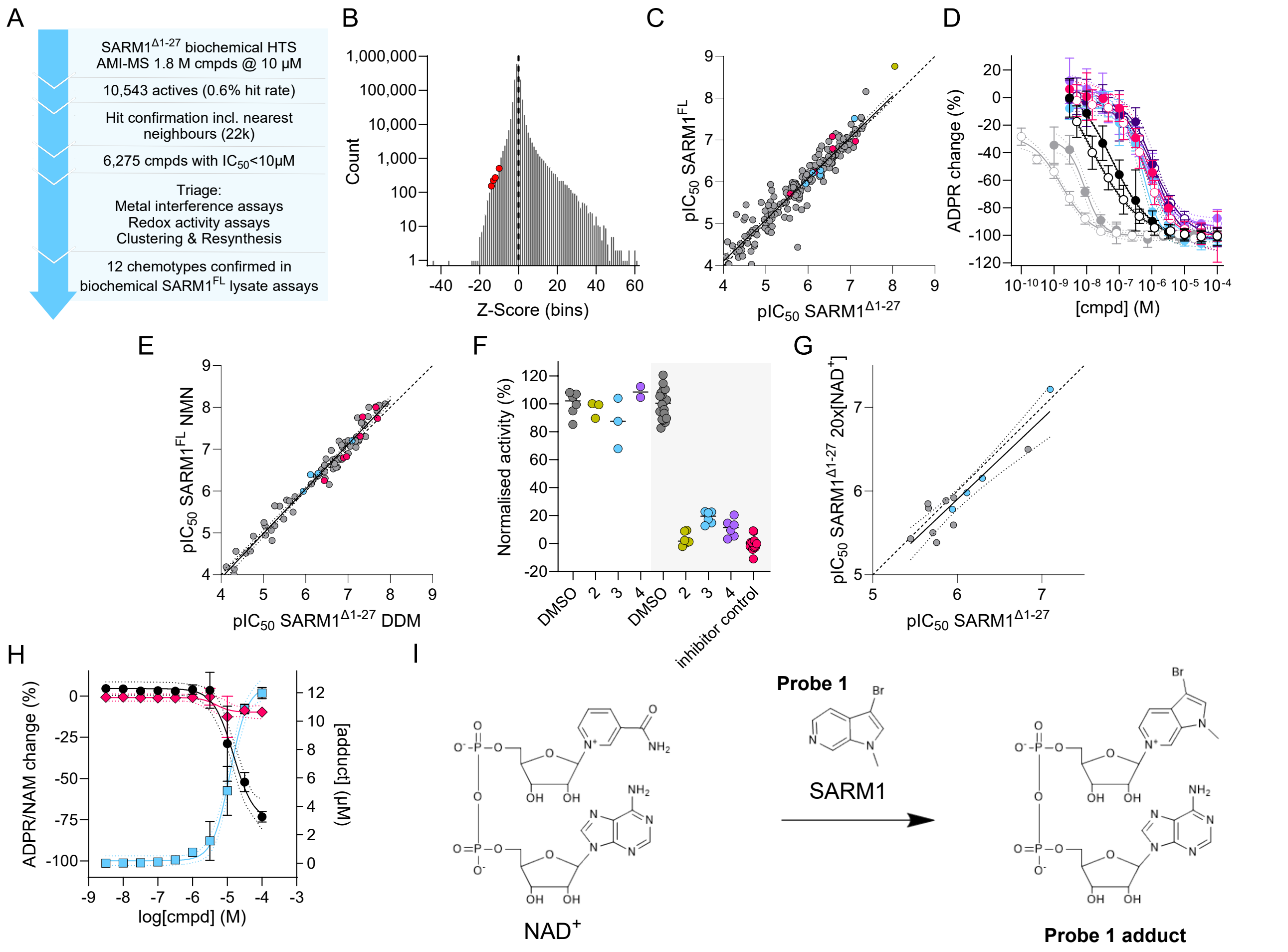


Supplementary Fig. 2 | Hit counter screening and triage. A) Summary of the HTS and hit annotation process resulting in the identification of multiple chemical clusters including Cmpd **1**, the activity and structure-activity relationship of which is described in the main text. B) Illustration of the HTS output in the form of counts of binned Z-scores and highlighting the location of four out of five hits in Table 1 of the main manuscript. The fifth hit was not part of the original screen as a pure compound, but it was identified through follow-up studies as a degradation product in one of the active solutions. C) Correlation of observed potencies between two complementary biochemical assay formats, with the HTS assay based on truncated recombinant SARM1$^{\Delta1-27}$ on the x-axis and cell lysates overexpressing SARM1$^{FL}$ on the y-axis. Symbol colours represent different BEXi chemotypes (●, Table 1 hits; ●, Table 2 compounds including competitor compounds; ●, Adduct **6**; ●, Other BEXi). The solid line represents the best-fit straight line with a slope of 0.99, a correlation coefficient $r^2$=0.90 and with the 95% confidence bands shown by dotted lines. The dashed line represents equality. D) Reproduction of concentration response data from the two biochemical assays as depicted in Fig. 1 of the main text, included here to show standard deviations from data obtained at independent test occasions (N). The dotted lines represent the 95% confidence bands from the four-parameter logistic equation fit in GraphPad Prism. E) Correlation of observed potencies in biochemical assay formats based on SARM1$^{\Delta1-27}$ activated by DDM, as employed for the HTS campaign (x-axis), versus cell lysates overexpressing SARM1$^{FL}$ activated by physiologically relevant NMN (y-axis) with lines and symbols as denoted in 2B. The slope of the solid line is 1.06 with a correlation coefficient $r^2$=0.97. F) Data from jump dilution experiments for compounds **2** (●), **3** (●), and **4** (●), with DMSO vehicle (●) and inhibitor controls (●). While the rapid 500-fold dilution of pre-incubated samples at 10 µM compound concentration (non-greyed area to the left of the graph) largely recovered full activity, non-diluted samples (greyed area to the right) at 10 µM compound concentration were nearly fully inhibited. The solid lines represent sample means with standard deviations. G) Correlation of observed potencies in biochemical assay formats based on truncated recombinant SARM1$^{\Delta1-27}$ at two different NAD$^+$ concentrations, with the concentration close to $K_m$ applied in the HTS campaign plotted on the x-axis and a 20-fold higher concentration on the y-axis. Lines and symbols are as denoted in 2B. The slope of the solid line is 0.96, with a correlation coefficient $r^2$=0.95. H) The concentration dependent redirection of SARM1$^{\Delta1-27}$ enzymatic activity by a base-exchange substrate (**Probe 1** – outlined in 2H) is differently reflected measuring the release of NAM (◆) and formation of ADPR (●), respectively. The inhibition of ADPR formation coincided with the appearance of the base-exchange adduct (■), while NAM formation is less affected. I) Outline of the expected base-exchange reaction for **Probe 1** in the active site of SARM1, whereby the parent base-exchange substrate replaces nicotinamide to form a new dinucleotide species.

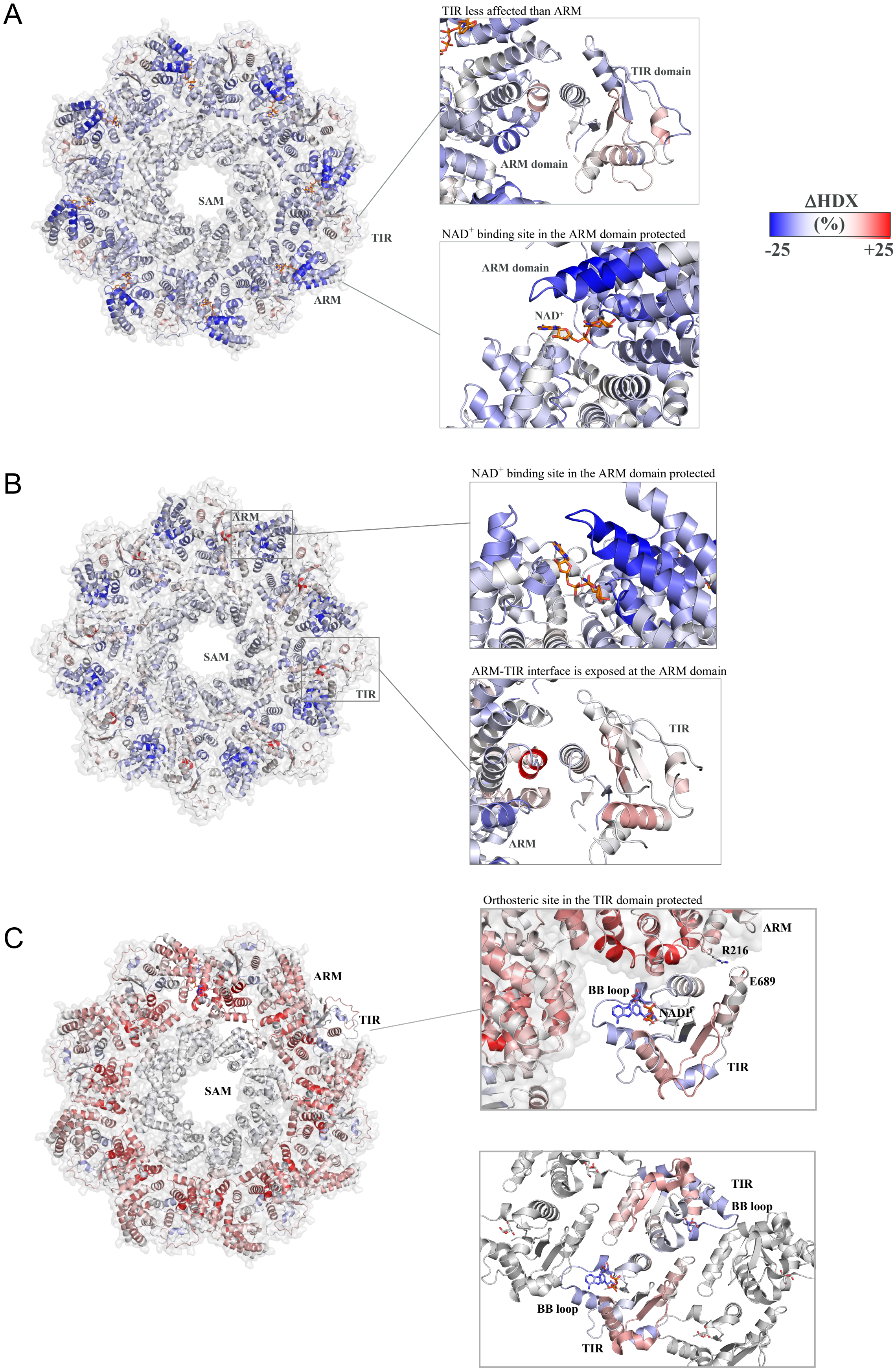
A
TIR less affected than ARM
TIR domain
ARM domain
SAM
TIR
ARM
NAD+ binding site in the ARM domain protected
ARM domain
NAD+
ΔHDX
(%)
-25
+25
B
ARM
NAD+ binding site in the ARM domain protected
SAM
TIR
ARM-TIR interface is exposed at the ARM domain
TIR
ARM
C
Orthosteric site in the TIR domain protected
ARM
R216
E689
BB loop
NADP
TIR
ARM
TIR
SAM
TIR
BB loop
BB loop
TIR

Supplementary Fig. 3 | Hydrogen-Deuterium Exchange Mass Spectrometry data. $NAD^+$, NMN and BEXi adduct **6** leads to different deuterium labelling patterns of the SARM1 octamer mapped on a composite structure based on the full length cryo-EM structure (7CM6), taking the BB-loop from the crystal structure (6O0Q) and NADP (in stick representation) from overlay with the crystal structure of a CD38-NADP complex (2I65). A) Incubation of SARM1 with 5 mM $NAD^+$ shows increased protection from hydrogen-deuterium exchange in the $NAD^+$ binding site in the ARM domain (inset, bottom), but moderate effects on the TIR domain (inset, top). B) Incubation with 5 mM NMN showed increased protection in the $NAD^+$ binding site in the ARM domain (inset, top), but two helices in the ARM-TIR interface were more exposed compared to the ligand-free state (inset, bottom). C) Incubation with 0.3 mM **6** resulted in a dramatic increase in exposure of the entire ARM domain, including the $NAD^+$ binding site. In contrast, the active site in the TIR domain was more protected (inset, top), suggesting binding to the active site, dimerization of TIR or both. The bottom inset shows the crystallographic dimer of TIR domains (6O0Q). Symmetry related molecules in the crystal structure are shown in grey. NADP (in stick representation) is derived from overlay with the crystal structure of a CD38-NADP complex (2I65). The scale bar at the top right shows the level of protection/deprotection in the maps.

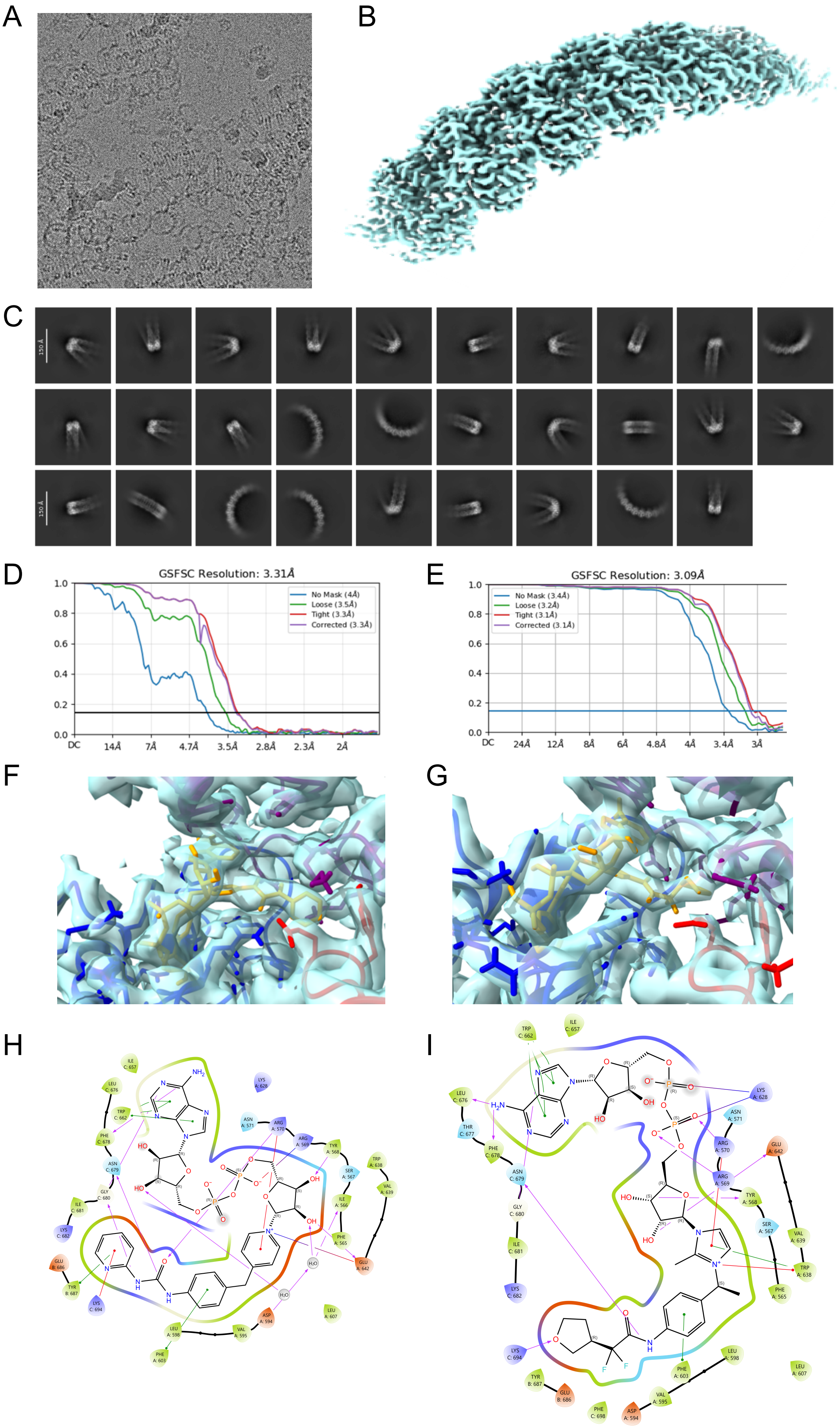
A
B
C
150 Å
D
GSFSC Resolution: 3.31Å
No Mask (4Å)
Loose (3.5Å)
Tight (3.3Å)
Corrected (3.3Å)
1.0 0.8 0.6 0.4 0.2 0.0
DC 14Å 7Å 4.7Å 3.5Å 2.8Å 2.3Å 2Å
E
GSFSC Resolution: 3.09Å
No Mask (3.4Å)
Loose (3.2Å)
Tight (3.1Å)
Corrected (3.1Å)
1.0 0.8 0.6 0.4 0.2 0.0
DC 24Å 12Å 8Å 6Å 4.8Å 4Å 3.4Å 3Å
F
G
H
I

Supplementary Fig. 4 | Structural studies using cryo-EM. A) Electron micrograph of TIR filaments formed in the presence of adduct **6** (the image is 269 nm x 269 nm). B) 3D Reconstruction of the TIR filaments in the presence of adduct **6**. (C) 2D class averages of TIR adduct **6** filament particles. D) and E) Fourier shell correlation plots for adduct **6** and **17**, respectively. F) and G) 3D maps and TIR/adduct model of the orthosteric site in the presence of adduct **6** and **17**, respectively. H) and I) Interaction maps showing interactions between adduct **6** and **17**, respectively, with the three TIR monomers that make up the binding site. Residues are coloured according to charge type and labelled according to whether they come from monomer $TIR^A$, $TIR^B$ or $TIR^C$ (see the main text and Fig. 2C for the nomenclature). It is noted that the highlighted interactions are assigned in Maestro.

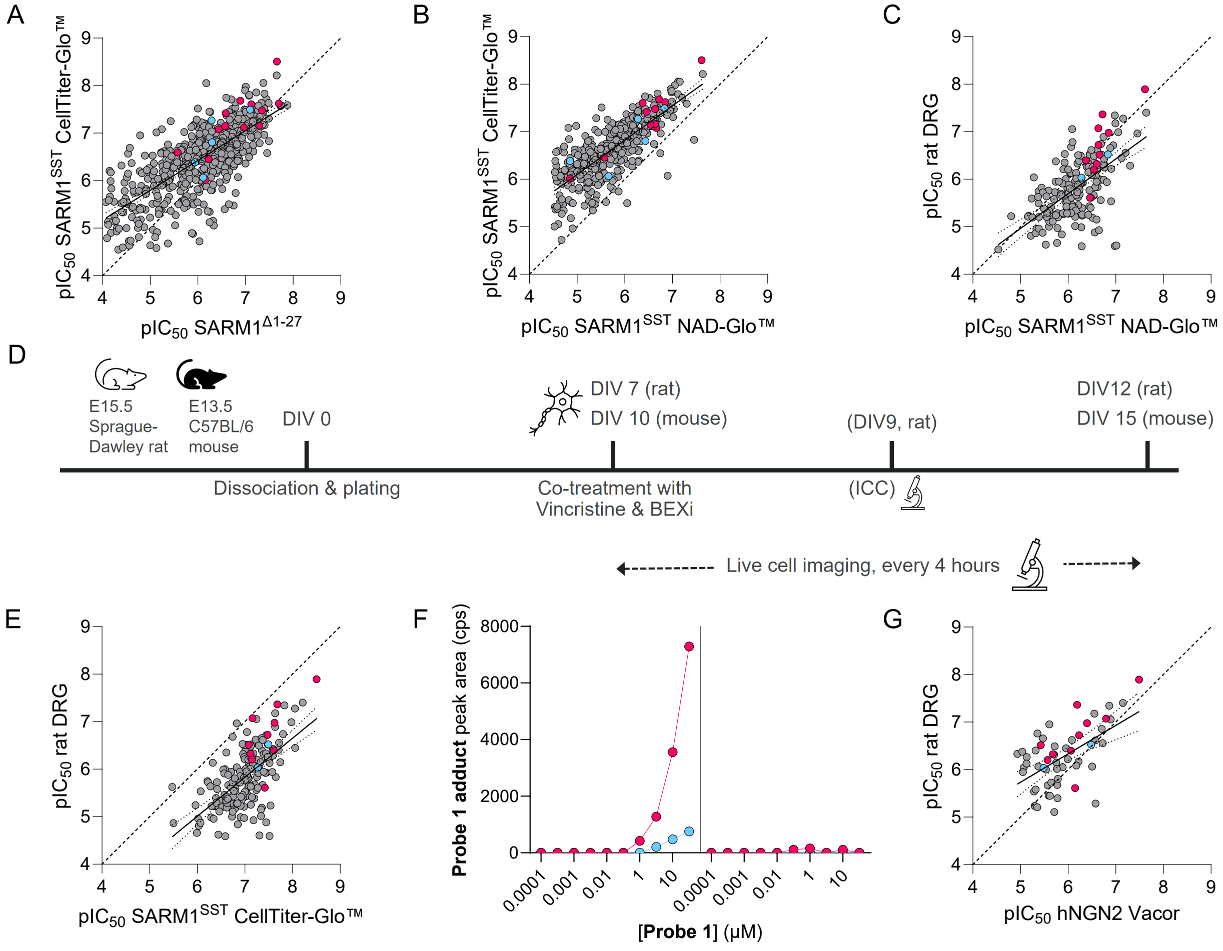


Supplementary Fig. 5 | *In vitro* cell assays. A) Correlation of observed potencies between the key SAR-driving biochemical assay (x-axis) and a cell viability assay in HEK293 cells overexpressing SARM1$^{SST}$ (y-axis). Symbol colours represent different BEXi chemotypes (●, Table 1 hits; ●, Table 2 compounds including competitor compounds; ●, Other BEXi). The solid line represents the best-fit straight line with a slope of 0.63, a correlation coefficient $r^2$=0.52 and with the 95% confidence bands shown by dotted lines. The dashed line represents equality. B) Correlation of observed potencies between the two overexpressed SARM1$^{SST}$ assays in HEK293 cells, with the intracellular NAD$^+$ readout (x-axis) versus the cell viability readout (y-axis). Lines and symbols are as denoted in 5A. The slope of the solid line is 0.72, with a correlation coefficient $r^2$=0.62. C) Correlation of potencies in the rat DRG assay (y-axis) with the intracellular NAD$^+$ measurements in HEK293 cells overexpressing SARM1$^{SST}$ (x-axis). Lines and symbols are as denoted in 5A. The slope of the solid line is 0.73, with a correlation coefficient $r^2$=0.37. D) Schematic illustration of the mouse and rat DRG assay procedures, starting with isolation and plating of DRG from the embryos of Sprague-Dawley rats or C57/BL mice, and subsequent culture in the plates before co-treatment with vincristine and BEXi and reading out through immunocytochemistry or live cell imaging. E) Correlation of potencies in the rat DRG assay (y-axis) with cell viability measurements in HEK293 cells overexpressing SARM1$^{SST}$ (x-axis). Lines and symbols are as denoted in 5A. The slope of the solid line is 0.82, with a correlation coefficient $r^2$=0.38. F) Measured formation of **Probe 1 adduct** in HEK293 cells expressing constitutively active SARM$^{SST}$ (left) versus non-expressing parental cells (right). LC/MS was used for the analysis of cell lysates following incubation with **Probe 1** for 4 (●) and 24h (●), respectively. G) Correlation of potencies in the rat DRG assay (y-axis) with those in the dox-inducible human NGN2 cell line expressing SARM1$^{FL}$ (x-axis). Lines and symbols are as denoted in 5A. The slope of the solid line is 0.60, with a correlation coefficient $r^2$=0.33.

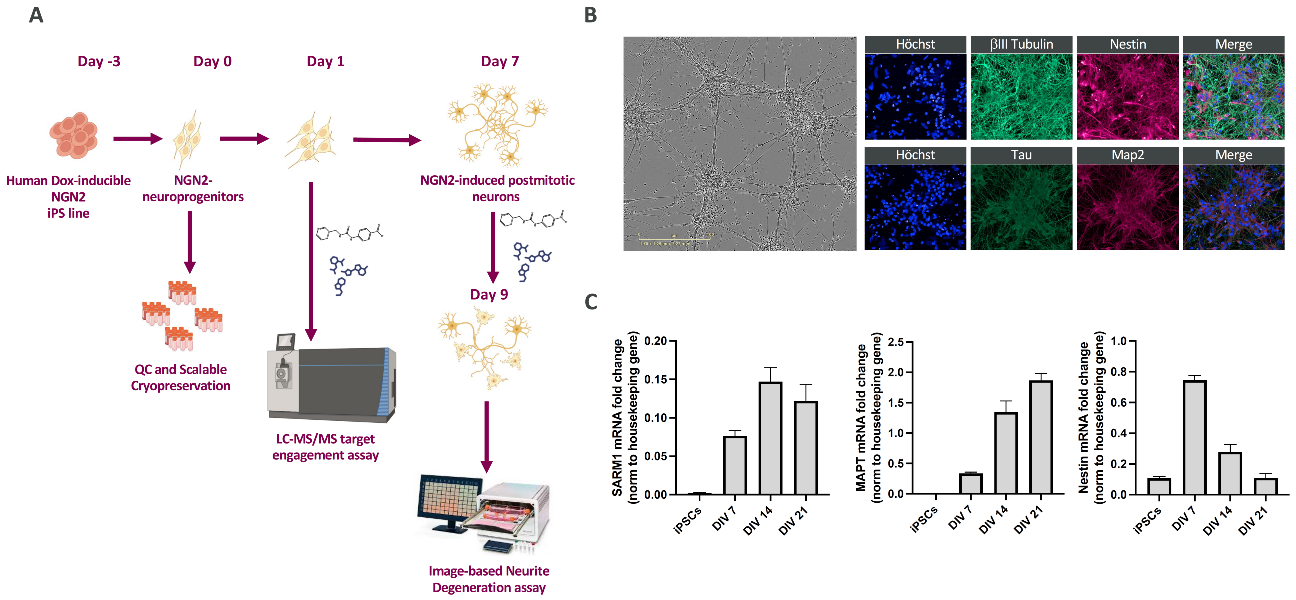


Supplementary Fig. 6 | Differentiation and characterization of NGN2-induced human neurons. A) Schematic of human iPSC-derived NGN2 neurons differentiation workflow. Doxycycline-inducible NGN2 iPSCs were induced into NGN2-neuroprogenitors (day 0), which were either cryopreserved in large batches or further differentiated into postmitotic neurons (day 7). Cells were treated with Vacor to induce SARM1 activation in presence of SARM1 BEXi (or DMSO as control) either at day 1 in a LC-MS/MS–based target engagement assays or at day 7 prior to perform an image-based neurite degeneration assay at day 9. B) Representative brightfield image of NGN2-induced neurons at day 7 showing extensive neurite networks (left side, scale bar: 400 µm). Right side shows immunofluorescence staining part of the QC panel confirming neuronal identity: Hoechst (blue), βIII-tubulin (green), and Nestin (magenta) (top row); Hoechst (blue), MAPT/Tau (green), and MAP2 (magenta) (bottom row). C) qRT–PCR analysis of neuronal differentiation. SARM1 and MAPT/Tau mRNA levels increased over time, whereas Nestin expression peaked at day 7 and declined with neuronal maturation. Data are mean ± SEM, normalized to housekeeping genes.

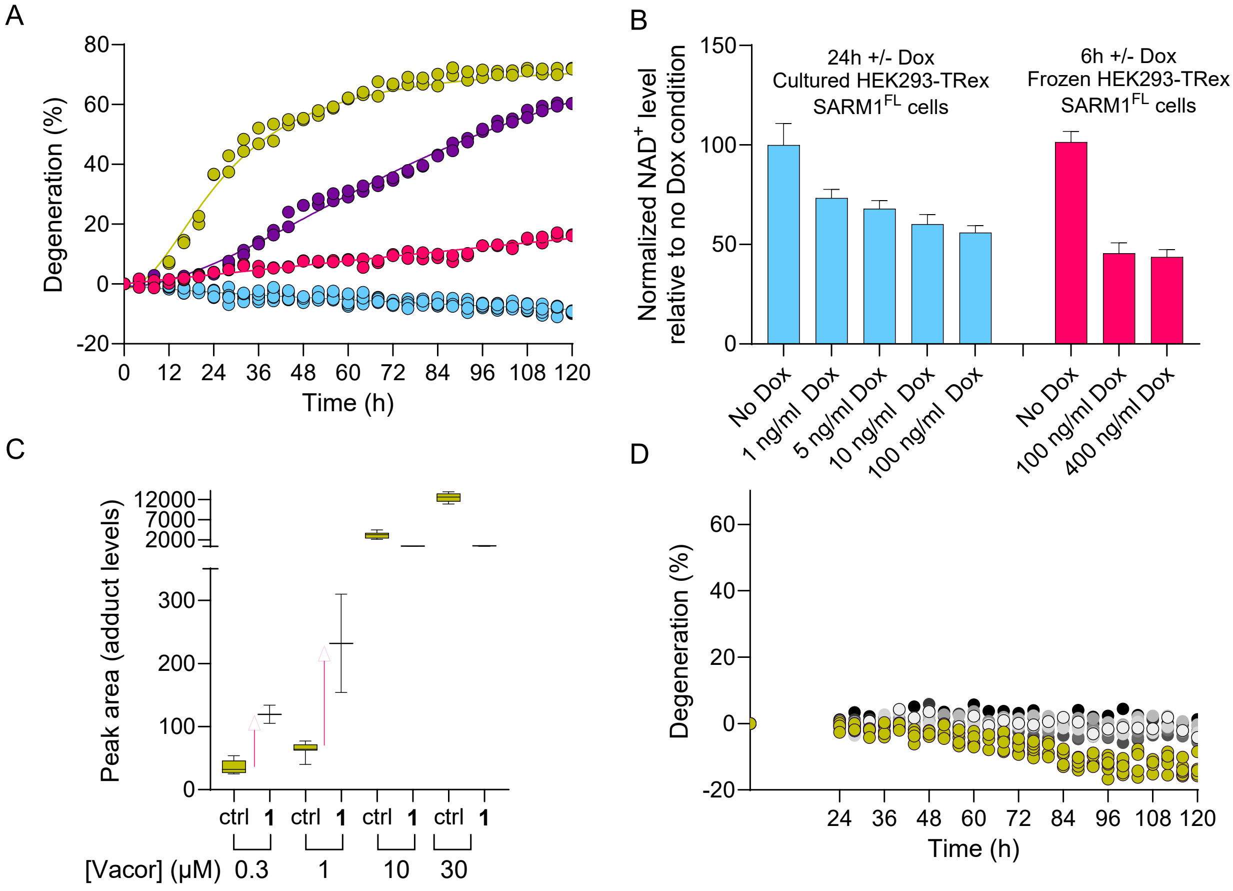


Supplementary Fig. 7 | Additional activatory effects in cell assays. A) Protective effects of heterozygous (+/-; ●) and homozygous (-/-; ●) knockout of SARM1 in mouse DRGs vs wildtype control (+/+; ●) treated with vincristine, with each symbol representing a separate technical replicate (n=2). Data are also included from an untreated wild-type control (●; n=6). B) Doxycycline titration in stably transfected HEK293 with TRex SARM1$^{FL}$ measuring NAD$^{+}$ levels. Data are shown from two independent experiments titrating different doxycycline ranges. Data are shown as the mean and standard deviations of 11-14 technical replicates. C) Reproduction of the data from Fig. 3E in the main text, where the human iPSC NGN2 cells are titrated with Vacor in the absence and presence of 10 µM **1**. The arrows denote how the SARM1 base-exchange activity is increased at the two lower doses of Vacor when **1** is present (red upward pointing arrows). D) Time course of neurite degeneration in rat DRG neurons treated with compound **14** over 120 h. The symbols denote DMSO control (●) and concentrations from 0.0014 (●) to 10 µM (●). Each technical replicate is shown (n=2; control n=6).

### Supplementary Tables

**Supplementary Table 1 | Data reproduced from Table 1 in main text including uncertainty estimates and data from the SARM1$^{FL}$ biochemical assay**

| Compound | SARM1$^{\Delta 1\text{-}27}$ biochemical $pIC_{50}$ | SARM1$^{FL}$ biochemical $pIC_{50}$ | SARM1$^{SST}$ cell $pIC_{50}$ | Neurite protection rat DRG[a] $pIC_{50}$ |
|---|---|---|---|---|
| **1** | 7.10 ± 0.17 (232) | 7.51 ± 0.14 (22) | 6.83 ± 0.14 (51) | 6.52 ± 0.12 (86) |
| **2** | 6.08, 6.15 (2) | 6.22 ± 0.12 (21) | 5.66 ± 0.20 (4) | <4 (1) |
| **3** | 6.30 ± 0.14 (6) | 6.41, 6.43 (2)[b] | 6.44 ± 0.26 (3) | <4, <4 (2) |
| **4** | 5.91, 5.98 (2) | 5.96 ± 0.04 (3) | 4.85 (1) | ND |
| **5** | 6.28, 6.30 (2) | 6.41 ± 0.15 (3)[b] | 6.28 ± 0.14 (3) | 6.04 (1) |

Data are provided as means and standard deviations of $pIC_{50}$ ($-\log_{10}[IC_{50}]$) values from the indicated number (N) of independent test occasions. All values are provided when N is less than three.

[a] Measured at Pharmaron, Beijing.

[b] Data are provided from the assay using NMN-activated SARM$^{FL}$ when the N numbers are greater than in the DDM-activated assay. As shown in Supplementary Fig. 2D these assays give equivalent data for BEXi.

ND: not determined.

**See separate excel files for Supplementary Tables 2 and 3.**

**Supplementary Table 4 | Cryo-EM data collection, refinement and validation statistics**

| | Adduct **6**<br>(EMDB-xxxx)<br>(PDB xxxx) | Adduct **17**<br>(EMDB-xxxx)<br>(PDB xxxx) |
|---|---|---|
| **Data collection and processing** | | |
| Magnification | 120k | 120k |
| Voltage (kV) | 300 | 300 |
| Electron exposure (e–/Å$^2$) | 43 | 60 |
| Defocus range (μm) | -2.2 to -0.8 | -2.4 to -1.6 |
| Pixel size (Å) | 0.656 | 0.642 |
| Symmetry imposed | No | No |
| Initial particle images (no.) | 2,535,468 | 7,667,327 |
| Final particle images (no.) | 110,926 | 2,643,954 |
| Map resolution (Å)<br>FSC threshold | 3.31 (0.143) | 3.09 (0.143) |
| Map resolution range (Å) | 2.4-3.6 | 3.0 |
| **Refinement** | | |
| Initial model used (PDB code) | n/a | |
| Model resolution (Å)<br>FSC threshold | 3.2 (0.143) | 3.0 |
| Model resolution range (Å) | 2.9-3.4 | 2.5-3.0 |
| Map sharpening *B* factor (Å$^2$) | -80 | -196 |
| Model composition<br>Non-hydrogen atoms<br>Protein residues<br>Ligands | 9112, 552, 4 | 9088,552,4 |
| *B* factors (Å$^2$)<br>Protein<br>Ligand | | |
| R.m.s. deviations<br>Bond lengths (Å)<br>Bond angles (°) | 0.004,0.575 | 0.007,0.741 |
| Validation<br>MolProbity score<br>Clashscore<br>Poor rotamers (%) | 1.85,5.23,2.52 | 2.68,12.30,3.83 |
| Ramachandran plot<br>Favored (%)<br>Allowed (%)<br>Disallowed (%) | 96.14,3.86,0 | 86.58,10.11,3.31 |

### Supplementary Table 5 | Activity for selected compounds at key NAD$^+$-related enzymes

| | Compound | SARM1 IC$_{50}$ µM | NAMPT[a] IC$_{50}$ µM | CD38[a] IC$_{50}$ µM | NMNAT1[a] IC$_{50}$ µM |
|---|---|---|---|---|---|
| **1** | 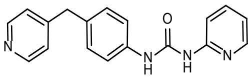 | 0.080 | 46 | >100 | >100 |
| **3** | 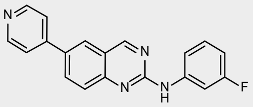 | 0.50 | 59 | >100 | >100 |
| **4** | 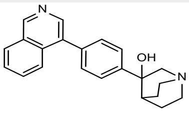 | 1.1 | >100 | ND | >100 |
| **5** | 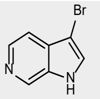 | 0.52 | >100 | >100 | >100 |
| **9** | 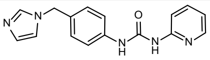 | 0.26 | ND | >100 | ND |
| **10** | 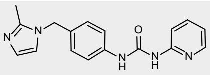 | 0.075 | ND | >100 | ND |
| **Vacor** | 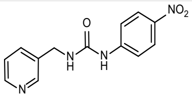 | >100 | 25 | ND | >100 |

[a] Enzyme inhibition measured at Pharmaron, Beijing.
ND: not determined.

### Supplementary Table 6 | Cyp isoform[a] inhibition & basicity of selected compounds

| | Compound | CYP3A4 Midazolam $IC_{50}$ (μM) | CYP2D6 Bufuralol $IC_{50}$ (μM) | CYP2C9 Diclofenac $IC_{50}$ (μM) | CYP2C19 S-Meph $IC_{50}$ (μM) | CYP1A2 Phenacetin $IC_{50}$ (μM) | pKa (most basic)[b] |
|---|---|---|---|---|---|---|---|
| **1** | 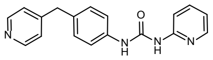 | 1.4 | <0.10 | 0.67 | 1.0 | 0.33 | 5.7 |
| **3** | 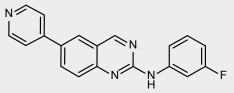 | 4.2 | 2.8 | 0.65 | 1.3 | 4.1 | ND |
| **4** | 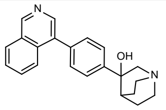 | >30 | 18 | >30 | >30 | >30 | ND |
| **5** | 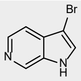 | >30 | 0.90 | >30 | >30 | 13.4 | ND |
| **8** | 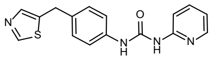 | 2.8 | 0.41 | 0.66 | 1.5 | 0.61 | ND |
| **9** | 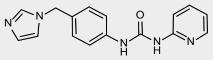 | 1.5 | 0.16 | 0.54 | 1.5 | 0.32 | ND |
| **10** | 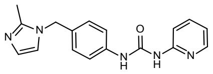 | >30 | >30 | >30 | >30 | >30 | ND |
| **11** | 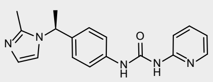 | >30 | 21 | >30 | >30 | >30 | ND |
| **13** | 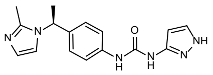 | >30 | >30 | >30 | >30 | >30 | 7.6 |
| **14** | 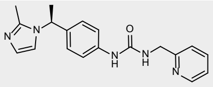 | >30 | >30 | >30 | >30 | >30 | ND |
| **15** | 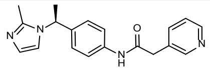 | >30 | >30 | >30 | >30 | >30 | 7.7 |
| **DSRM-3716** | 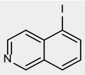 | 9.5 | 8.7 | 1.2 | 11 | 0.86 | ND |
| **NB-2** | 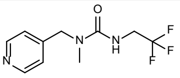 | >30 | >30 | >30 | >30 | 20 | ND |
| **NB-3** | 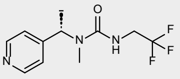 | 4.7 | >30 | >30 | >30 | 12 | 5.2 |
| **NB-7** | 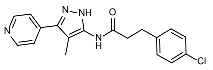 | 2.0 | 1.1 | 7.3 | 0.45 | 0.19 | 4.9 |

[a] Inhibition of Cyp enzymes in Human Liver microsomes, measured at Pharmaron, Beijing; The respective substrates are indicated for each isoform.
[b] $pK_a$ of most basic centre experimentally determined at pION, East Sussex, UK.

**Supplementary Table 7 | Data reproduced from Table 2 in the main text including uncertainty estimates**

| Compound | SARM1$^{\Delta1\text{-}27}$ biochem $pIC_{50}$ | SARM1$^{FL}$ biochem $pIC_{50}$ | SARM1$^{SST}$ cell $pIC_{50}$ | NGN2 cell $pIC_{50}$ | Neurite protection rat DRG[a] $pIC_{50}$ |
|---|---|---|---|---|---|
| **1** | 7.10±0.17 (232) | 7.51±0.14 (22) | 6.83±0.14 (51) | 6.48±0.26 (54) | 6.52±0.12 (86) |
| **7** | 5.58 (1) | <5, <4 (2)[b] | <4.5 (1) | <5.0, <5.0 (2) | ND |
| **8** | 6.07, 6.24 (2) | 6.23 (1) | 4.85 (1) | <5.0, <5.0 (2) | <4.0, <4.0 (2) |
| **9** | 6.58±0.23 (4) | 7.09 (1) | 6.54±0.10 (3) | 5.57 (1) | 6.20, 6.21 (2) |
| **10** | 7.12±0.07 (3) | 6.97 (1) | 6.52, 6.23 (2) | 6.06 (1) | 6.40 (1) |
| **11** | 7.35±0.14 (8) | 7.77 (1)[b] | 6.64±0.22 (6) | 6.23±0.10 (5) | 6.72±0.24 (5) |
| **12** | 6.23 (1) | 6.53, 6.24 (2)[b] | 5.57 (1) | 5.42 (1) | ND |
| **13** | 7.70±0.06 (3) | 7.74 (1)[b] | 6.85, 6.85 (2) | 6.51, 6.28 (2) | 6.98±0.21 (4) |
| **14** | 7.29±0.04 (3) | 7.31±0.06 (4)[b] | 6.64±0.36 (3) | 6.80±0.24 (3) | 7.07, 7.08 (2) |
| **15** | 6.43±0.16 (7) | ND | 6.19±0.16 (7) | ND | 5.96, 6.06 (2) |
| **16** | 6.90, 7.04 (2) | 6.79, 6.85 (2)[b] | 6.63, 6.58 (2) | 5.68±0.48 (3) | 6.32 (1) |
| **DSRM-3716** | 6.50, 6.66 (2) | 6.74, 6.84 (2) | 6.46±0.09 (3) | 6.15±0.25 (5) | 5.61 (1) |
| **NB-2** | 6.44±0.44 (3) | 6.23, 6.27 (2)[b] | 6.66 (1) | 5.58, 5.28 (2) | 6.51 (1) |
| **NB-3** | 6.89±0.24 (4) | 6.80 (1) | 6.72 (1) | 6.24, 6.13 (2) | 7.37 (1) |
| **NB-7** | 7.57, 7.75 (2) | 8.00±0.17 (85) | 7.62±0.19 (4) | 7.49±0.12 (55) | 7.90 (1) |

Data are provided as means and standard deviations of $pIC_{50}$ values from the indicated number (N) of independent test occasions. All values are provided when N is less than three.

[a] Measured at Pharmaron, Beijing.

[b] Data are provided from the assay using NMN-activated SARM$^{FL}$ when the N numbers are greater than in the DDM-activated assay. As shown in Supplementary Fig. 2D these assays give equivalent data for BEXi.

ND: not determined

**Supplementary Table 8 | Secondary Pharmacology Screen for selected compounds**

| Target | Compound | | |
|---|---|---|---|
| | 1 | 14 | 16 |
| PDE3A $IC_{50}$ (μM) | >100 | >100 | >100 |
| PDE4D $IC_{50}$ (μM) | >30 | >100 | >100 |
| ACHE $IC_{50}$ (μM) | >100 | >100 | >100 |
| ROCK2 $IC_{50}$ (μM) | >100 | >100 | >100 |
| KDR $IC_{50}$ (μM) | 3.61 | >100 | >100 |
| INSR $IC_{50}$ (μM) | >100 | >100 | >100 |
| A1 $IC_{50}$ (μM) | 31.7 | >100 | >100 |
| a1A $IC_{50}$ (μM) | 50.6 | >100 | >100 |
| a2A $IC_{50}$ (μM) | >100 | >100 | >100 |
| b1 $IC_{50}$ (μM) | >100 | >100 | >100 |
| b1 $EC_{50}$ (μM) | >100 | >100 | >100 |
| b1 $IC_{50}$ (μM) | 2.61 | >100 | >100 |
| D1 $IC_{50}$ (μM) | 66 | >100 | >100 |
| 5HT2B $EC_{50}$ (μM) | >100 | | |
| 5HT2B $IC_{50}$ (μM) | 20.6 | | |
| CB1 $EC_{50}$ (μM) | 46.8 | >100 | >100 |
| CB1 $IC_{50}$ (μM) | 12.3 | >100 | >100 |
| M2 $IC_{50}$ (μM) | >100 | 37.9 | 48.2 |
| OPRm1 $IC_{50}$ (μM) | 38.2 | 27.2 | 96.3 |
| GABA TBPS $IC_{50}$ (μM) | 20.8 | | |
| NAchA1 $IC_{50}$ (μM) | >100 | >100 | >100 |
| NAchA4 $IC_{50}$ (μM) | >100 | >100 | >30 |
| NMDA AGO $IC_{50}$ (μM) | >100 | | >100 |
| CaV-L VER $IC_{50}$ (μM) | >100 | >100 | >100 |
| DAT $IC_{50}$ (μM) | 5.84 | >100 | >100 |
| NET $IC_{50}$ (μM) | 16.1 | >100 | >100 |
| GABAA BNZ $IC_{50}$ (μM) | 2.16 | >100 | >100 |
| PDE10A2 $IC_{50}$ (μM) | >100 | >100 | >100 |
| PDE6 $IC_{50}$ (μM) | 17.1 | >100 | >100 |
| CatS $IC_{50}$ (μM) | >100 | >100 | >100 |
| PDPK1 $IC_{50}$ (μM) | >30 | >100 | >100 |
| MMP2 $IC_{50}$ (μM) | >100 | >100 | |
| Na+/K+ ATPase $IC_{50}$ (μM) | >100 | >100 | >100 |
| TXA2 sy $IC_{50}$ (μM) | 2.16 | 46.8 | >100 |
| cKIT $IC_{50}$ (μM) | | >100 | >100 |
| ROCK1 $IC_{50}$ (μM) | >100 | >100 | >100 |
| ALK4 $IC_{50}$ (μM) | >100 | >30 | >100 |
| FGFR1 $IC_{50}$ (μM) | >100 | >100 | >100 |
| EGFR kinase $IC_{50}$ (μM) | >30 | >100 | >100 |
| AurKA $IC_{50}$ (μM) | >100 | >100 | >100 |
| src $IC_{50}$ (μM) | 4.79 | >100 | >100 |
| MAP3K7 $IC_{50}$ (μM) | >30 | >100 | >100 |
| GSK3b $IC_{50}$ (μM) | >100 | >100 | >100 |
| TRKA $IC_{50}$ (μM) | 18.4 | >100 | >100 |
| A2A $IC_{50}$ (μM) | 6.34 | 18.5 | >100 |
| a1B $IC_{50}$ (μM) | >100 | >100 | >100 |
| a1B $EC_{50}$ (μM) | >100 | >100 | >100 |
| a1B $IC_{50}$ (μM) | >100 | >100 | >100 |

| | | | |
|---|---|---|---|
| a2C $IC_{50}$ (μM) | >100 | 57.6 | >100 |
| A2A $EC_{50}$ (μM) | >100 | | |
| A2A $IC_{50}$ (μM) | >100 | | |
| b2 $IC_{50}$ (μM) | >100 | >100 | >100 |
| D3 $IC_{50}$ (μM) | >100 | >100 | >100 |
| D3 $EC_{50}$ (μM) | >100 | >100 | >100 |
| D3 $IC_{50}$ (μM) | >100 | >100 | >100 |
| Ang2 AT1 $IC_{50}$ (μM) | 36.4 | >100 | >100 |
| Ghre $IC_{50}$ (μM) | >100 | >100 | >100 |
| H1 $IC_{50}$ (μM) | >100 | >100 | >100 |
| EtA $IC_{50}$ (μM) | >100 | >100 | >100 |
| H2 $IC_{50}$ (μM) | >100 | >100 | >100 |
| H2 $EC_{50}$ (μM) | >100 | >100 | >100 |
| H2 $IC_{50}$ (μM) | >30 | >100 | >100 |
| M1 $IC_{50}$ (μM) | 76.9 | 17 | 61.9 |
| DOP2 $IC_{50}$ (μM) | 47.6 | >100 | >100 |
| M5 $IC_{50}$ (μM) | >100 | 65.9 | >100 |
| SST4 $IC_{50}$ (μM) | >100 | >100 | >100 |
| MR2 $IC_{50}$ (μM) | >100 | >100 | >100 |
| 5-HT1A $IC_{50}$ (μM) | 14.8 | >100 | >100 |
| 5-HT1D $IC_{50}$ (μM) | >100 | >100 | >100 |
| 5-HT2C $IC_{50}$ (μM) | >100 | >100 | >100 |
| 5-HT2C $EC_{50}$ (μM) | >100 | >100 | >100 |
| 5-HT2C $IC_{50}$ (μM) | >100 | >100 | >100 |
| 5HT4 $IC_{50}$ (μM) | 82.3 | >100 | >100 |
| 5HT7 $IC_{50}$ (μM) | 44.8 | >100 | >100 |
| 5-HT7 $EC_{50}$ (μM) | >100 | >100 | >100 |
| 5-HT7 $IC_{50}$ (μM) | >100 | >100 | >100 |
| NK1 $IC_{50}$ (μM) | 47.5 | >100 | >100 |
| BK2 $IC_{50}$ (μM) | >100 | >100 | >100 |
| Sigma1 $IC_{50}$ (μM) | 67.4 | >30 | >100 |
| PPARgamma $IC_{50}$ (μM) | 29.1 | >100 | >100 |
| RARa $IC_{50}$ (μM) | >100 | >100 | >100 |
| GR $IC_{50}$ (μM) | 61 | >100 | >100 |
| CaV-L DIL $IC_{50}$ (μM) | | >100 | >100 |
| 5HT3 $IC_{50}$ (μM) | 63 | >100 | >100 |
| NAchA7 $IC_{50}$ (μM) | >100 | >100 | >100 |
| NMDA PCP $IC_{50}$ (μM) | >100 | >100 | >100 |
| GABAa1b2g2 AGO $IC_{50}$ (μM) | >100 | >100 | >100 |
| GlyR $IC_{50}$ (μM) | >100 | >100 | >100 |
| AT Gpig $IC_{50}$ (μM) | 7.41 | >100 | >100 |
| SET $IC_{50}$ (μM) | 9.94 | >100 | >100 |

### Compound Synthesis

Unless stated otherwise, starting materials were commercially available. All solvents and commercial reagents were of laboratory grade and were used as received.

### Abbreviations

DCM: dichloromethane; DIPEA: N,N-diisopropylethylamine; DMF: N,N-dimethylformamide; DMSO: dimethyl sulfoxide; Et2O: diethyl ether; EtOAc: ethyl acetate; FA: formic acid; HATU: hexafluorophosphate azabenzotriazole tetramethyl uronium; HCl: hydrogen chloride; HEPES: 4-(2-Hydroxyethyl)piperazine-1-ethanesulfonic acid, EtOH; ethanol; HPLC: high performance liquid chromatography; MeCN: acetonitrile; NMP: N-methylpyrrolidinone; Pd(dppf)Cl2: 1,1'-bis(diphenylphosphino)ferrocene]dichloropalladium(II); Pd/C: palladium on carbon: triphenylphosphine; [Rh(COD)Cl]2: chloro(1,5-cyclooctadiene)rhodium(I) dimer; SCF: supercritical fluid chromatography; TCFH: N′-tetramethylformamidinium hexafluorophosphate; TEA: triethylamine; TEA: triethylamine; TFA: trifluroacetic acid; THF: tetrahydrofuran.

### Analytical and Purification Methods

LCMS experiments were performed using a Shimadzu LCMS-2020 with electrospray ionization in positive ion detection mode with 20ADXR pump, SIL-20ACXR autosampler, CTO-20AC column oven, M20A PDA Detector and LCMS 2020 MS detector. LCMS was run in one of three set ups: method 1 [Halo C18 column (2.0 µm 3.0 x 30 mm) in combination with a gradient (5-100% B in 1.2 min.) of water and FA (0.1%) (A) and MeCN and FA (0.1%) (B) at a flow rate of 1.5 mL/min]; method 2 [Halo C18 column (2.0 µm 3.0 x 30 mm) in combination with a gradient (5-100% B in 1.2 min) of water and TFA (0.05%) (A) and MeCN and TFA (0.05%) (B) at a flow rate of 1.5 mL/min]; method 3 [Poroshell HPH C18 column (2.7 µm 3.0 x 50 mm) in combination with a gradient (10-95% B in 2 min.) of aqueous 46 mM ammonium carbonate/ammonia buffer at pH 10 (A) and MeCN (B) at a flow rate of 1.2 mL/min]. The Column Oven (CTO-20AC) temperature was 40 °C. The injection volume was 1 µL. PDA (SPD-M20A) detection was in the range λ (190–400) nm. The MS detector was configured with electrospray ionization as ionizable source; acquisition mode: Scan; nebulizing gas flow:1.5 L/min; drying gas flow:15 L/min; detector voltage: 0.95-1.25 kv; DL T: 250°C; heat block T: 250°C scan range: 90.00 – 900.00 *m/z*.

NMR Spectra were recorded on a Bruker AVANCE III HD 400 (400 MHz) or Bruker AVANCE NEO 400 (400 MHz) or Bruker AVANCE III 400 (400 MHz) or Bruker AVANCE II 300 (300 MHz) or Bruker AVANCE III 300 (300 MHz) or Bruker AVANCE III HD 300 (300 MHz).

Preparative reverse phase HPLC was performed on a Waters instrument (2545 or 2767 or 2489) fitted with a QDa or SQ Detector 2 ESCi mass spectrometers.

Preparative Chiral SFC was performed on a Waters instrument SFC (80 or 100 or 150 or 350) fitted with UV2489 (or mass spectrometer).

### Synthesis of hit compounds 1-5

The synthesis of Compounds **3** and **5** was previously reported in the literature: **3** (WO 2008/020203 A1 pyridinylquinazolinamine derivatives and their use as B-Raf inhibitors); **5** (WO 2020/081923 A1 inhibitors of SARM1 in combination with $NAD^+$ or a $NAD^+$ precursor and WO 2019/236884 A1 inhibitors of SARM1)

*1-(pyridin-2-yl)-3-(4-(pyridin-4-ylmethyl)phenyl)urea (**1**)*

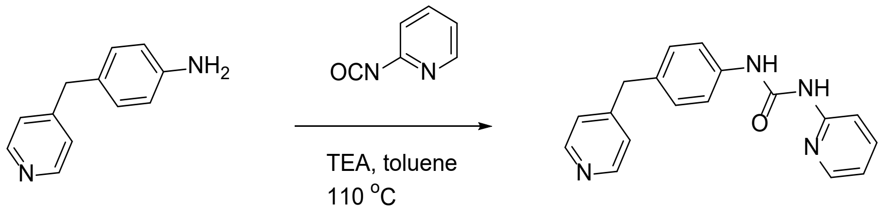


**1**

4-(Pyridin-4-ylmethyl)aniline (1.70 g, 9.22 mmol, 1.0 equiv) was added to a stirred mixture of 2-isocyanatopyridine (2.22 g, 18.48 mmol, 2.0 equiv) and TEA (1.87 g, 18.48 mmol, 2.0 equiv) in toluene (133 mL) at room temperature. The mixture was stirred at 110 °C for 1 hour. The solvent was removed under reduced pressure, and the residue was purified by flash silica chromatography, eluting with 0 to 85% EtOAc in petroleum ether to afford 1-(pyridin-2-yl)-3-[4-(pyridin-4-ylmethyl)phenyl]urea **1** (2.4 g, 42.8%) as a white solid. MS ($ES^+$, *m/z*): 305.20 $[M + H]^+$. $^1$H NMR (400 MHz, DMSO-*d6*) δ 10.46 (s, 1H), 9.41 (s, 1H), 8.49 – 8.42 (m, 2H), 8.28 – 8.24 (m, 1H), 7.76 – 7.72 (m, 1H), 7.52 – 7.42 (m, 3H), 7.25 – 7.15 (m, 4H), 7.02 – 6.99 (m, 1H), 3.93 (s, 2H).

*Synthesis of 3-(benzo[d][1,3]dioxol-5-yl)-N-(3-cyano-4-(pyridin-4-yl)phenyl)propanamide (**2**)*

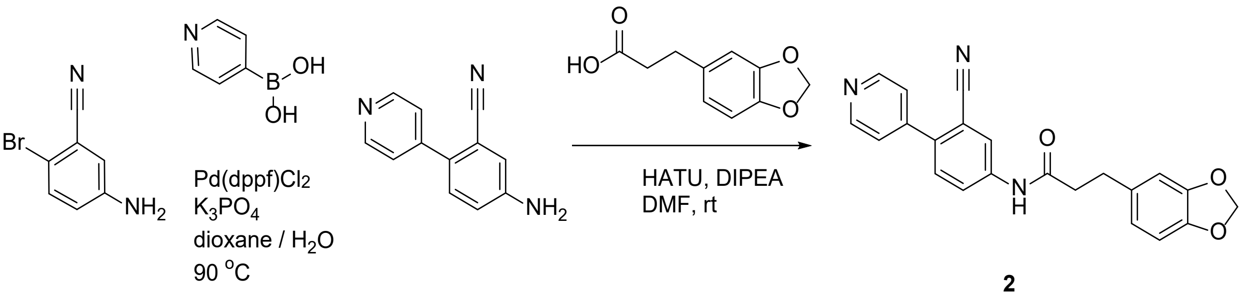


**2**

Step 1. $Pd(dppf)Cl_2$ (1.85 g, 2.54 mmol, 0.1 equiv) was added to a stirred degassed mixture of 5-amino-2-bromobenzonitrile (5 g, 25.38 mmol, 1.0 equiv), pyridin-4-yl boronic acid (3.74 g, 30.45 mmol, 1.2 equiv) and $K_3PO_4$ (10.77 g, 50.75 mmol, 1.6 equiv) in 1,4-dioxane (150 mL) and water (21.5 mL). The resulting mixture was stirred at 90 °C for 12 hours, diluted with EtOAc (200 mL), and washed with water (50 mL) and brine (50 mL). The organic layer was dried over $MgSO_4$, filtered and concentrated under reduced pressure. The crude product was purified by flash silica chromatography, eluting with 3 to 7% MeOH in DCM to afford 5-amino-2-(pyridin-4-yl)benzonitrile (4.0 g, 81%) as a yellow solid. $^1$H NMR (300 MHz, DMSO-*d6*) δ 5.91 (d, 2H), 6.94 (dd, 1H), 7.00 (d, 1H), 7.37 (d, 1H), 7.55 (bs, 2H), 8.78 (bs, 2H).

Step 2. 5-Amino-2-(pyridin-4-yl)benzonitrile (200 mg, 1.02 mmol, 1.0 equiv) was added to a stirred mixture of 3-(benzo[d][1,3]dioxol-5-yl)propanoic acid (219 mg, 1.13 mmol, 1.1 equiv), HATU (428 mg, 1.13 mmol, 1.1 equiv), and DIPEA (397 mg, 3.07 mmol, 2.5 equiv) in DMF (4 mL). The resulting mixture was stirred at temperature for 4 hours, diluted with EtOAc (10 mL), and washed with water (3 x 5 mL) and brine (10 mL). The organic layer was dried over $MgSO_4$, filtered and concentrated under reduced pressure. The crude product was purified by flash silica chromatography, eluting with

0 to 35% EtOAc in petroleum ether to afford 3-(benzo[d][1,3]dioxol-5-yl)-N-(3-cyano-4-(pyridin-4-yl)phenyl)propanamide **2** (0.024 g, 6.27%) as a white solid. MS (ES$^+$, *m/z*): 372.0 [M + H]$^+$. $^1$H NMR (300 MHz, DMSO-*d6*) δ 10.38 (s, 1H), 8.73 – 8.71 (m, 2H), 8.27 (d, J = 2.2 Hz, 1H), 7.89 (dd, J = 8.6, 2.3 Hz, 1H), 7.65 (d, J = 8.6 Hz, 1H), 7.61 – 7.59 (m, 2H), 6.85 – 6.80 (m, 2H), 6.70 (dd, J = 7.9, 1.7 Hz, 1H), 5.96 (s, 2H), 2.86 (t, J = 7.5 Hz, 2H), 2.65 (t, J = 7.5 Hz, 2H).

*Synthesis of (S)-3-(4-(isoquinolin-4-yl)phenyl)quinuclidin-3-ol (**4**)*

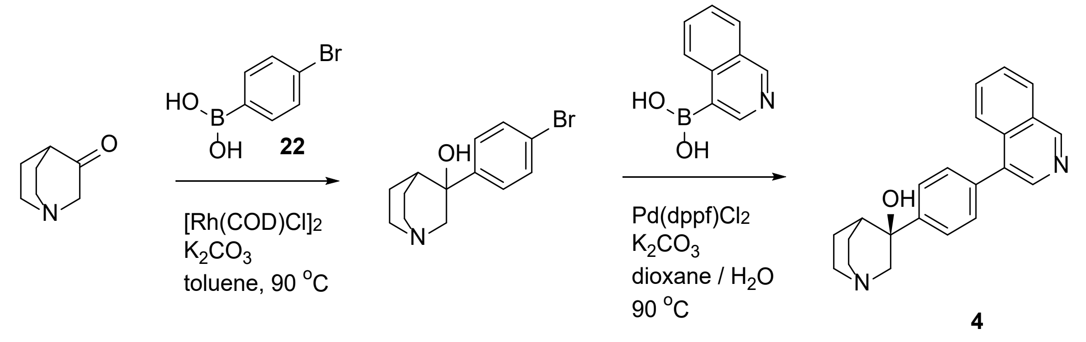


Step 1. Quinuclidin-3-one (700 mg, 5.59 mmol, 1.0 equiv) was added to a stirred mixture of (4-bromophenyl)boronic acid (1.18 g, 5.87 mmol, 1.05 equiv), [Rh(COD)Cl]2 (276 mg, 0.56 mmol, 0.1 equiv) and $K_2CO_3$ (2.32 g, 16.78 mmol, 3.0 equiv) in toluene (10 mL) under one atmosphere of nitrogen. The resulting mixture was stirred at 90 °C for 18 hours, then concentrated under reduced pressure. The crude product was purified by flash C18 chromatography, eluting with 0 to 35% MeCN in water to afford 3-(4-bromophenyl)quinuclidin-3-ol (0.65 g, 41.2%) as a yellow solid. MS (ES$^+$, *m/z*): 282.0 [M + H]$^+$. $^1$H NMR (400 MHz, DMSO-*d6*) δ 1.10 – 1.46 (m, 4H), 1.87 (t, J = 3.4 Hz, 1H), 2.02 – 2.17 (m, 1H), 2.62 – 2.74 (m, 3H), 2.76 – 2.89 (m, 2H), 5.18 (s, 1H), 7.44 – 7.53 (m, 4H).

Step 2. 3-(4-Bromophenyl)quinuclidin-3-ol (205 mg, 0.73 mmol, 1.0 equiv) was added to a mixture of isoquinolin-4-ylboronic acid (151 mg, 0.87 mmol, 1.2 equiv), Pd(dppf)$Cl_2$ (53.2 mg, 0.07 mmol, 0.1 equiv) and $K_2CO_3$ (301 mg, 2.18 mmol, 3.0 equiv) in 1,4-dioxane (5 mL) and water (0.2 mL) under one atmosphere of nitrogen. The resulting mixture was stirred at 80 °C for 16 hours. The crude product was purified sequentally by flash C18 chromatography, eluting with 50 to 60% MeCN in water and by preparative HPLC (Column: XBridge Prep OBD C18, 30 × 150mm, 5 µm; Mobile Phase A: water (10 mM $NH_4HCO_3$ + 0.1% $NH_4OH$), Mobile Phase B: MeCN; Flow rate: 60 mL/min; Gradient: 23 B to 43 B in 7 min; 254 / 220 nm; RT: 6.25 min) to afford (S)-3-(4-(isoquinolin-4-yl)phenyl)quinuclidin-3-ol **4** (0.030 g, 12.53%) as a white solid. MS (ES$^+$, *m/z*): 331.0 [M + H]$^+$. $^1$H NMR (400 MHz, DMSO-*d6*) δ 1.37 (q, J = 5.6, 7.5 Hz, 2H), 1.42 – 1.53 (m, 1H), 2.02 (s, 1H), 2.18 (d, J = 10.5 Hz, 1H), 2.66 – 2.79 (m, 3H), 2.87 (t, J = 10.1 Hz, 1H), 2.95 (dd, J = 1.8, 14.4 Hz, 1H), 3.40 – 3.48 (m, 1H), 5.23 (s, 1H), 7.50 – 7.57 (m, 2H), 7.68 – 7.78 (m, 3H), 7.81 (ddd, J = 1.5, 6.7, 8.4 Hz, 1H), 7.88 (d, J = 8.4 Hz, 1H), 8.23 (d, J = 8.1 Hz, 1H), 8.45 (s, 1H), 9.34 (s, 1H).

**Synthesis of intermediate amines**

*4-((2-methyl-1H-imidazol-1-yl)methyl)aniline*

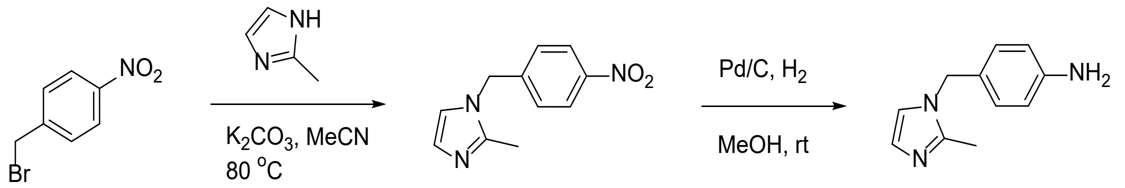


Step 1. 2-Methyl-1H-imidazole (3.04 g, 37.03 mmol, 5.0 equiv) was added to a solution of 1-(bromomethyl)-4-nitrobenzene (2 g, 9.26 mmol, 1 equiv) and $K_2CO_3$ (3.84 g, 27.77 mmol, 3.0 equiv) in MeCN (5 mL) at room temperature. The reaction mixture was stirred at 80 °C for 3 hours, cooled down and filtered. The filtrate was concentrated under reduced pressure and the crude product was purified by flash silica chromatography, eluting with 0 to 10% MeOH in DCM, to afford 2-methyl-1-(4-nitrobenzyl)-1H-imidazole **27** (1.7 g, 85%) as an orange solid. $^1$H NMR (300 MHz, MeOH-*d4*) δ 2.29 (s, 3H), 5.33 (s, 2H), 6.90 (d, J = 1.5 Hz, 1H), 7.10 (d, J = 1.4 Hz, 1H), 7.27 – 7.38 (m, 2H), 8.17 – 8.27 (m, 2H).

Step 2. 10% Pd/C (w/w) (0.78 g, 0.74 mmol, 0.1 equiv) was added to a solution of 2-methyl-1-(4-nitrobenzyl)-1H-imidazole (1.6 g, 7.37 mmol, 1.0 equiv) in MeOH (5 mL). The reaction mixture was stirred at room temperature under one atmosphere of hydrogen for 2 hours, then it was filtered through a pad of Celite®. The filtrate was concentrated under reduced pressure to afford 4-((2-methyl-1H-imidazol-1-yl)methyl)aniline (1.0 g, 72.5%). MS (ES$^+$, *m/z*): 188.3 [M + H]$^+$; 1H NMR (300 MHz, MeOH-d4) δ 2.29 (s, 3H), 4.97 (s, 2H), 6.65 – 6.72 (m, 2H), 6.86 – 6.96 (m, 4H), the exchangeable protons are not visible.

*Synthesis of intermediates (R) and (S) -4-(1-(2-methyl-1H-imidazol-1-yl)ethyl)aniline*

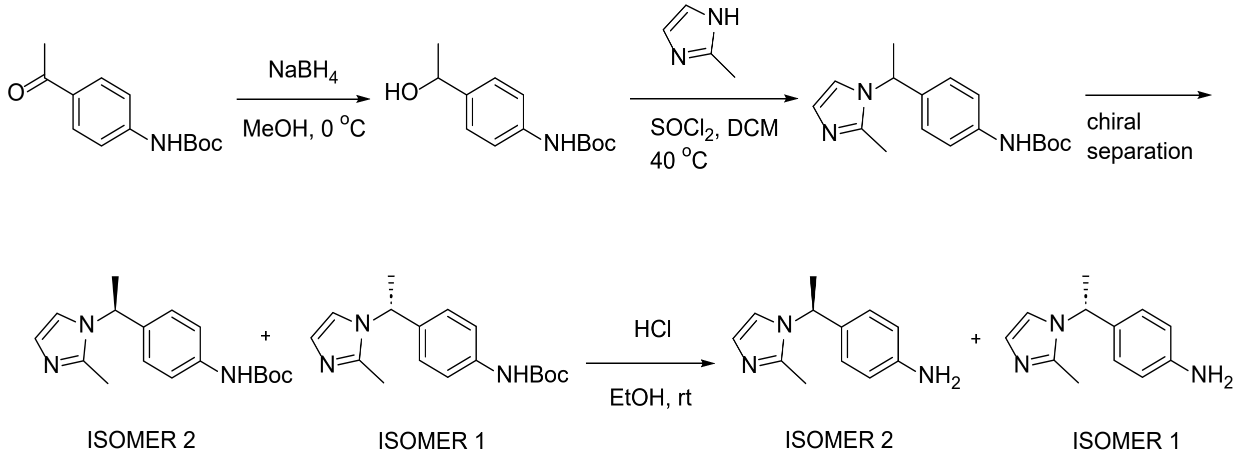


Step 1. $NaBH_4$ (3.54 g, 93.50 mmol, 2.0 equiv) was added to a solution of tert-butyl (4-acetylphenyl)carbamate (11 g, 46.75 mmol, 1.0 equiv) in MeOH (300 mL) at 0 °C. The reaction mixture was stirred at room temperature for 2 hours, then it was concentrated under reduced pressure. The residue was dissolved in DCM (200 mL) and washed with water (100 mL) and brine (100 mL). The organic layer was dried over $Na_2SO_4$, filtered and evaporated under reduced pressure. The crude product was purified by flash silica chromatography, eluting with 0 to 10% MeOH in DCM, to afford tert-butyl (4-(1-hydroxyethyl)phenyl)carbamate (10 g, 90%) as a white solid. MS (ES$^+$, *m/z*): 164.1

[M - OtBu]+; 1H NMR (300 MHz, CDCl3) δ 1.47 (d, J = 6.4 Hz, 3H), 1.52 (s, 9H), 4.86 (q, J = 6.4 Hz, 1H), 6.48 (s, 1H), 7.23 – 7.38 (m, 4H), OH proton is not visible.

Step 2. $SOCl_2$ (1.5 ml, 20.68 mmol, 1.6 equiv) was added dropwise to a solution of tert-butyl (4-(1-hydroxyethyl)phenyl)carbamate (3 g, 12.64 mmol, 1.0 equiv) and 2-methyl-1H-imidazole (7.5 g, 91.35 mmol, 7.2 equiv) in DCM (90 mL) at room temperature. The reaction mixture was stirred at 40 °C for 2 hours. The solvent was removed under reduced pressure and the crude product was purified by flash silica chromatography, eluting with 0 to 10% MeOH in DCM, to afford (rac)-tert-butyl (4-(1-(2-methyl-1H-imidazol-1-yl)ethyl)phenyl)carbamate (2.78 g, 68.5%) as a white solid. MS ($ES^+$, *m/z*): 302.1 [M +H]+; 1H NMR (400 MHz, $CDCl_3$) δ 1.51 (s, 9H), 1.79 (d, J = 7.1 Hz, 3H), 2.35 (s, 3H), 5.28 (q, J = 7.0 Hz, 1H), 6.48 (s, 1H), 6.94 – 7.03 (m, 4H), 7.29 – 7.36 (m, 2H).

Step 3. Chiral separation: (rac)-(4-(1-(2-Methyl-1H-imidazol-1-yl)ethyl)phenyl)carbamate (9.7 g, 32.18 mmol) was resolved by SFC (Column: Venusil Chiral OD-H, 21 x 250 mm, 5 µm; Mobile Phase A: $CO_2$, Mobile Phase B: MeOH (0.1% 2 M $NH_3$-MeOH); RT1: 8.4 min, RT2: 11.6 min, to afford:

ISOMER 1: (R)-(4-(1-(2-methyl-1H-imidazol-1-yl)ethyl)phenyl)carbamate (4.53 g, 46.7%), RT 8.47 min, as a colourless oil. MS ($ES^+$, *m/z*): 302.0 $[M + H]^+$; 1H NMR (300 MHz, DMSO-*d6*) δ 1.46 (s, 9H), 1.69 (d, J = 7.0 Hz, 3H), 2.20 (s, 3H), 3.18 (d, J = 5.0 Hz, 2H), 6.76 (d, J = 1.4 Hz, 1H), 7.03 – 7.12 (m, 2H), 7.21 (d, J = 1.4 Hz, 1H), 7.35 – 7.45 (m, 2H), ee: 99.56%.

ISOMER 2: (S)-(4-(1-(2-Methyl-1H-imidazol-1-yl)ethyl)phenyl)carbamate (3.95 g, 40.7%), RT 11.67 min, as a colourless oil. MS ($ES^+$, *m/z*): 302.0 $[M + H]^+$; 1H NMR (300 MHz, DMSO-*d6*) δ 1.46 (s, 9H), 1.69 (d, J = 7.0 Hz, 3H), 2.20 (s, 3H), 5.37 (q, J = 7.0 Hz, 1H), 6.76 (d, J = 1.4 Hz, 1H), 7.03 – 7.12 (m, 2H), 7.21 (d, J = 1.4 Hz, 1H), 7.36 – 7.44 (m, 2H), 9.35 (s, 1H), ee: 98.7%.

The absolute stereochemistry of isomer 2 was determined to be (*S*) by comparison with chirally pure material synthesised by an alternative procedure.

Step 4. (R) or (S)-(4-(1-(2-methyl-1H-imidazol-1-yl)ethyl)phenyl)carbamate was added to a 4 M HCl solution in EtOH. The reaction mixture was stirred at room temperature for 2 hours. The solvent was removed under reduced pressure to afford (R)-4-(1-(2-methyl-1H-imidazol-1-yl)ethyl)aniline or (S)-4-(1-(2-methyl-1H-imidazol-1-yl)ethyl)aniline as dihydrochloride salts and yellow solids. MS ($ES^+$, *m/z*): 202.0 $[M + H]^+$; 1H NMR (300 MHz, DMSO-*d6*) δ 1.84 (d, J = 6.9 Hz, 3H), 2.62 (d, J = 1.5 Hz, 3H), 5.82 (d, J = 7.4 Hz, 1H), 7.35 – 7.52 (m, 4H), 7.66 (d, J = 2.1 Hz, 1H), 7.88 (q, J = 2.3 Hz, 1H).

**Synthesis of ureas 7-14**

*1-(pyridin-2-yl)-3-(4-(pyridin-3-ylmethyl)phenyl)urea (7)*

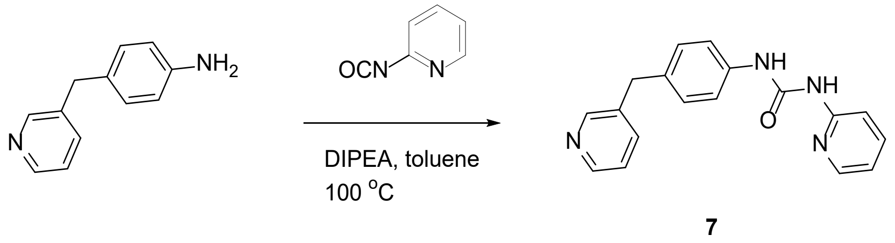


**7**

4-(pyridin-3-ylmethyl)aniline (100 mg, 0.54 mmol, 1.0 equiv) was added to a stirred mixture of 2-isocyanatopyridine (78 mg, 0.65 mmol, 1.0 equiv) and DIPEA (142 µl, 0.81 mmol, 1.25 equiv) in toluene (3 mL) at room temperature under one atmosphere of nitrogen. The mixture was stirred at 100 °C for 16 hours. The solvent was removed under reduced pressure, and the crude product was purified by preparative HPLC (Column: XBridge Shield RP18 OBD, 30 ×150mm, 5 μm; Mobile Phase A: water (0.05% $NH_4OH$), Mobile Phase B: MeCN; Flow rate: 60 mL/min; Gradient: 37 B to 45 B in 7 min; Detector: UV 254 / 220 nm; RT: 6.82 min) to afford 1-(pyridin-2-yl)-3-(4-(pyridin-3-ylmethyl)phenyl)urea **7** (0.064 g, 38.7%) as a white solid. MS ($ES^+$, *m/z*): 305.20 $[M + H]^+$. $^1$H NMR (300 MHz, DMSO-*d6*) δ 3.91 (s, 2H), 6.98 (ddd, J = 7.3, 5.0, 1.0 Hz, 1H), 7.13 – 7.23 (m, 2H), 7.29 (dd, J = 7.8, 4.7 Hz, 1H), 7.38 – 7.50 (m, 3H), 7.59 (dt, J = 7.8, 2.0 Hz, 1H), 7.72 (ddd, J = 8.9, 7.3, 2.0 Hz, 1H), 8.25 (dt, J = 4.9, 1.6 Hz, 1H), 8.39 (dd, J = 4.8, 1.6 Hz, 1H), 8.48 (d, J = 2.2 Hz, 1H), 9.38 (s, 1H), 10.42 (s, 1H).

*1-(pyridin-2-yl)-3-(4-(thiazol-5-ylmethyl)phenyl)urea (**8**)*

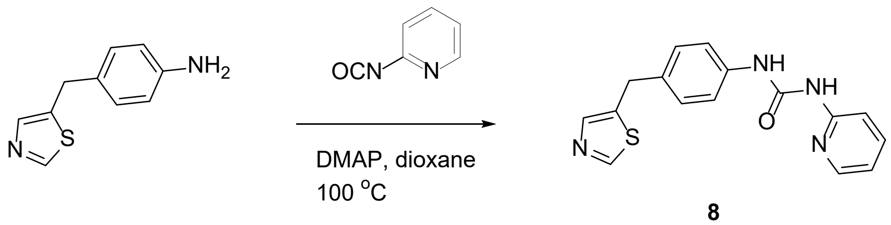


**8**

2-Isocyanatopyridine (25.3 mg, 0.21 mmol, 1.0 equiv) was added to a stirred mixture of 4-(thiazol-5-ylmethyl)aniline (40 mg, 0.21 mmol, 1.0 equiv) and DMAP (2.57 mg, 0.02 mmol, 0.10 equiv) in 1,4-dioxane (10 mL) at room temperature under one atmosphere of nitrogen. The mixture was stirred at 100 °C for 3 hours. The solvent was removed under reduced pressure, and the residue was purified by preparative HPLC (Column: XBridge Shield RP18 OBD, 30 × 150 mm, 5 μM; Mobile Phase A: water (10 mM $NH_4HCO_3$ + 0.1% $NH_4OH$), Mobile Phase B: MeCN; Flow rate: 60 mL/min; Gradient: 20 B to 30 B in 8 min; 254 / 220 nm; RT: 7.17 min) to afford 1-(pyridin-2-yl)-3-(4-(thiazol-5-ylmethyl)phenyl)urea **8** (0.027 g, 41.4%) as a white solid. MS ($ES^+$, *m/z*): 311.0 $[M + H]^+$. $^1$H NMR (400 MHz, DMSO-*d6*) δ 4.16 (s, 2H), 6.95-7.06 (m, 1H), 7.17 – 7.25 (m, 2H), 7.42 – 7.52 (m, 3H), 7.68 – 7.79 (m, 2H), 8.27 (ddd, J = 0.9, 1.9, 5.0 Hz, 1H), 8.93 (d, J = 0.8 Hz, 1H), 9.41 (s, 1H), 10.47 (s, 1H).

1-(4-((1H-imidazol-1-yl)methyl)phenyl)-3-(pyridin-2-yl)urea (**9**)

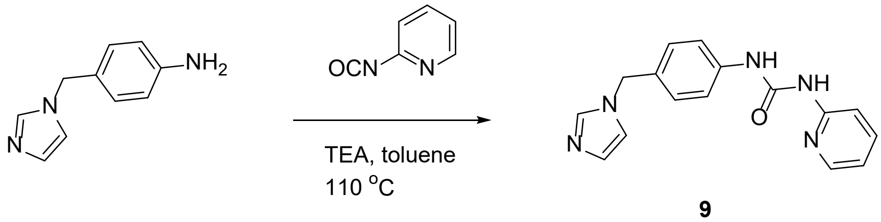


**9**

It was prepared from 4-((1H-imidazol-1-yl)methyl)aniline and 2-isocyanatopyridine following the same procedure described for 1 and isolated as a white solid after preparative HPLC purification (20.9 mg, 35.1%). MS ($ES^+$, *m/z*): 294.0 $[M + H]^+$; 1H NMR (400 MHz, DMSO-*d6*) δ 5.13 (s, 2H), 6.89 (s, 1H), 7.01 (dd, J = 5.1, 7.2 Hz, 1H), 7.17 (d, J = 1.2 Hz, 1H), 7.19 – 7.27 (m, 2H), 7.46 – 7.55 (m, 3H), 7.69 – 7.77 (m, 2H), 8.24 – 8.30 (m, 1H), 9.46 (s, 1H), 10.53 (s, 1H).

*1-(4-((2-methyl-1H-imidazol-1-yl)methyl)phenyl)-3-(pyridin-2-yl)urea (**10**)*

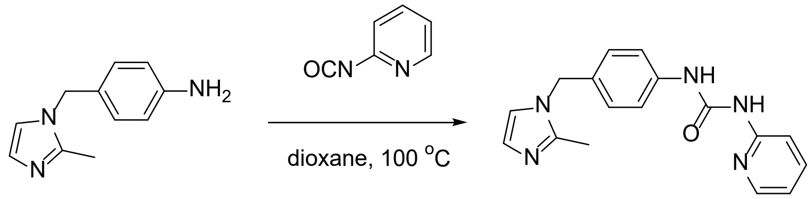


4-((2-Methyl-1H-imidazol-1-yl)methyl)aniline (200 mg, 1.07 mmol, 1.0 equiv) was added to a stirred solution of 2-isocyanatopyridine (154 mg, 1.28 mmol, 1.2 equiv) in 1,4-dioxane (3 mL) under one atmosphere of nitrogen. The resulting mixture was stirred at 100 °C for 5 hours. The solvent was removed under reduced pressure and the crude product was purified by preparative HPLC (Column: XSelect CSH Fluoro Phenyl, 30 × 150 mm, 5 µm; Mobile Phase A: water (0.1% FA), Mobile Phase B: MeCN; Flow rate: 60 mL/min; Gradient: 2% B to 12% B in 6 min, 12% B; Detector: UV 254 / 220 nm; RT: 5.85 min) to afford 1-(4-((2-methyl-1H-imidazol-1-yl)methyl)phenyl)-3-(pyridin-2-yl)urea **10** (0.28 g, 71.2%) as a formic acid salt and white solid. MS (ES$^+$, *m/z*): 308.0 [M + H]$^+$; 1H NMR (300 MHz, DMSO-*d6*) δ 2.25 (s, 3H), 5.07 (s, 2H), 6.79 (s, 1H), 6.99 (t, 1H), 7.07 – 7.16 (m, 3H), 7.43 – 7.54 (m, 3H), 7.73 (t, 1H), 8.13 (s, 1H), 8.20 – 8.35 (m, 1H), 9.41 (s, 1H), 10.49 (s, 1H).

*(S)-4-(1-(2-methyl-1H-imidazol-1-yl)ethyl)aniline (**11**)*

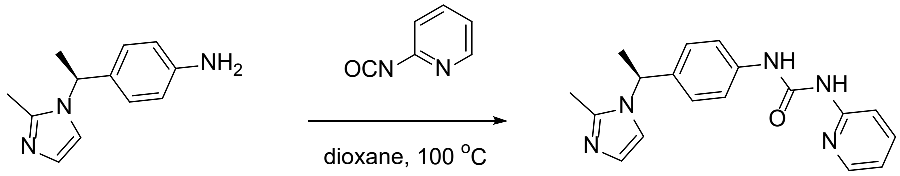


It was prepared following the same procedure described for **10** and isolated as a white solid after preparative HPLC purification (86 mg, 35.9%). MS (ES$^+$, *m/z*): 322.05 [M + H]$^+$; $^1$H NMR (300 MHz, DMSO-*d6*) 1.72 (d, J = 7.0 Hz, 3H), 2.23 (d, J = 1.7 Hz, 3H), 5.43 (d, J = 7.1 Hz, 1H), 6.81 (s, 1H), 7.01 (dd, J = 5.2, 7.4 Hz, 1H), 7.15 (d, J = 8.1 Hz, 2H), 7.28 (s, 1H), 7.49 (dd, J = 3.2, 8.7 Hz, 3H), 7.75 (td, J = 1.9, 7.1, 7.8 Hz, 1H), 8.23 – 8.31 (m, 1H), 9.42 (s, 1H), 10.47 (s, 1H).

*(R)-4-(1-(2-methyl-1H-imidazol-1-yl)ethyl)aniline (**12**)*

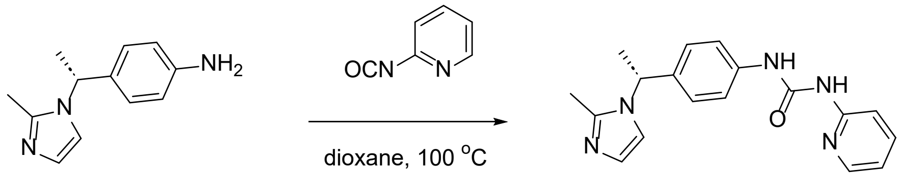


It was prepared following the same procedure described for **10** and isolated as a white solid after preparative HPLC purification (24 mg, 28.1%). MS (ES$^+$, *m/z*): 322.05 [M + H]$^+$; $^1$H NMR (300 MHz, DMSO-*d6*) δ 1.72 (d, J = 7.0 Hz, 3H), 2.23 (d, J = 1.7 Hz, 3H), 5.43 (d, J = 7.1 Hz, 1H), 6.81 (s, 1H),

7.01 (dd, J = 5.2, 7.4 Hz, 1H), 7.15 (d, J = 8.1 Hz, 2H), 7.28 (s, 1H), 7.49 (dd, J = 3.2, 8.7 Hz, 3H), 7.75 (td, J = 1.9, 7.1, 7.8 Hz, 1H), 8.23 – 8.31 (m, 1H), 9.42 (s, 1H), 10.47 (s, 1H).

#### (S)-1-(4-(1-(2-Methyl-1H-imidazol-1-yl)ethyl)phenyl)-3-(1H-pyrazol-3-yl)urea (**13**)

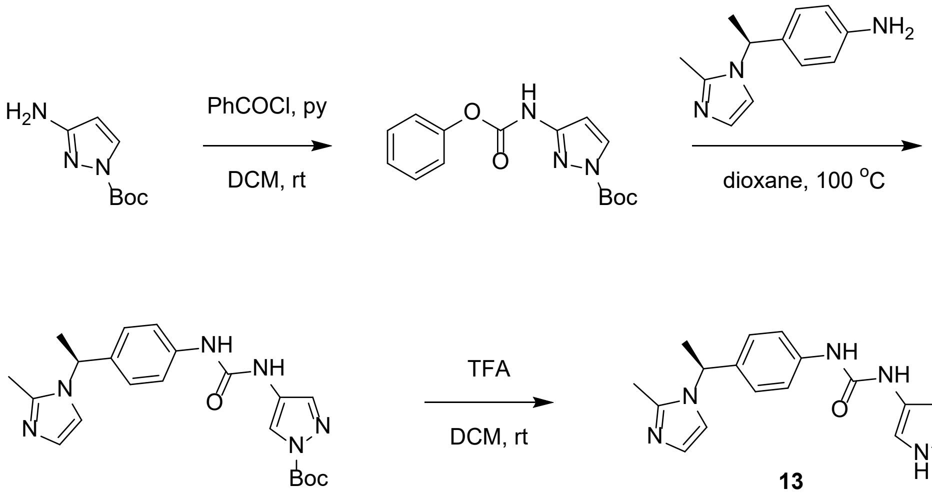


Step 1. Pyridine (2.59 g, 32.75 mmol, 3.0 equiv) was added to a mixture of tert-butyl 3-amino-1H-pyrazole-1-carboxylate (2 g, 10.92 mmol, 1.0 equiv) and phenyl carbonochloridate (2.05 g, 13.10 mmol, 1.2 equiv) in DCM (20 mL). The resulting mixture was stirred at room temperature for 3 hours, then it was quenched with water and extracted with EtOAc (3 x 25 mL). The combined organic layer was dried over $Na_2SO_4$, filtered and concentrated under reduced pressure to afford tert-butyl 3-((phenoxycarbonyl)amino)-1H-pyrazole-1-carboxylate (3.0 g, 91%) as a yellow solid. MS ($ES^+$, *m/z*): 304.0 $[M + H]^+$; $^1H$ NMR (300 MHz, DMSO-*d6*) δ 11.12 (s, 1H), 8.56 (s, 1H), 8.17 (d, J = 2.9 Hz, 1H), 7.79 – 7.07 (m, 7H), 6.81 – 6.61 (m, 1H), 1.55 (s, 9H).

Step 2. tert-Butyl 3-((phenoxycarbonyl)amino)-1H-pyrazole-1-carboxylate (332 mg, 1.09 mmol, 1.5 equiv) was added to (S)-4-(1-(2-methyl-1H-imidazol-1-yl)ethyl)aniline dihydrochloride (200 mg, 0.73 mmol, 1.0 equiv) and TEA (0.407 mL, 2.92 mmol, 4.0 equiv) in 1,4-dioxane (2 mL) under nitrogen. The resulting mixture was stirred at 70 °C for 2 hours, then it was poured into water (10 mL), extracted with EtOAc (3 x 5 mL) and washed with water (2 x 10 mL) and brine (2 x 10 mL). The organic layer was dried over $Na_2SO_4$, filtered and concentrated under reduced pressure to afford (S)-tert-butyl 3-(3-(4-(1-(2-methyl-1H-imidazol-1-yl)ethyl)phenyl)ureido)-1H-pyrazole-1-carboxylate (290 mg, 97%) as a yellow oil, which was used without further purification. MS ($ES^+$, *m/z*): 411.0 $[M + H]^+$; $^1H$ NMR (300 MHz, $CDCl_3$) δ 1.55 (s, 9H), 1.69 (d, J = 7.0 Hz, 3H), 2.23 (s, 3H), 5.18 (q, J = 7.0 Hz, 1H), 6.52 (s, 1H), 6.75 – 6.77 (m, 2H), 6.92 (d, J = 1.5 Hz, 1H), 7.07 – 7.15 (m, 2H), 7.34 – 7.45 (m, 2H), 7.83 (d, J = 3.0 Hz, 1H).

Step 3. (S)-tert-Butyl 3-(3-(4-(1-(2-methyl-1H-imidazol-1-yl)ethyl)phenyl)ureido)-1H-pyrazole-1-carboxylate (430 mg, 1.05 mmol, 1.0 equiv) was stirred in 4 M HCl in EtOH (5 mL, 20.0 mmol, 20.0 equiv) under one atmosphere of nitrogen for 1 hour. The solvent was removed under reduced pressure and the crude product was purified by preparative HPLC (Column: XBridge Shield RP18 OBD, 30 × 150 mm, 5 µm; Mobile Phase A: water (10 mM $NH_4HCO_3$ + 0.1% $NH_4OH$), Mobile Phase B: MeCN; Flow rate: 60 mL/min; Gradient: 0 to 40% B in 10 min; Detector: UV 254 / 220 nm; RT: 9.2 min) to afford (S)-1-(4-(1-(2-methyl-1H-imidazol-1-yl)ethyl)phenyl)-3-(1H-pyrazol-3-yl)urea **13** (121 mg, 37.2%) as a white solid. MS (ES$^+$, *m/z*): 311.15 $[M + H]^+$; 1H NMR (300 MHz, DMSO-*d6*) δ 1.71 (d, J = 7.0 Hz, 3H), 2.23 (d, J = 1.1 Hz, 3H), 5.40 (q, J = 7.1 Hz, 1H), 6.23 (s, 1H), 6.79 (s, 1H), 7.11 (d, J = 8.4 Hz, 2H), 7.25 (d, J = 1.6 Hz, 1H), 7.41 (d, J = 8.5 Hz, 2H), 7.58 (s, 1H), 8.93 (s, 1H), 9.07 (s, 1H), 12.23 (s, 1H).

*(S)-1-(4-(1-(2-Methyl-1H-imidazol-1-yl)ethyl)phenyl)-3-(pyridin-2-ylmethyl)urea (**14**)*

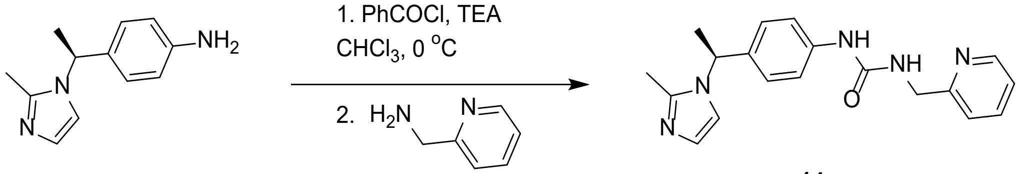


**14**

A solution of phenyl carbonochloridate (103 mg, 0.66 mmol, 1,1 equiv) in DCM (0.5 mL) was added to a stirred mixture of (S)-4-(1-(2-methyl-1H-imidazol-1-yl)ethyl)aniline dihydrochloride (150 mg, 0.55 mmol, 1.0 equiv) and TEA (166 mg, 1.64 mmol, 3.0 equiv) in $CHCl_3$ (2 mL) at 0 °C under one atmosphere of nitrogen. The reaction mixture was stirred for 1 hour at 0 °C, then pyridin-2-ylmethanamine (118 mg, 1.09 mmol, 2.0 equiv) and TEA (0.229 mL, 1.64 mmol, 3.0 equiv) were added. Stirring continued for 16 hours at room temperature. The solvent was removed under reduced pressure. The crude product was purified by preparative HPLC (Column: XBridge Prep Shield RP18 OBD, 19 × 250 mm, 5 µm; Mobile Phase A: water (10 mM $NH_4HCO_3$), Mobile Phase B: MeCN; Flow rate: 25 mL/min; Gradient: 5% B to 5% B in 2 min, 5% B to 30% B in 2.5 min, 30% B to 40% B in 10 min; Detector: UV 254 / 220 nm; RT: 6.8 min) to afford (S)-1-(4-(1-(2-methyl-1H-imidazol-1-yl)ethyl)phenyl)-3-(pyridin-2-ylmethyl)urea **14** (120 mg, 65.4 %) as a white solid. MS (ES$^+$, *m/z*): 336.20 $[M + H]^+$; 1H NMR (300 MHz, DMSO-*d6*) δ 1.69 (d, J = 7.0 Hz, 3H), 2.20 (s, 3H), 4.39 (d, J = 5.7 Hz, 2H), 5.36 (q, J = 7.0 Hz, 1H), 6.70 – 6.80 (m, 2H), 7.01 – 7.11 (m, 2H), 7.18 – 7.42 (m, 5H), 7.71 – 7.83 (m, 1H), 8.48 – 8.56 (m, 1H), 8.81 (s, 1H).

**Synthesis of amides 15 & 16**

TCFH (0.30 mmol) was added to a stirred mixture of the appropriate acid (0.30 mmol), 1-methylimidazole (1.24 mmol) and (S)-4-(1-(2-methyl-1H-imidazol-1-yl)ethyl)aniline (0.25 mmol) in MeCN (2 mL). The mixture was stirred at room temperature for 15 hours. The crude product was purified by preparative (chiral) HPLC.

Compound **15**: (S)-N-(4-(1-(2-methyl-1H-imidazol-1-yl)ethyl)phenyl)-2-(pyridin-3-yl)acetamide

MS (ES$^+$, *m/z*): 321.16 $[M + H]^+$; $^1$H NMR (300 MHz, DMSO-$d_6$) δ 1.68 (d, *J* = 7.0 Hz, 3H), 2.18 (s, 3H), 3.67 (s, 2H), 5.38 (q, *J* = 7.0 Hz, 1H), 6.75 (d, *J* = 1.3 Hz, 1H), 7.11 (d, 2H), 7.21 (d, *J* = 1.4 Hz, 1H), 7.30 – 7.39 (m, 1H), 7.53 (d, 2H), 7.71 (dt, *J* = 7.9, 2.0 Hz, 1H), 8.41 – 8.54 (m, 2H), 10.26 (s, 1H).

Compound **16**: 2,2-difluoro-N-(4-((S)-1-(2-methyl-1H-imidazol-1-yl)ethyl)phenyl)-2-((R*)-tetrahydrofuran-3-yl)acetamide

MS (ES$^+$, *m/z*): 350.16 [M + H]$^{+;}$ $^1$H NMR (400 MHz, DMSO-$d_6$) δ 1.72 (d, *J* = 7.0 Hz, 3H), 1.85 – 1.97 (m, 1H), 1.97 – 2.10 (m, 1H), 2.21 (s, 3H), 3.06 – 3.23 (m, 1H), 3.64 (q, *J* = 7.6 Hz, 1H), 3.78 (dd, *J* = 6.1, 11.8 Hz, 3H), 5.44 (q, *J* = 7.0 Hz, 1H), 6.79 (d, *J* = 1.4 Hz, 1H), 7.15 – 7.22 (m, 2H), 7.27 (d, *J* = 1.4 Hz, 1H), 7.61 – 7.67 (m, 2H), 10.65 (s, 1H).

**Synthesis of adduct 6**

*Sodium ((2R,3S,4R,5R)-5-(6-amino-9H-purin-9-yl)-3,4-dihydroxytetrahydrofuran-2-yl)methyl (1H-imidazol-1-yl)phosphonate*

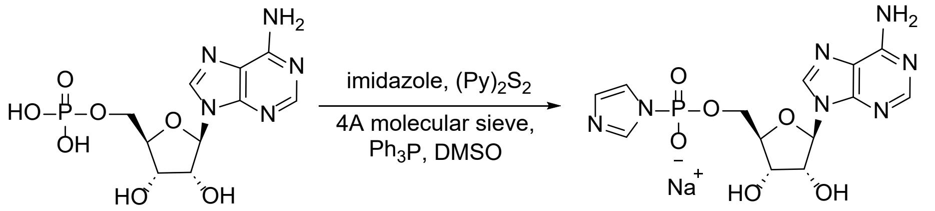


A suspension of ((2*R*,3*S*,4*R*,5*R*)-5-(6-amino-9*H*-purin-9-yl)-3,4-dihydroxytetrahydrofuran-2-yl)methyl dihydrogen phosphate (600 mg, 1.72 mmol, 1.0 equiv), imidazole (1.18 g, 17.28 mmol, 10.0 equiv), $PPh_3$ (1.36 g, 5.18 mmol, 3.0 equiv) and 4 A molecular sieves (100 mg, activated) in anhydrous DMSO (6 mL) was stirred for 1 hour at room temperature. A solution of 2,2-dipyridyldisulfide (1.14 g, 5.18 mmol, 3.0 equiv) in anhydrous DMSO (3 mL) was added dropwise over 15 minutes. The resulting mixture was stirred for 3 hours at room temperature, then slowly added to a solution of NaI in acetone (0.1 M, 90 mL). The precipitate was collected by filtration and dried under reduced pressure to afford sodium ((2*R*,3*S*,4*R*,5*R*)-5-(6-amino-9*H*-purin-9-yl)-3,4-dihydroxytetrahydrofuran-2-yl)methyl (1*H*-imidazol-1-yl)phosphonate (700 mg, 86.9%) as a light-yellow semi-solid. MS (ES$^+$, *m/z*): 398.15 [M + H]$^+$. $^1$H NMR (400 MHz, DMSO-*d6*) δ 8.40 (s, 1H), 8.14 (d, *J* = 4.0 Hz, 1H), 7.72 (d, *J* = 8.0 Hz, 1H), 7.29 (s, 2H), 7.12 (d, *J* = 4.0 Hz, 1H), 6.90 (s, 1H), 5.93 – 5.88 (m, 1H), 5.46 (s, 1H), 5.28 (s, 1H), 4.58 (s, 1H), 4.03 (s, 1H), 3.94 – 3.80 (m, 1H), 3.80 – 3.71 (m, 2H). $^{31}$P NMR (162 MHz, DMSO-*d6*) δ -10.39.

*(2R,3R,4R)-2-(acetoxymethyl)-5-chlorotetrahydrofuran-3,4-diyl diacetate*

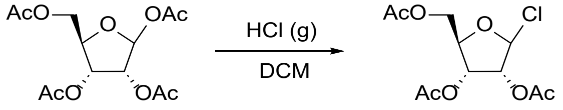


Anhydrous HCl gas was bubbled into a stirred mixture of (3*R*,4*R*,5*R*)-5-(acetoxymethyl)tetrahydrofuran-2,3,4-triyl triacetate (1.80 g, 1.0 equiv) in DCM (70 mL) at 0 °C for 1 hour. The reaction mixture was concentrated under reduced pressure. The residual solvent was removed by azeotropic distillation with toluene (3 x 10 mL) to afford (2*R*,3*R*,4*R*)-2-(acetoxymethyl)-5-chlorotetrahydrofuran-3,4-diyl diacetate (1.6 g) as a colorless oil which was used without further purification. MS (ES$^+$, *m/z*): 258.90 [M - Cl]$^+$.

*1-((2R,3R,4R,5R)-3,4-diacetoxy-5-(acetoxymethyl)tetrahydrofuran-2-yl)-4-(4-(3-(pyridin-2-yl)ureido) benzyl)pyridin-1-ium chloride*

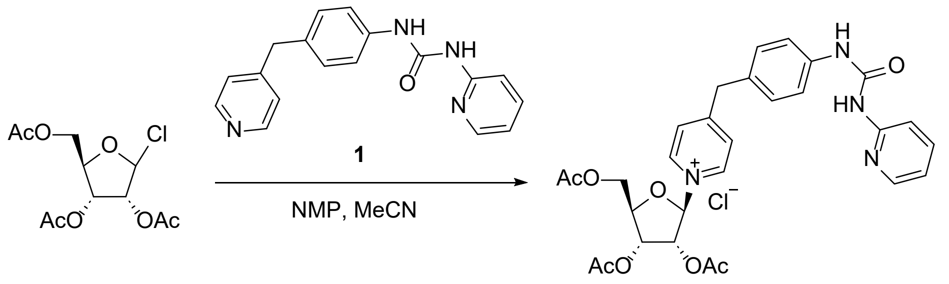


1-(Pyridin-2-yl)-3-(4-(pyridin-4-ylmethyl)phenyl)urea **1** (991.51 mg, 3.258 mmol, 0.6 equiv) was added to a stirred mixture of freshly prepared (2*R*,3*R*,4*R*)-2-(acetoxymethyl)-5-chlorotetrahydrofuran-3,4-diyl diacetate (1.6 g, 5.43 mmol, 1.0 equiv) in MeCN (70 mL) and NMP (10 mL) at room temperature. The resulting mixture was stirred 16 hours, then it was concentrated under reduced pressure and the residue was triturated with $CHCl_3$ / $Et_2O$ (1/5, 30 mL) to afford 1-((2*R*,3*R*,4*S*,5*R*)-3,4-dihydroxy-5-(hydroxymethyl)tetrahydrofuran-2-yl)-4-(4-(3-(pyridin-2-yl)ureido)benzyl)pyridin-1-ium chloride (1.6 g, 39% purity, 61% impurity **1**) as a yellow solid. MS ($ES^+$, *m/z*): 563.05 $[M - Cl]^+$.

*1-((2R,3R,4S,5R)-3,4-dihydroxy-5-(hydroxymethyl)tetrahydrofuran-2-yl)-4-(4-(3-(pyridin-2-yl)ureido) benzyl)pyridin-1-ium chloride*

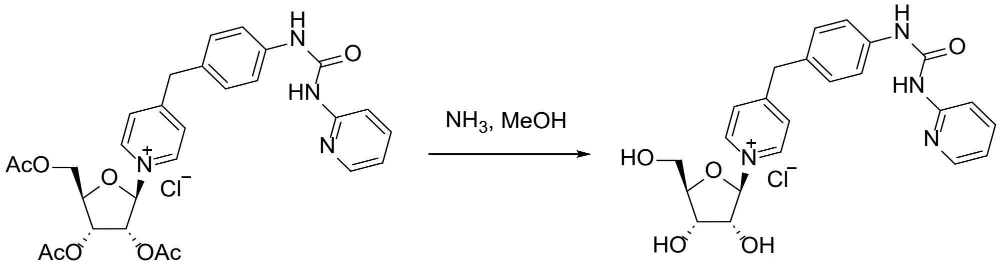


$NH_3$ (4 M in MeOH, 0.62 mL, 4.34 mmol, 4.0 equiv) was added dropwise into a stirred solution of 1-((2*R*,3*R*,4*S*,5*R*)-3,4-dihydroxy-5-(hydroxymethyl)tetrahydrofuran-2-yl)-4-(4-(3-(pyridin-2-yl)ureido)benzyl)pyridin-1-ium chloride (650 mg, 1.085 mmol, 1.0 equiv) in MeOH (13 mL) at -10 °C under one atmosphere of argon. The resulting mixture was stirred at -10 °C for 16 hours, then concentrated under reduced pressure. The crude product was purified by preparative HPLC (Column: Atlantis Prep T3 OBD, 19 x 250 mm, 10 μm; Mobile Phase A: water (0.05% TFA), Mobile Phase B: MeCN; Flow rate: 20 mL / min; Gradient: 20% B to 45% B in 6.5 min, Detector: UV 210 / 254 nm; RT: 5.3 min) to afford 1-((2*R*,3*R*,4*S*,5*R*)-3,4-dihydroxy-5-(hydroxymethyl)tetrahydrofuran-2-yl)-4-(4-(3-(pyridin-2-yl)ureido)benzyl)pyridin-1-ium chloride (100 mg, 19.5%) as a light yellow solid. MS ($ES^+$, *m/z*): 437.10 $[M - Cl]^+$; $^1H$ NMR (400 MHz, $D_2O$) δ 8.84 (d, $J$ = 4.0 Hz, 2H), 8.19 – 8.14 (m, 2H), 7.86 (s, 2H), 7.42 – 7.22 (m, 6H), 5.99 (s, 1H), 4.33 (s, 2H), 4.26 – 4.22 (m, 3H), 3.88 (s, 1H), 3.79 – 3.76 (m, 1H).

*Ammonium ((2R,3S,4R,5R)-3,4-dihydroxy-5-(4-(4-(3-(pyridin-2-yl)ureido) benzyl)pyridin-1-ium-1-yl)tetrahydrofuran-2-yl)methyl phosphate chloride*

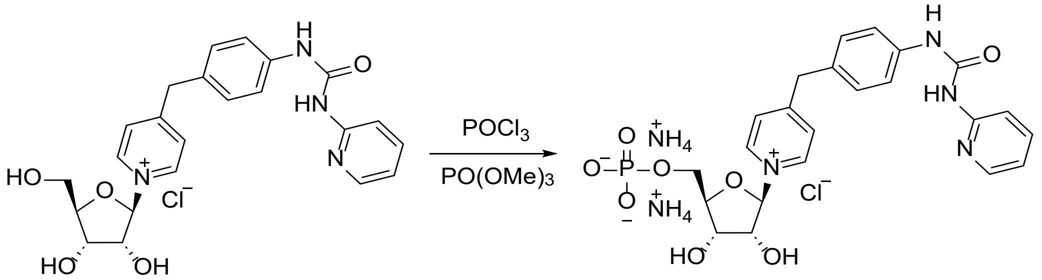


A solution of $POCl_3$ (259.37 mg, 1.69 mmol, 10.0 equiv) in trimethyl phosphate (0.5 mL) was added dropwise into a stirred mixture of 1-((2*R*,3*R*,4*S*,5*R*)-3,4-dihydroxy-5-(hydroxymethyl)tetrahydrofuran-2-yl)-4-(4-(3-(pyridin-2-yl)ureido)benzyl)pyridin-1-ium chloride (80 mg, 0.169 mmol, 1.0 equiv) in trimethyl phosphate (2 mL) at 0 °C under one atmosphere of argon. The reaction mixture was stirred at -5 °C for 5 hours, then it was quenched with water (4 mL) at 0 °C, extracted with $Et_2O$ (2 x 4 mL) and $CHCl_3$ (2 x 4 mL). The aqueous layer was concentrated under reduced pressure and the crude product was purified by preparative HPLC (Column: Atlantis Prep T3 OBD, 19 x 250 mm 10 µm; Mobile Phase A: water (10 mM $NH_4HCO_3$), Mobile Phase B: MeCN; Flow rate: 18 mL / min; Gradient: 5% B to 25% B in 7 min, Detector: UV 254 / 210 nm; RT: 5.2 min) to afford ammonium ((2*R*,3*S*,4*R*,5*R*)-3,4-dihydroxy-5-(4-(4-(3-(pyridin-2-yl)ureido)benzyl) pyridin-1-ium-1-yl)tetrahydrofuran-2-yl)methyl phosphate chloride (20 mg, 20.14%) as a white solid. MS ($ES^+$, *m/z*): 517.10 $[M - Cl - 2\ NH_3]^+$; $^1H$ NMR (300 MHz, $D_2O$) δ 8.90 – 8.88 (m, 2H), 8.32 – 8.19 (m, 2H), 7.92 (d, *J* = 6.0 Hz, 2H), 7.45 – 7.36 (m, 4H), 7.33 – 7.30 (m, 2H), 6.03 - 6.02 m, 1H), 4.53 – 4.51 (m 1H), 4.43 – 4.36 (m, 2H), 4.31 (s, 2H), 4.25 – 4.18 (m, 1H), 4.12 – 4.10 (m, 1H). $^{31}P$ NMR (121 MHz, $D_2O$) δ -0.09.

*Diammonium 1-[(2R,3R,4S,5R)-5-[[([[(2R,3S,4R,5R)-5-(6-aminopurin-9-yl)- 3,4-dihydroxyoxolan-2-yl]methyl phosphono]oxy(oxido)phosphoryl)oxy]methyl]-3,4-dihydroxyoxolan-2-yl]-4-[(4-[[(pyridin-2-yl)carbamoyl]amino]phenyl)methyl]-1λ⁵-pyridin-1-ylium chloride (**6**)*

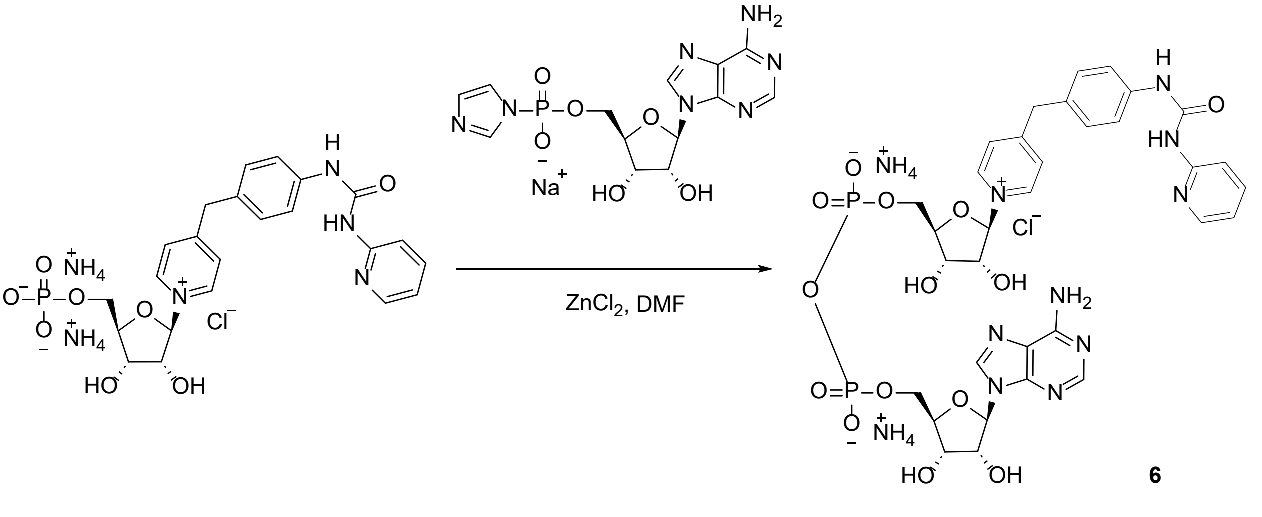


**6**

A mixture of anhydrous $ZnCl_2$ (53.41 mg, 0.392 mmol, 10.0 equiv) and ammonium ((2*R*,3*S*,4*R*,5*R*)-3,4-dihydroxy-5-(4-(4-(3-(pyridin-2-yl)ureido)benzyl) pyridin-1-ium-1-yl)tetrahydrofuran-2-

yl)methyl phosphate chloride (23 mg, 0.039 mmol, 1.0 equiv) in DMF (1 mL) was stired at 40 °C until it became a clear solution. A solution of sodium ((2R,3S,4R,5R)-5-(6-amino-9H-purin-9-yl)-3,4-dihydroxytetrahydrofuran-2-yl)methyl (1H-imidazol-1-yl)phosphonate (32.8 mg, 0.032 mmol, 2.0 equiv) in DMF (1.5 mL) dropwise over 5 minutes at 30 °C. The resulting solution was stirred at 25 °C for 15 hours, then it was slowly poured into 1 M aqueous $NH_4HCO_3$ (2.0 mL) at 0 °C with stirring. The solid was filtered out, the filtrate was purified by preparative HPLC (Column: Xbridge Phenyl OBD, 19 × 150 mm 5 μm; Mobile Phase A: water (10 mM $NH_4HCO_3$), Mobile Phase B: MeCN; Flow rate: 20 mL / min; Gradient: 3% B to 20% B in 6 min, Detector: UV 210 / 254 nm; RT: 5.2 min) to afford diammonium 1-[(2*R*,3*R*,4*S*,5*R*)-5-[[(([[(2*R*,3*S*,4*R*,5*R*)-5-(6-aminopurin-9-yl)-3,4-dihydroxyoxolan-2-yl]methyl phosphono]oxy(oxido)phosphoryl)oxy]methyl]-3,4-dihydroxyoxolan-2-yl]-4-[(4-[[(pyridin-2-yl)carbamoyl]amino]phenyl)methyl]-1$\lambda^5$-pyridin-1-ylium chloride **6** (11 mg, 30.64% yield) as a light yellow solid. MS ($ES^+$, *m/z*): 846.10 $[M - Cl - 2\ NH_3]^+$; $^1$H NMR (400 MHz, $D_2O$) δ 8.72 – 8.70 (m, 2H), 8.16 (s, 1H), 8.08 – 8.07 (m, 1H), 7.91 (d, *J* = 4.0 Hz, 1H), 7.75 – 7.73 (m, 2H), 7.65 – 7.58 (m, 1H), 7.26 – 7.21 (m, 2H), 7.11 – 7.05 (m, 2H), 7.02 – 6.89 (m, 2H), 5.87 – 5.80 (m, 2H), 4.56 – 4.53 (m, 1H), 4.44 – 4.40 (m, 1H), 4.34 – 4.27 (m, 5H), 4.15 – 4.11 (m, 2H), 4.04 – 4.02 (m, 3H). $^{31}$P NMR (162 MHz, $D_2O$) δ -11.362, -11.407.

**Biocatalytic Synthesis of Adduct 6**

*Immobilisation of SARM1$^{\Delta 1\text{-}27}$*

A SARM1$^{\Delta 1\text{-}27}$ stock solution (1.2 mL, 1.68 mg/mL) was diluted with reaction buffer (50 mM HEPES and 150 mM NaCl at pH 7.5, 2.8 mL) and $Ni^{2+}$ resin (Purolite Ltd. Chromalite MIDA/M/$Ni^{2+}$ resin, 200 mg) was added. The mixture was incubated at room temperature on a rotary mixer for 20 h and the protein concentration in the supernatant was determined using a NanoDrop spectrophotometer to calculate the amount of immobilized SARM1$^{\Delta 1\text{-}27}$ (1.86 mg on 200 mg resin). The mixture was centrifuged at 2000 rpm for 1 min, the supernatant was removed and the resin was washed with reaction buffer (3 x 3 mL). The resin containing immobilized SARM1$^{\Delta 1\text{-}27}$ was either used immediately or stored in reaction buffer at 5 °C for subsequent reactions.

*Enzymatic Reaction*

To a Falcon tube containing immobilized SARM1$^{\Delta 1\text{-}27}$, stock solutions of $NAD^+$ (20 mM, 2.4 mL, 48 μmol, 20 equiv) and NMN (20 mM, 2.4 mL, 48 μmol, 20 equiv) in reaction buffer, and **1** (25 mM, 96 μL, 2.4 μmol, 1 equiv) in DMSO were sequentially added and the reaction mixture was incubated at 35 °C and agitated at 500 rpm in a thermocycler for 16–20 h. The reaction mixture was centrifuged (2000 rpm, 1 min.), the supernatant was collected, and the resin was washed with washing buffer (reaction buffer containing 5 vol% DMSO, 2 x 3 mL), while shaking at 400 rpm on a thermocycler for 30–60 minutes. Supernatant and washing solutions were collected. The synthesis cycle was repeated 5 times to ensure the production of sufficient adduct for purification. For storage, the resin was kept in reaction buffer at 4°C until further use.

*Samples preparation and purification*

Sample preparation was performed on a Waters OASIS HLB SPE cartridge in accordance with the manufacturer's recommended protocol. In brief, a Waters OASIS HLB SPE cartridge (6 cc, 200 mg) was conditioned with MeOH (4 mL) and equilibrated with Milli-Q water (2 x 4 mL) using slight overpressure supplied by a stream of nitrogen. The supernatant from each synthesis cycle was introduced to the column, and elution was similarly carried out with gentle nitrogen pressure. The column was washed with Milli-Q water (2 x 4 mL), and **6** was eluted using a 1:1 mixture of MeCN and water (2 x 4 mL). The eluent was concentrated and purified by preparative HPLC (Column: XBridge, 10 × 100 mm, 5 µm; Mobile phase A: 97% water, 3% ACN and 0.8 g/L $NH_4HCO_3$; Mobile phase B: MeCN; Flow rate: 8 mL / min; Gradient: 0% B to 30% B in 6 min; Detector 210 / 254 nm; RT: 3.8 min ) to afforded **6** (0.2 mg) in 96% purity. The analytical data are in alignment with the data reported for the chemical synthesis route of **6**.

$^1$H NMR of adduct **6**

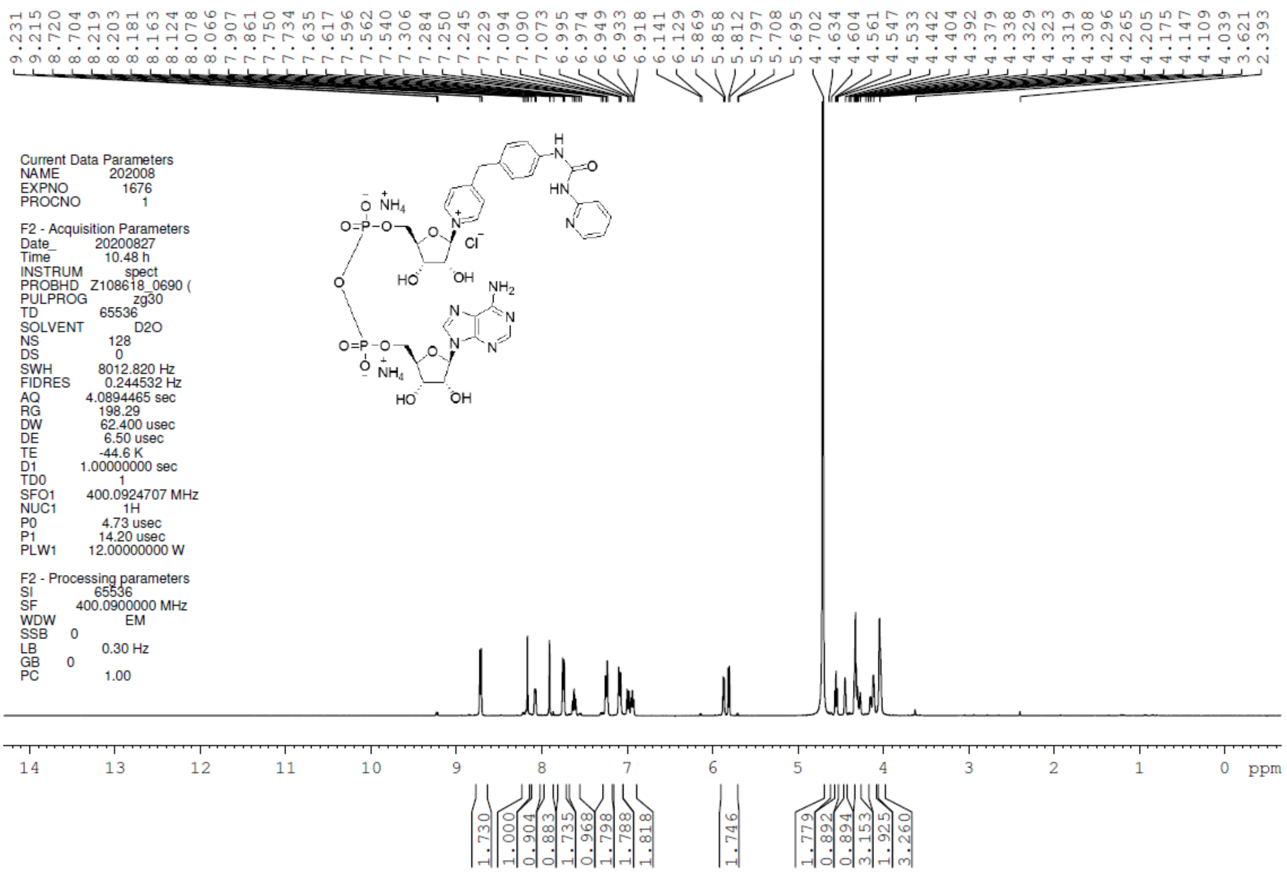

[1]H NMR of urea **14**

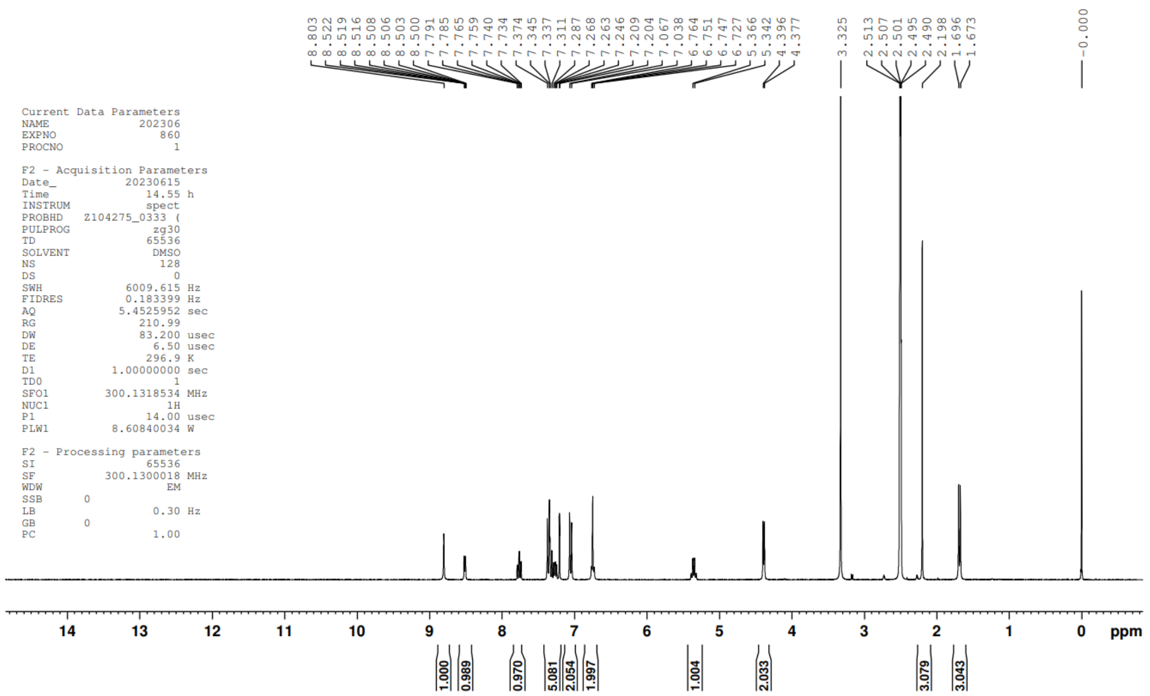

[1]H NMR of amide **16**

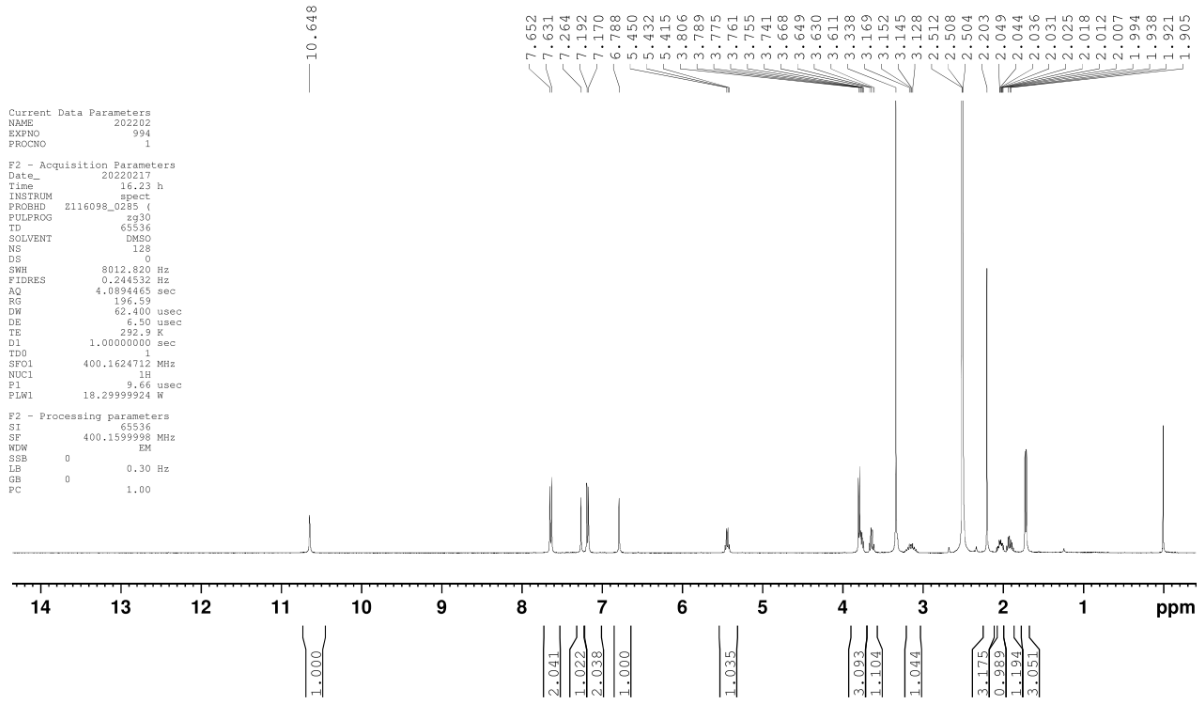
